# Systematic Mapping of Bacterial CRISPRa Design Rules and Implications for Synergistic Gene Activation

**DOI:** 10.1101/2025.05.14.654134

**Authors:** Cholpisit Kiattisewee, Ava V. Karanjia, Ryan A. L. Cardiff, Kira E. Olander, Pansa Leejareon, Sarah S. Alvi, James M. Carothers, Jesse G. Zalatan

## Abstract

CRISPR gene activation (CRISPRa) tools have shown great promise for bacterial strain engineering but often require customization for each intended application. Our goal is to create generalizable CRISPRa tools that can overcome previous limitations of gene activation in bacteria. In eukaryotic cells, multiple activators can be combined for synergistic gene activation. To identify potential effectors for synergistic activation in bacteria, we systematically characterized bacterial activator proteins with a set of engineered synthetic promoters. We found that optimal target sites for different activators could vary by up to 200 bases in the region upstream of the transcription start site (TSS). These optimal target sites qualitatively matched previous reports for each activator, but the precise targeting rules varied between different promoters. By characterizing targeting rules in the same promoter context, we were able to test activator combinations with each effector positioned at its optimal target site. We did not find any activator combinations that produced synergistic activation, and we found that many combinations were antagonistic. This systematic investigation highlights fundamental mechanistic differences between bacterial and eukaryotic transcriptional activation systems, and suggests that alternative strategies will be necessary for strong bacterial gene activation at arbitrary endogenous targets.

## Introduction

CRISPR-Cas based transcriptional regulators have become a pivotal tool in genetic programming. CRISPR activation (CRISPRa) and CRISPR interference (CRISPRi) enable massively parallel perturbations at a genome-wide scale.^1–3^ In bacterial systems, CRISPRa systems that recruit transcriptional activators to target genes have been used for combinatorial pathway engineering,^4^ activating endogenous targets,^5–7^ multi-layer dynamic circuits,^8–10^ and metabolic engineering in non-model organisms.^11,12^ Although past work has estimated that approximately 60% of *E. coli* promoters are targetable with CRISPRa,^7^ challenges such as inconsistent activator performance across bacterial promoters, strict design constraints specific to each activator, and interference from native regulatory machinery at endogenous gene targets continue to hinder broader implementation of CRISPRa tools.^6,7,13–15^

To expand our capabilities in bacterial systems, we can look to eukaryotic CRISPRa systems,^16–19^ where several groups have demonstrated that delivering multiple effectors can produce potent and synergistic transcriptional activation.^20–22^ In both eukaryotic and bacterial systems, synergistic transcriptional effects have been well-established prior to the development of CRISPR technologies.^23–25^ Typically, these effects arise from simultaneous recruitment of multiple transcription factors, which can produce expression levels greater than the summed effect of each individual activator. Synergistic CRISPRa in bacteria has yet to be achieved, possibly due to the stringent target site and sequence requirements that govern activation.^6,13– 15,26–30^ There has been limited progress in engineered bacterial synergistic activators in the ∼30 years since the first report of synergistic activation in bacteria, and new approaches are needed to identify bacterial gene activation methods that rival the robust programmability and effectiveness of eukaryotic CRISPRa systems.^31–35^

To determine if improved and potentially synergistic CRISPRa can be achieved, we evaluated gene activation in *E. coli* with combinations of CRISPRa effectors. To identify candidate combinations, we first performed a systematic evaluation of multiple bacterial CRISPRa systems, including several with distinct activation mechanisms.^6,13,14^ After identifying optimal target sites for each individual CRISPRa system, we defined strategies to recruit combinations of CRISPRa effectors at their respective optimal target sites. Design rules for these systems have been previously reported,^6,13,14^ but we found that it was critical to characterize candidate systems in the same promoter context because the optimal target sites can vary between different promoters.

Although we identified multiple scenarios where combinations of effectors could be recruited to their respective optimal target sites at a single promoter, we did not observe any significant synergistic CRISPRa effects. Instead, when two effectors were recruited to a single CRISPR-Cas complex, we observed antagonistic effects on activation. When two effectors were recruited to distinct optimal target positions with orthogonal CRISPR-Cas systems, we observed no increase in activation relative to the best single effector. Bacterial effectors typically interact directly with RNA polymerase (RNAP), and our results may indicate that RNAP cannot bind to the minimal promoter and simultaneously interact with two DNA-bound activators, at least for the pairs of effectors tested here. These findings underscore ongoing challenges for synergistic gene activation in bacteria. Although synergistic transcriptional activation has already been achieved in eukaryotes with combinations of CRISPRa effectors,^20–22^ our results suggest that alternative strategies may be needed to obtain reliable high-level gene activation at endogenous genes in bacteria.

## Materials and Methods

### Bacterial strains and plasmid constructs

The bacterial strains used in this study are listed in Table S1. All PCR fragments were amplified with Phusion DNA Polymerase (Thermo-Fisher Scientific) for Infusion Cloning (Takara Bio). Plasmids were transformed into chemically competent *E. coli* DH10B (Thermo Fisher Scientific) or NEB turbo (New England Biolabs) cells, plated on LB-agar, and cultured in LB media with the appropriate antibiotics used in the following concentrations: 100 μg/mL Carbenicillin, 25 μg/mL Chloramphenicol, 30 μg/mL Kanamycin. Plasmids were constructed following previously described methods.^6,27,36^ Detailed descriptions of each plasmid used in this work are provided in Table S2. Different CRISPRa plasmids harboring catalytically-inactive Cas9 (dCas9) were constructed using pCD442 (constitutive expression) or pSLQ1055 (aTc-inducible) as backbones.^6,37^ dCas9 and Spy-dCas9 refer to *Streptococcus pyogenes* dCas9. Sth1-dCas9 refers to a dCas9 homolog from *Streptococcus thermophilus*.^38^ *Lachnospiraceae bacterium* dCas12a (Lb-dCas12a or Lb-dCpf1) was used for all dCas12a experiments.^39,40^ The SoxS effector used in this study contains two mutations (R93A/S101A) as previously described.^6^ The PspF effector used in this study lacks a DNA binding domain as previously reported.^14^ AsiA_m2.1, referred to as AsiA in this study, was designed with the amino acid sequence as reported previously by Ho and coworkers.^13^ Modified guide RNA sequences were constructed as described previously.^14,27^ Throughout the manuscript, “-” will indicate direct protein fusions and “/” will indicate non-covalent recruitment. For example, dCas9-AsiA represents direct fusion of dCas9 and AsiA while dCas9/MCP-SoxS represents MCP-SoxS recruitment to dCas9 via the interaction of MCP with the MS2 RNA hairpin. All plasmid constructs were confirmed by Sanger sequencing (Genewiz/Azenta Life Sciences) or Nanopore sequencing (Primordium/Plasmidsaurus). All relevant sequences are provided in the Supporting Information DNA sequences section. Plasmid maps are deposited as Supporting Information. Selected plasmids will be available on Addgene.

### Reporter design and distance screening

To characterize the target site preferences of each CRISPRa system, promoters were designed with J1 upstream sequences (170 bp) containing NGG PAMs every 10bp on both strands.^27^ Because the average intergenic region upstream of *E. coli* genes is ∼140 bp, the 170 bp sequence covers most targetable sites of *E. coli* promoters.^41^ The previously described J1-BBa_J23117-mRFP reporter was used for the σ^70^-compatible CRISPRa systems. For AsiA, where the optimal target position is further upstream, the J1 array was moved further upstream by inserting additional sequences (100, 200, or 122 bp) between the J1 array and the BBa_J23117 minimal promoter. J1-J3(100) has 100 bp of J3 inserted between J1 and the minimal promoter. J1-J2(100)-J3(100) has 100 bases of J2 and 100 bases of J3 inserted (200 bases total). J1-J3(122) has 122 bases of J3 inserted. The J2 and J3 sequences were described previously.^6,10^ Complete sequences of J1-J3(100), J1-J2(100)-J3(100), and J1-J3(122) are provided in the Supporting Information. These designs mimic the W1 promoter used for AsiA activation.^13,26^ The J3 sequence adjacent to the minimal promoter includes a J306 target site positioned to be compatible with SoxS/αNTD/TetD CRISPRa (Figure S6 & S7). For PspF-mediated CRISPRa at σ^54^ promoters, the BBa_J23117 minimal promoter was replaced with 35 bp of the pspAp minimal promoter.^14^ gRNA target sites are listed in Table S3. Single nucleotide phase-shift promoters were designed by addition of N bp (+N) between J1 and the minimal promoter or by deletion of N bp (−N) proximal to the minimal promoter. For alternative dCas proteins with PAM preferences different from the NGG used by Spy-dCas9, new promoter systems were designed by mutating the NGG PAM sequence into a compatible PAM or using a strategy previously used for J1 sequence design by tiling PAMs (see Supporting Methods).^27^

### Gene expression measurements via fluorescent reporter

Single colonies of bacteria from LB plates were inoculated in 400 μL of EZ-RDM (Teknova) supplemented with the appropriate antibiotics and grown in 96-deep-well plates at 37 °C with shaking overnight at 900 RPM on a Heidolph titramax 1000 for 18 h. 150 μL of the overnight culture were transferred into flat, clear-bottomed black 96-well plates (Corning) and the OD_600_ and fluorescence were measured in a Biotek Synergy HTX plate reader. Data were analyzed using the BioTek Gen5 2.07.17 software. For mRFP1 detection, the excitation wavelength was 540 nm and emission wavelength was 600 nm. For constructs controlled by an aTc-inducible promoter (pCK320, pCK321, and pAK007), single colonies were first inoculated in the inducer-free EZ-RDM. After overnight incubation, the starter cultures were diluted 100-fold into EZ-RDM containing appropriate antibiotics with or without 200 nM anhydrotetracycline (aTc), unless specified. The cultures were further incubated overnight before fluorescent protein measurement.

## Results

Different bacterial CRISPRa strategies have distinct target site preferences depending on their mechanism of interaction with RNAP.^42,43^ Here, we focus on four CRISPRa effectors with different RNAP interaction mechanisms: 1) SoxS,^6,27^ 2) N-terminal domain of RpoA (αNTD),^15,27^ 3) AsiA,^13^ and 4) PspF (Figure 1).^14^ We hypothesized that we could target multiple effectors to the same promoter to increase RNAP recruitment and improve activation. For effectors with the same optimal target sites, we could recruit two effectors to the same CRISPR complex. Alternatively, for effectors with distinct optimal target sites, we could use two orthogonal CRISPR complexes to recruit effectors to their respective sites. However, it is challenging to identify optimal pairs of target sites from the available literature because many CRISPRa systems have been characterized with different model promoters, and the optimal target site position can be sensitive to promoter context.

**Figure 1.**
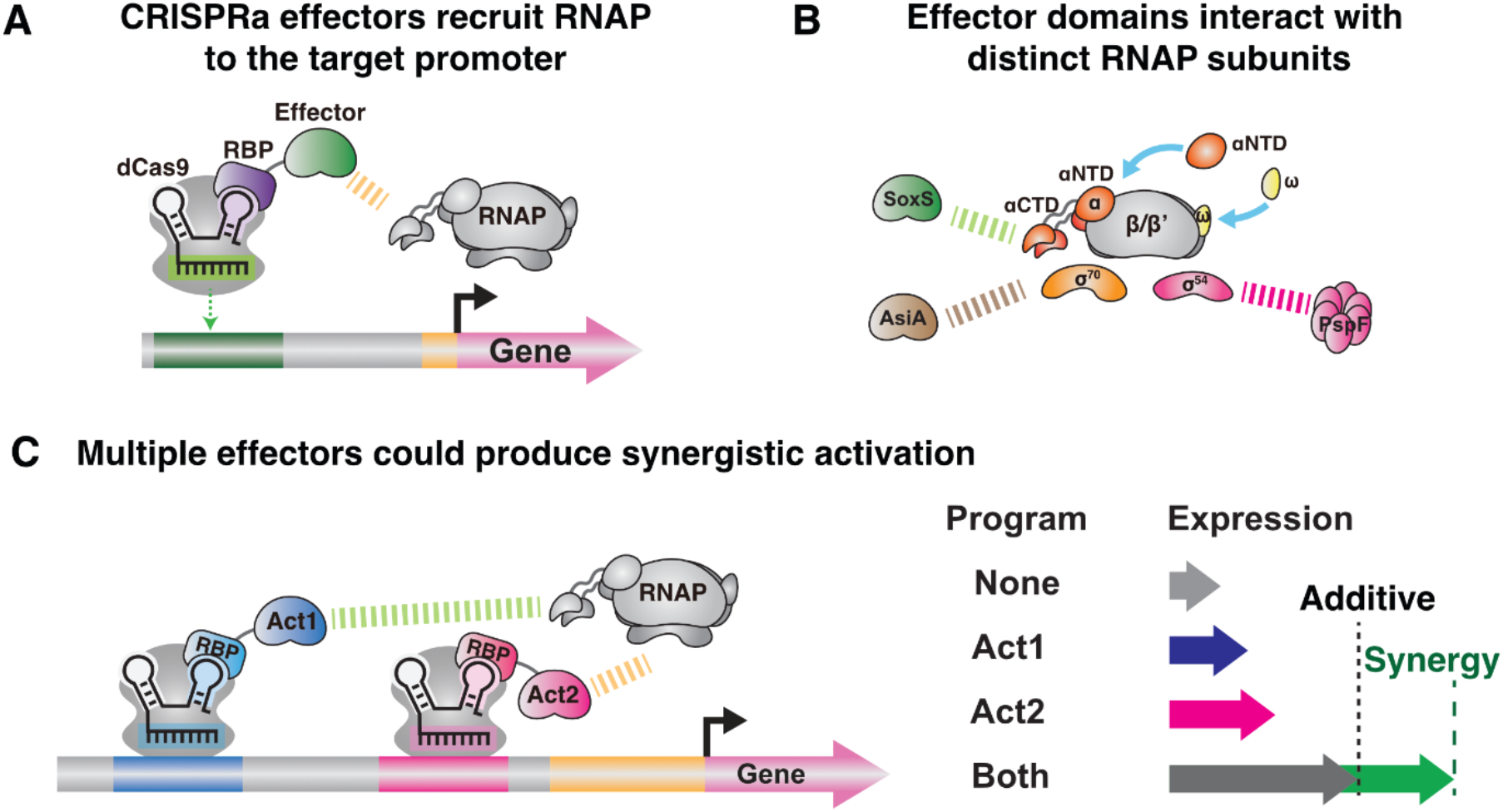
CRISPRa-recruited effector proteins activate gene expression. **A**) Bacterial CRISPRa systems recruit transcription factors that interact with RNAP. **B**) Bacterial CRISPRa effectors use various mechanisms to recruit RNAP, including interactions with the α-subunit (RpoA), the σ^70^-subunit (RpoD), or the σ^54^-subunit (RpoN). Some strategies use an RNAP subunit itself as the transcriptional effector in the CRISPR-Cas complex. **C**) Proposed model for synergistic activation with multiple CRISPRa complexes providing cooperative interactions with RNAP.

To facilitate comparisons between CRISPRa systems in a single model promoter, we modified a fluorescent reporter system previously used to characterize SoxS and αNTD.^6,27^ The J1 synthetic promoter contains a target site array that includes NGG PAMs oriented every 10 bp upstream from the TSS on both strands, starting from -35 bp from the TSS, and can be shifted by 1 to 12 base pairs by inserting single base pairs at the 5’ end of UP element to characterize target sites preferences at single base resolution. To construct a promoter system for multiple CRISPRa effectors, we modified the J1 promoter to be compatible for testing the design rules of AsiA and PspF activators. For AsiA, we inserted an additional sequence between the J1 target site array and the core promoter, which shifts the target site array further upstream of the TSS previously shown to be optimal for AsiA activation (Figure 2A).^13^ For PspF, we changed the core promoter from σ^70^ to σ^54^, which is required for PspF-mediated activation (Figure 2B). All optimized promoters contain the same J1 target site array, which should enable comparisons between effectors because the promoters have the same target sites and surrounding sequence context. We first validated each CRISPRa system in its original reporter construct and then proceeded to systematically compare these systems using J1 and modified J1 reporters.

**Figure 2.**
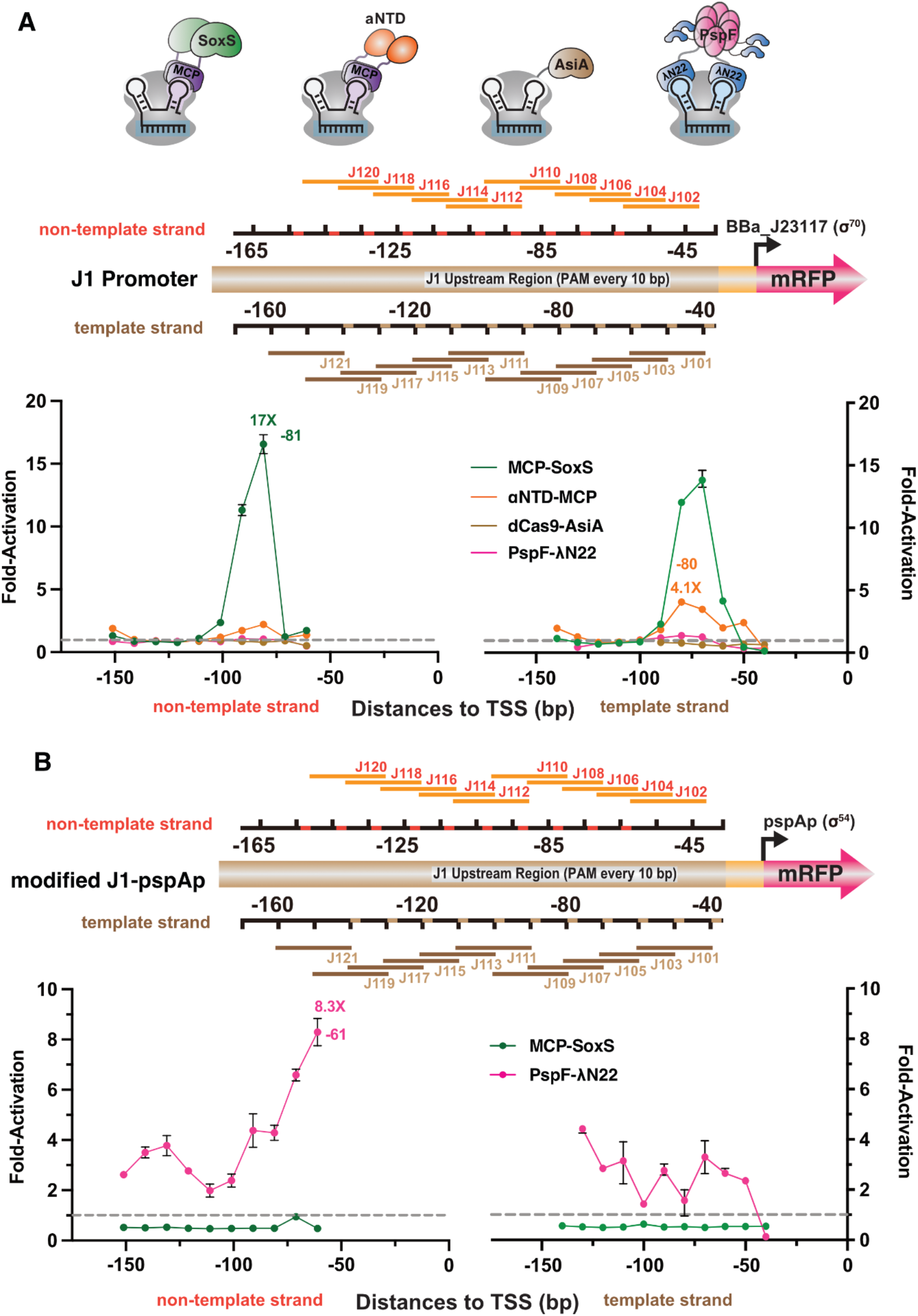
CRISPRa is highly sensitive to target site and promoter context. **A**) The J1 promoter was used to screen four CRISPRa effectors. J1 contains an array of PAM sequences every 10 bp on both strands upstream of the σ^70^ minimal promoter BBa_J23117 (DNA sequences available in Supporting Information). MCP-SoxS and αNTD-MCP showed optimal fold-activation at -81(NT) and -70(T), respectively. dCas9-AsiA and PspF-λN22 exhibited no activity at any J1 target sites. dCas9-AsiA has activity at target sites further upstream (Figure S7). **B**) Screening for activity with MCP-SoxS and PspF-λN22 using a J1 array upstream of a σ^54^ minimal promoter pspAp. Only PspF showed CRISPRa activity with maximum activity at -61(NT). Values in panel A and B represent the mean ± standard deviation calculated from n = 3.

### Design rules for SoxS family activators

The SoxS transcriptional regulator from *E. coli* is an effective CRISPR activation domain in multiple bacterial species.^12,27,36,44,45^ SoxS can be recruited as an MCP-SoxS fusion to an MS2 scaffold RNA (scRNA) and we have previously screened SoxS-mediated CRISPRa at single nucleotide resolution 40-120 bp upstream of the TSS in the J1 synthetic promoter (Figure 2A).^6^ SoxS-mediated CRISPRa can activate both the core σ^70^ (RpoD) promoter and other non-housekeeping sigma factor promoters, which includes >95% of *E. coli* promoters.^6^ To investigate whether the design rules for SoxS-mediated CRISPRa are generalizable to the other core promoters, we also screened σ^38^ (RpoS), σ^32^ (RpoH), σ^24^ (RpoE), and σ^54^ (RpoN) promoters at 10 bp resolution. We found that CRISPRa at σ^38^, σ^32^, and σ^24^ promoters is optimal in the same region as σ^70^ promoters, 60-100 bp upstream of the TSS (Figure S1A). For the σ^54^ promoter, which represents ∼3% of *E. coli* promoters, we observed no SoxS-mediated activation at any tested sites. To determine whether the sharp phase dependence observed with σ^70^ promoters was present at other sigma factor promoters, we screened the σ^38^-dependent sodCp promoter at 1 bp resolution. We found that SoxS-mediated CRISPRa maintains the same optimal target site and sharp phase dependence across σ^70^ and σ^38^ promoters (Figure S1B). These findings suggest that the observed distance preference for SoxS is inherent to the effector itself rather than the minimal sigma factor promoter.

SoxS orthologs are also potentially useful for bacterial CRISPRa. Here we focused on two SoxS orthologs in the AraC/XylS family, MarA and TetD. Previously, we observed modest CRISPRa with TetD and no detectable activation with MarA.^27^ We attempted to improve activation by mutating the native DNA binding motifs to prevent the effectors from being sequestered at off-target endogenous sequences. Although this strategy was effective with SoxS,^6,27^ the TetD mutants and MarA mutants exhibited no significant increase in CRISPRa (Figure S2). However, TetD remains a candidate for synergistic CRISPRa. Because TetD prefers similar target sites to SoxS,^27^ it could potentially be effective when recruited with SoxS to the same target site.

### Design rules for RNAP subunits and assembly mediators

RNAP subunits can act as synthetic transcriptional activators in bacteria through subunit-mediated recruitment of the functional RNAP holoenzyme.^46^ There are multiple strategies for subunit-mediated recruitment of RNAP to the CRISPR complex, including direct fusion to dCas9^26,30^ and noncovalent recruitment via protein-protein or protein-gRNA interactions.^15,27^ Several RNAP subunits have been characterized in the J1 reporter context.^6,15,27^ In this work, we focus on the N-terminal domain of the alpha subunit (RpoA), termed αNTD, because it has been the most effective and widely used of the RNAP subunits.^5,15,27,30^ αNTD is effective when recruited as an αNTD-MCP fusion to an MS2 scRNA, and has similar target site preferences to SoxS (Figure 2A).^6,27^

We first evaluated whether αNTD would be functional when recruited with a different RNA hairpin/RNA-binding protein pair. This approach could be used to independently control different genes, or to recruit multiple effectors to different sites on the same promoter. We tested an αNTD-PCP fusion protein, which can be recruited to a PP7 scRNA. αNTD-PCP exhibited similar target site preferences but lower activity compared to αNTD-MCP (Figure S3). This behavior is analogous to previous results with PCP-SoxS, which has similar target site preferences but modest activity compared to MCP-SoxS.^27^ These results provide the design rules necessary to test an orthogonal programming strategy where one CRISPR complex recruits αNTD-PCP and another recruits MCP-SoxS, or vice versa with αNTD-MCP and PCP-SoxS.

In addition to αNTD, RNAP assembly can be facilitated by other RNAP subunits or RNAP assembly factors, with moderate effectiveness in CRISPRa.^11,26,27,47^ In this work, we evaluated an additional candidate effector, the RNAP-σ^38^ assembly factor Crl.^48^ We recruited MCP-Crl to a σ^38^-dependent promoter and observed a small, 1.2-fold increase in gene expression compared to an off-target scRNA (Figure S4). We also found that the presence of MCP-Crl alone, in the absence of any recruitment to the CRISPR complex, increases reporter gene expression by 2.4-fold relative to a strain without MCP-Crl. This effect could arise from nonspecific Crl-mediated activation of σ^38^ promoters. Thus, Crl may not be a useful CRISPRa effector, and αNTD remains the most effective of the RNAP subunits or assembly factors tested for CRISPRa.

### Design rules for AsiA

AsiA is an anti-σ^70^ factor that can activate transcription when fused to heterologous DNA binding domains.^27,49,50^ An evolved variant AsiA_m2.1 mitigates the toxicity of wild-type AsiA and is effective for CRISPRa as a dCas9 fusion;^13^ this variant was used for all AsiA experiments in this work. dCas9-AsiA has been reported to preferentially activate at target sites 90-252 bp from the TSS, further than the 60-100 bp range for SoxS^13^. These findings were reported with a W1 reporter; we confirmed these target site preferences and observed optimal CRISPRa with dCas9-AsiA expressed at basal levels from a pTet promoter in the absence of an inducer (Figure S5).^26^ We then tested dCas9-AsiA using the J1 reporter with target sites 40-140 bp from the TSS, but we were unable to detect activation at target sites within this range (Figure 2A). To evaluate target sites further upstream of our original J1 reporter with dCas9-AsiA, we constructed novel synthetic promoters with the J1 array of gRNA target sites placed 100 bp upstream of the J3 promoter. The J3 promoter contains a set of target sites at optimal positions for SoxS-mediated CRISPRa.^6^ The new J1-J3(100) hybrid promoter allows us to screen AsiA target sites further than 140 bp at the upstream J1 sites, while maintaining activity with SoxS, αNTD, and TetD at the downstream J3 sites (Figure S6 & S7A). We observed up to 3.2-fold activation at target sites positioned from - 160 to -180 bp (Figure S7B), much less than the 21-fold activation observed at similar positions with the W1 reporter (Figure S5C). We also constructed another hybrid promoter J1-J2(100)-J3(100), which shifts the J1 target sites to 240-340 bp, allowing us to test the same distances as previously reported. With the J1-J2(100)-J3(100) promoter, we still observed only 1.9-fold activation at position -250 (Figure S7B). This small effect is consistent with our observations in the W1 reporter of weaker activation at H5 (−252bp) compared to H4 (−182 bp) (Figure S5C).

Because bacterial CRISPRa can be ineffective at target sites shifted by 1 or 2 bases from optimal sites,^6,13^ we constructed the J1-J3(122) promoter, which is shifted by 22 bases from J1-J3(100). This shift moves target sites to -182 bp and -252 bp, which correspond precisely to the optimal positions previously reported with the W1 reporter. Using J1-J3(122), we tested target sites from 162-262 bp upstream of the TSS with dCas9-AsiA. We observed strong, ∼30-fold gene activation at -192 or -202 bp target sites, (Figure S7C). Surprisingly, we did not detect significant activation from other target sites, including -182 bp which was the most effective site reported previously.^13^ We also found that dCas9-AsiA activation was biased toward the template strand, unlike the previously reported results where activation was observed from template and non-template strand targets. Our observations qualitatively agree with previous results for dCas9-AsiA target site preferences,^13^ but we find that the precise positions of effective sites can vary between promoters. Characterizing AsiA and SoxS in the same promoter context is thus a critical step towards a potential multi-effector CRISPRa strategy with AsiA and SoxS.

### Design rules for PspF

σ^54^ promoters are inaccessible to SoxS, but they can be activated by PspF.^6,14^ PspF is an ATP-dependent transcriptional activator that positively regulates *E. coli psp* operon σ^54^ promoters in response to cellular stresses.^51^ To benchmark PspF activity, we evaluated PspF-mediated CRISPRa with a previously-characterized CRISPRa-responsive σ^54^ promoter, pspAp-G6.^14^ We observed ∼16-fold activation from PspF-mediated CRISPRa (Figure S8A). Previous work has reported much larger fold-activation values for PspF-mediated CRISPRa,^14^ but these values were obtained using a different baseline subtraction method. When the raw data are directly compared, our results are consistent with the previously reported values (Figure S8A).

To systematically screen PspF target site distance preferences, we constructed a modified σ^54^ promoter with a J1 array of target sites upstream of a minimal σ^54^ minimal promoter (J1-pspAp). At the most effective target site, located at position -61 on the non-template strand, we observed 8-fold activation (Figure 2B). Compared to pspAp-G6, the optimal target site in J1-pspAp is shifted 30 bases closer to the TSS. This shift could be because J1-pspAp lacks the integration host factor (IHF) binding site that forms a loop to bring PspF closer to the minimal promoter.^52^

We proceeded to screen target site preferences in J1-pspAp at single base resolution. We observed a similar periodic effect as observed previously for both PspF and SoxS,^14,28^ and we observed a modest increase in activity at position -60 compared to -61 (Figure S8B). We proceeded to use the -60 target site position for subsequent PspF experiments. Because PspF has distinct target site preferences compared to SoxS and other activators, we expect that it may not be useful as a multi-effector fusion protein targeted to a single site. However, PspF might be effective when multiple, orthogonal CRISPR complexes are used to target different activators to their respective preferred target sites.

### Design rules guide strategies to combine CRISPRa effectors

We employed three strategies to combine CRISPRa systems: 1) tandem protein fusion of multiple effectors, 2) recruiting two effectors to a single CRISPR complex using a combination of direct dCas9 fusion and gRNA recruitment, and 3) targeting two orthogonal CRISPRa complexes to different preferred target sites. We used the optimal positioning for each CRISPRa activator to guide our choices on which activator combinations to test with each strategy (Table 1 and Table S7).

**Table 1.**
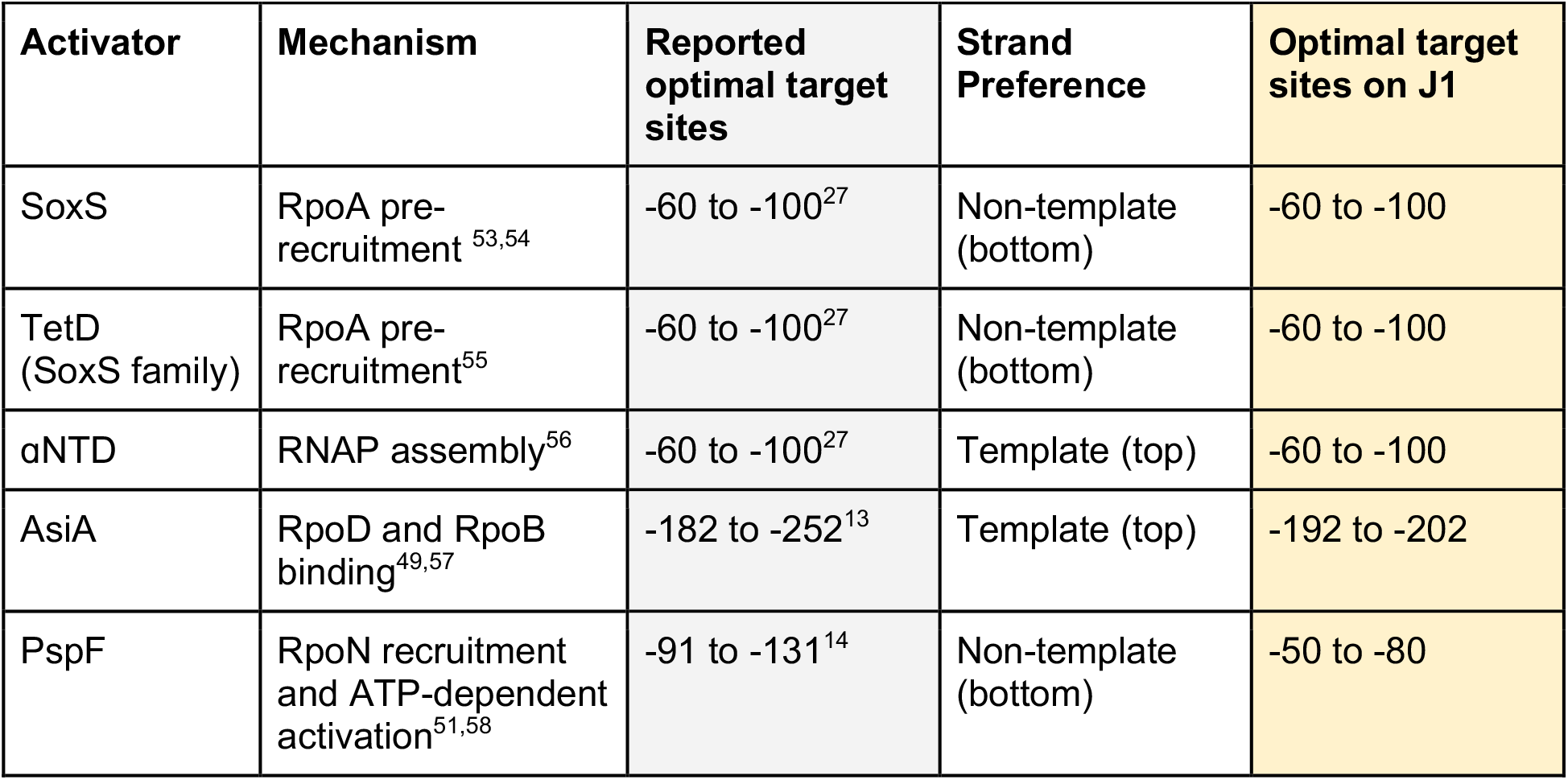
Design rules for different CRISPRa effectors.

| Activator | Mechanism | Reported optimal target sites | Strand Preference | Optimal target sites on J1 |
| --- | --- | --- | --- | --- |
| SoxS | RpoA pre-recruitment<sup>53,54</sup> | -60 to -100<sup>27</sup> | Non-template (bottom) | -60 to -100 |
| TetD (SoxS family) | RpoA pre-recruitment<sup>55</sup> | -60 to -100<sup>27</sup> | Non-template (bottom) | -60 to -100 |
| αNTD | RNAP assembly<sup>56</sup> | -60 to -100<sup>27</sup> | Template (top) | -60 to -100 |
| AsiA | RpoD and RpoB binding<sup>49,57</sup> | -182 to -252<sup>13</sup> | Template (top) | -192 to -202 |
| PspF | RpoN recruitment and ATP-dependent activation<sup>51,58</sup> | -91 to -131<sup>14</sup> | Non-template (bottom) | -50 to -80 |

### Fusion of tandem effectors decreases CRISPRa activity

To test tandem effector fusions, we used SoxS with two alternative activators, TetD and αNTD, that have similar target preferences. This strategy is comparable to the eukaryotic dCas9-VPR system, in which three activators are fused to dCas9 in tandem to produce a synergistic CRISPRa effect.^20^ For bacterial CRISPRa, we used a single MS2 scRNA to recruit multi-effector MCP fusion proteins. For SoxS and TetD, we constructed a C-terminal fusion MCP-SoxS-TetD because the individual C-terminal fusion MCP-SoxS and MCP-TetD are effective.^27^ We found that at their most effective target sites, MCP-SoxS-TetD produced 7-fold gene activation, while the individual MCP-SoxS or MCP-TetD effectors produced 27-fold or 5-fold activation, respectively.

Thus, MCP-SoxS-TetD is modestly more effective than MCP-TetD alone, but significantly impaired when compared to MCP-SoxS (Figure S9). In a similar strategy with SoxS and αNTD, we appended SoxS to the previously validated αNTD-MCP to generate αNTD-MCP-SoxS. We found that at their most effective target sites, αNTD-MCP-SoxS produced 1.3-fold gene activation, which was relatively weak compared to both αNTD-MCP (3.3-fold) and MCP-SoxS (27-fold) (Figure 3A). Our findings are consistent with previous unsuccessful attempts to improve bacterial gene activation using tandem SoxS and αNTD^29^ or tandem α and ω subunit effectors.^30^

**Figure 3.**
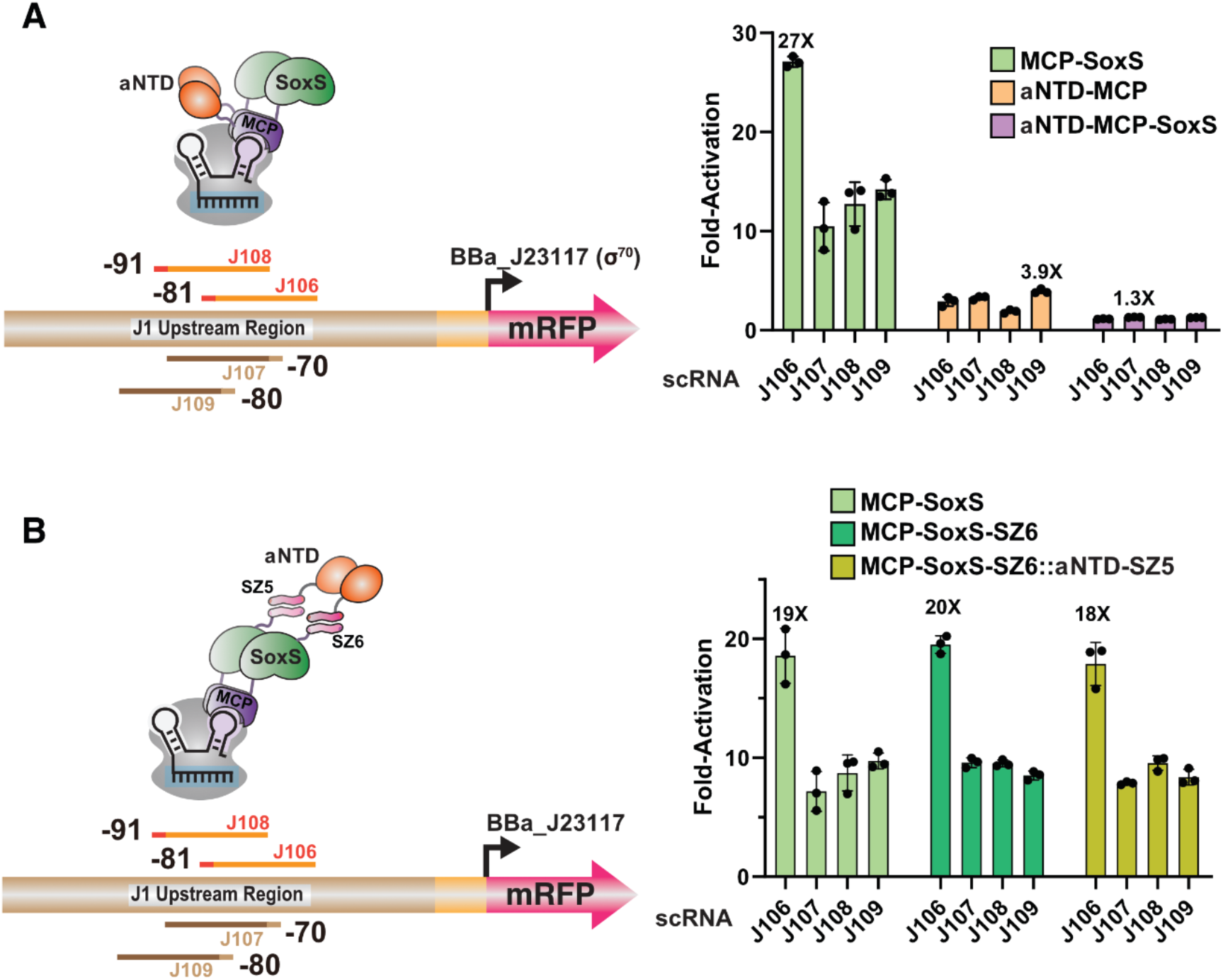
Combined recruitment of SoxS and αNTD is antagonistic or ineffective. **A)** Fusion of αNTD-MCP-SoxS (purple) yielded diminished CRISPRa activity compared to αNTD-MCP alone (orange) or MCP-SoxS alone (green) alone. **B)** SYNZIP-mediated recruitment of αNTD to MCP-SoxS does not increase gene expression compared to MCP-SoxS. Values in panel A and B represent the mean ± standard deviation calculated from n = 3.

We tested an alternative system for simultaneous recruitment of αNTD and SoxS using a SYNZIP pair to recruit αNTD to MCP-SoxS. SYNZIP recruitment has previously been effective for bacterial CRISPRa.^8,15^ We observed effective CRISPRa with αNTD-SYNZIP5/MCP-SYNZIP6, confirming that SYNZIP recruitment of αNTD can activate gene expression (Figure S10). However, gene activation did not increase with αNTD-SYNZIP5/MCP-SoxS-SYNZIP6 compared to MCP-SoxS or MCP-SoxS-SYNZIP6 (Figure 3B). Taken together, these results suggest that bacterial CRISPRa may be sensitive to the precise orientation or positioning of the activator fusion protein, and we lack sufficient information to design an effective tandem multi-effector fusion.

### Combined recruitment of AsiA and SoxS is antagonistic

We proceeded to test whether SoxS and AsiA activators could be effective when recruited to a single CRISPR complex. To recruit both activators, we used a dCas9-AsiA fusion and an MS2 scRNA to recruit MCP-SoxS to the same complex. This approach is analogous to the eukaryotic CRISPRa SAM system.^21^ However, AsiA and SoxS have non-overlapping optimal target sites (Figure 2), so we had low confidence that this strategy would improve activation. We targeted dCas9-AsiA/MCP-SoxS to an optimal AsiA site but observed no increase in gene activation compared to dCas9-AsiA alone (Figure 4A & S11). Unexpectedly, when dCas9-AsiA/MCP-SoxS was targeted to an optimal SoxS site, we observed a substantial decrease in gene expression compared to only dCas9/MCP-SoxS system (Figure 4A). We also targeted dCas9-AsiA/MCP-SoxS to both the optimal AsiA and SoxS target sites simultaneously by expressing two scRNAs, but again observed a substantial decrease relative to dCas9/MCP-SoxS (Figure 4A). These observations suggest that if SoxS and AsiA are recruited to the same target site, they may have incompatible interactions with RNAP that prevent optimal gene activation.

**Figure 4.**
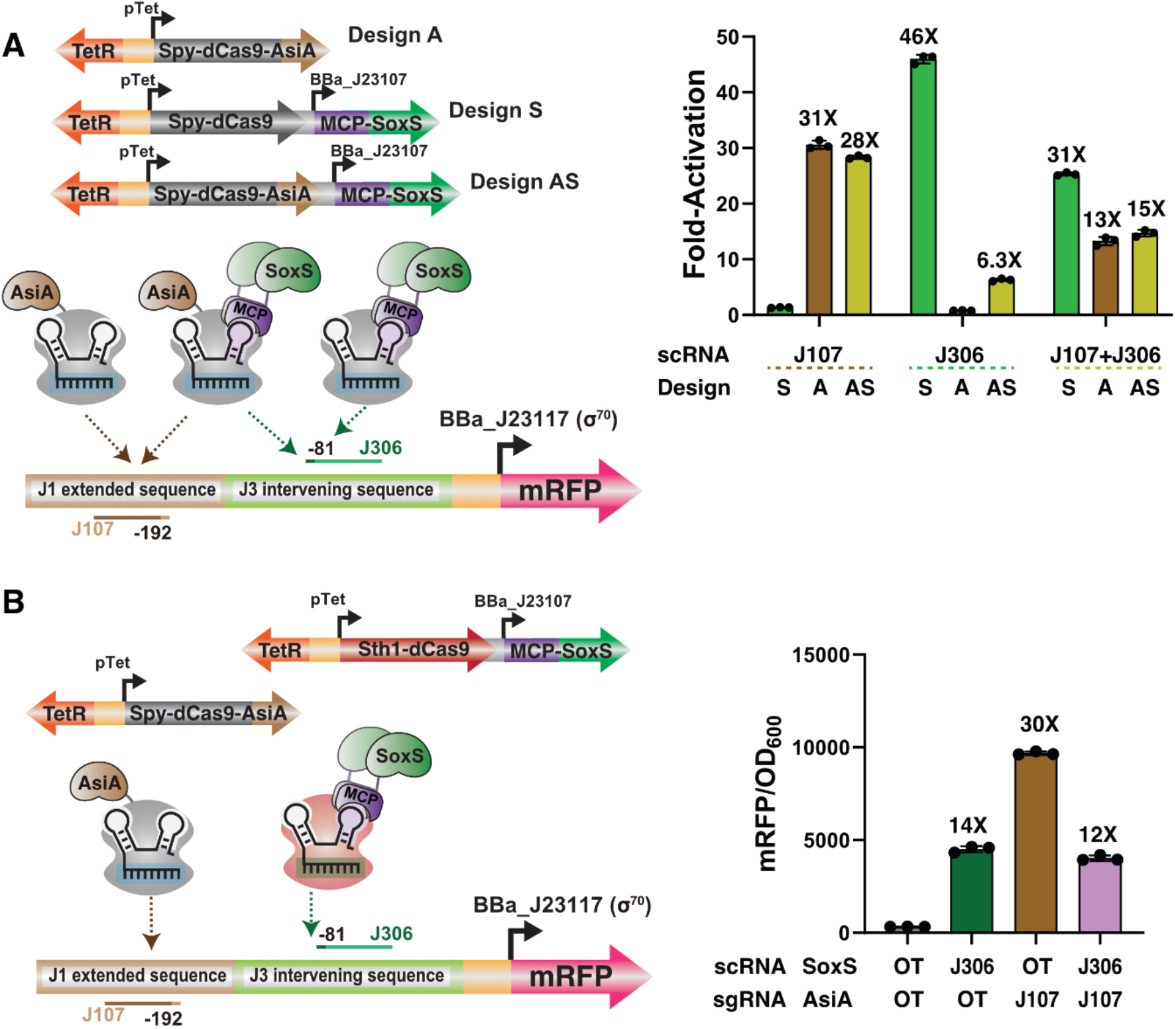
SoxS and AsiA CRISPRa exhibit antagonistic effects. **A)** dCas9-AsiA, dCas9/MCP-SoxS, or dCas9-AsiA/MCP-SoxS recruitment to optimal target sites for AsiA (J107, -192) or SoxS (J306, -81). At the optimal AsiA site, dCas9-AsiA/MCP-SoxS does not improve activation relative to dCas9-AsiA. At the optimal SoxS site, dCas9-AsiA/MCP-SoxS is substantially impaired compared to dCas9/MCP-SoxS. With both gRNAs expressed to target the CRISPRa complex to both sites simultaneously, dCas9-AsiA/MCP-SoxS is impaired compared to dCas9/MCP-SoxS and roughly equivalent to dCas9-AsiA. **B)** Targeting AsiA and SoxS to their respective optimal sites with Spy-dCas9-AsiA and Sth1-dCas9/MCP-SoxS. Combined AsiA/SoxS produced lower activation (12-fold) than that observed with either AsiA alone (30-fold) or SoxS alone (14-fold). Values in panel A and B represent the mean ± standard deviation calculated from n = 3.

As an alternative strategy, we used orthogonal CRISPR complexes to target AsiA and SoxS separately to their respective optimal target sites. The current AsiA and SoxS CRISPRa complexes are not orthogonal, with AsiA recruited as an *S. pyogenes* dCas9 (Spy-dCas9) fusion and SoxS recruited to Spy-dCas9 with a scRNA. Therefore, we needed to develop alternative orthogonal targeting mechanisms. One option could be recruitment of SoxS and AsiA to different sites with Spy-dCas9 and orthogonal scRNAs, so we tested MS2 scRNA-mediated recruitment of MCP-AsiA. However, we observed only small, <1.5-fold CRISPRa effects (Figure S12). Another option could be fusion of AsiA to an alternative, orthogonal Cas protein. We tested AsiA fusions to alternative dCas9 ortholog Sth1-dCas9 (Figure S13) or to dCas12a (Figure S14). There was no detectable activation with Sth1-dCas9 fusion and small, <2-fold CRISPRa effects with dCas12a fusions. Finally, we used MS2 scRNAs to recruit SoxS to the Sth1-dCas9 ortholog.^38^ We tested three engineered MS2 scRNA variants and obtained 8-fold gene activation from the scRNA design analogous to the most effective MS2 scRNA for Spy-dCas9 (Figure S15). Although the 8-fold activation with Sth1-dCas9/MCP-SoxS is smaller than the 20-to 40-fold activation effects observed with Spy-dCas9/MCP-SoxS in the same promoter context (Figure S6A & S9A), the Sth1 system provides sufficient activation to proceed with testing as part of a multi-activator CRISPRa system.

We proceeded to deliver AsiA and SoxS to their respective optimal target sites using orthogonal dCas9 proteins for each effector, Spy-dCas9-AsiA and Sth1-dCas9/MCP-SoxS (Figure 4B). We used the modified J1-J3(122) promoter with an Sth1-dCas9-compatible PAM at the -81 target (J306) and Spy-dCas9-compatible PAM at the -192 target. With orthogonal dCas9s and both AsiA and SoxS, we observed ∼12-fold gene activation, which is lower than the activation observed with either AsiA alone (30-fold) or SoxS alone (14-fold). Thus, there is an antagonistic effect between SoxS and AsiA even when they are recruited by orthogonal CRISPRa complexes placed >100 bp away from each other. From these results, we believe that there might be competitive, incompatible interactions with RNAP from SoxS and AsiA together. We also cannot rule out the possibility that AsiA and SoxS directly interact and interfere with each other. Although our design rules suggest plausible synergistic activation with AsiA and SoxS, we are unable to design an effective multi-activator CRISPRa system.

### Combined recruitment of SoxS and PspF is ineffective

Finally, we evaluated recruitment of the SoxS and PspF activators, each to their respective optimal target site. Because these established systems are based on orthogonal engineered scRNAs, we can recruit SoxS to one dCas9 complex and PspF to another. However, SoxS and PspF have apparently incompatible promoter requirements. PspF interacts directly with σ^54^ and is effective only at σ^54^-dependent promoters (Figure 2),^14,59^ while SoxS functions at most promoters except σ^54^ (Figure S1A). The ineffectiveness of SoxS at σ^54^ promoters is unsurprising because σ^54^ RNAP requires ATP-dependent remodeling.^58^ However, it might be possible for SoxS to enhance PspF-dependent activation at a σ^54^ promoter, with PspF engaging directly with the sigma factor while SoxS interacts with the RpoA subunit. We therefore tested simultaneous recruitment using J1-pspAp, the modified σ^54^ promoter with an upstream array of J1 gRNA target sites. As SoxS and PspF prefer distinct minimal promoters, simultaneous recruitment in tandem fusion to the same target site was ineffective (Figure S16).

PspF and SoxS have optimal target sites that are relatively close, at -60 and -81 from the TSS, respectively. Because these sites might overlap, we targeted PspF with a boxB scRNA to its optimal site at -60 and targeted SoxS with an MS2 scRNA to -81 or to additional upstream sites (Figure 5A). This strategy was chosen because SoxS retains activity at some sites upstream of its optimal -81 site (Figure 2A). Several SoxS target sites produced improvements relative to PspF alone. The largest effect was with PspF at -60 and SoxS at -100, which activated gene expression 18-fold, a 1.8-fold improvement relative to PspF alone (Figure 5B, 5C, & S16). Notably, SoxS alone produces no detectable activation with this promoter, indicating a potential synergistic 1.8-fold improvement. We observed similar effects when SoxS was replaced with αNTD or TetD (Figure 5D, S17, & S18). To determine if this apparent synergistic effect results from recruitment of both activators, we targeted dCas9 alone to the -100 bp site with an sgRNA or an MS2 scRNA and kept recruiting PspF to the -60 bp site. Without SoxS, we observed 1.2-fold and 1.6-fold increases relative to PspF alone (Figure 5E). With these results in mind, we believe the 1.8-fold synergistic effect arises from a combination of small effects: 1.2-fold from recruiting a second dCas9 complex upstream of PspF, 1.3-fold (1.6/1.2) from the MS2 hairpin, and 1.1-fold (1.8/1.6) from SoxS recruitment together with PspF. Taken together, we observed what appears to be a true synergistic effect for the first time with bacterial CRISPRa, but the total 1.8-fold effect is relatively small and only 1.1-fold of the effect comes from the cooperative action of the PspF and SoxS activators.

**Figure 5.**
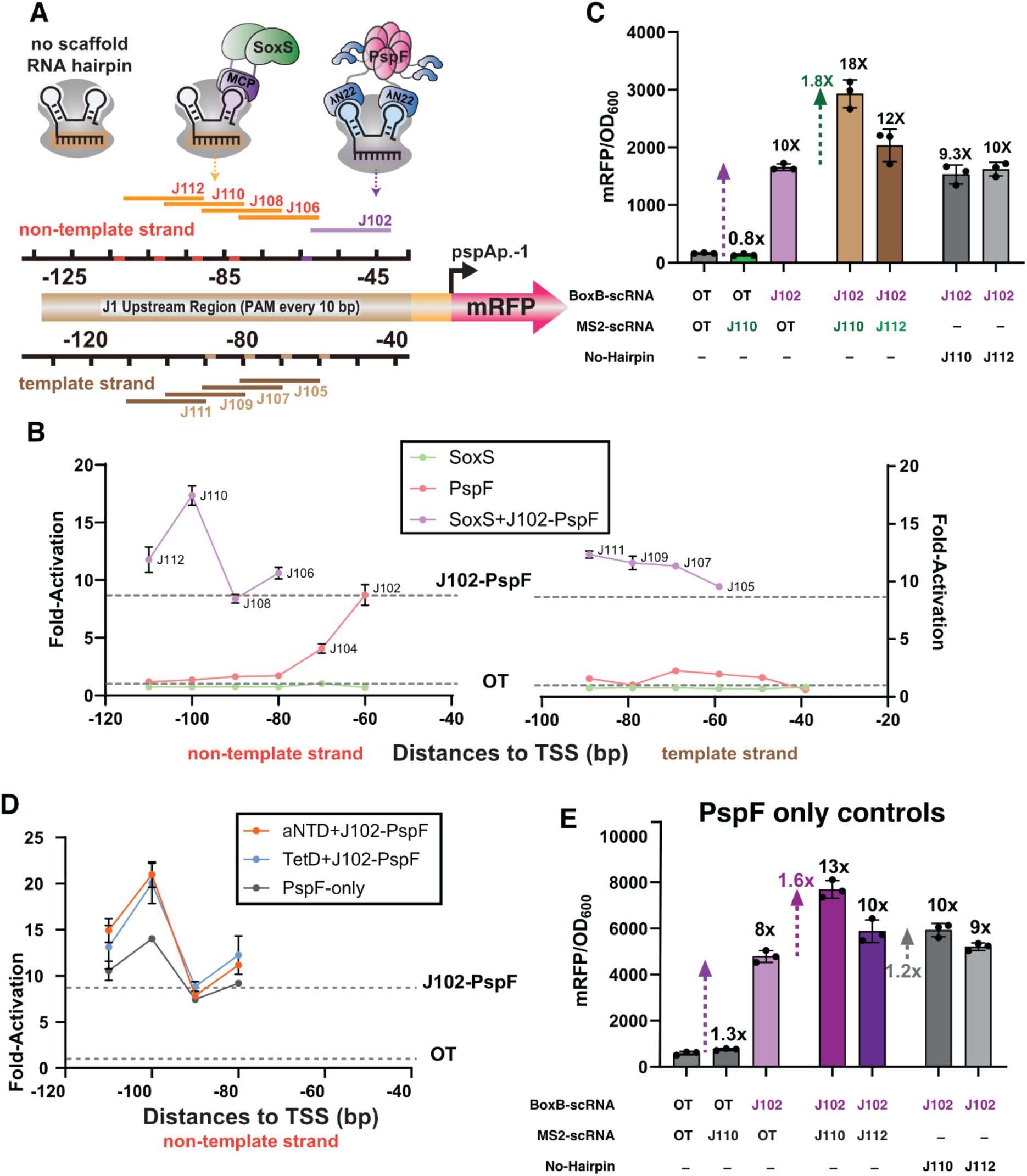
Combined recruitment of PspF and SoxS is ineffective. **A**) MCP-SoxS and PspF-λN22 CRISPRa complexes are recruited with orthogonal MS2 and boxB scRNAs, respectively, to the σ^54^ J1-pspAp-1 promoter (Figure S8B). PspF is targeted to its optimal position -60 bp (J102) upstream of TSS and the SoxS-mediated CRISPR complex was screened at target sites from - 60 to -110 bp. **B)** CRISPRa with SoxS alone, PspF alone, or SoxS and PspF at a range of sites with the σ^54^ J1-pspAp-1 promoter. For simultaneous recruitment of PspF and SoxS, PspF is fixed at -60 and the x-axis refers to the target site for SoxS. **C)** Comparison of SoxS alone, PspF alone, or SoxS and PspF with PspF at -60 (J102) and SoxS at -100 (J110) or -110 (J112). With SoxS at -100 and PspF at -60, there is an 18-fold increase in gene expression, above what would be predicted from the combination of SoxS alone (0.8-fold) and PspF alone (10-fold). This increase is dependent on the presence of the MS2 hairpin, which recruits SoxS to the J110 or J112 target sites. **D)** CRISPRa with αNTD+PspF, TetD+PspF, or PspF alone. For simultaneous recruitment of PspF with αNTD or TetD, PspF was fixed at -60 and the x-axis refers to the target site for αNTD or TetD. With αNTD or TetD at -100 and PspF at -60, there is a 20-fold increase in gene expression, similar to the effect observed with SoxS and PspF. **E)** Controls for PspF CRISPRa in the absence of a second activator. With PspF recruited to -60, recruitment of dCas9 to the -100 site with an sgRNA increases gene expression 1.2-fold, and using an MS2 scRNA at -100 increases gene expression 1.6-fold relative to PspF alone. Values in panels B-E represent the mean ± standard deviation calculated from n = 3.

## Discussion

To improve gene activation with bacterial CRISPRa, we attempted to recruit multiple effectors to simultaneously engage RNAP and cooperatively activate transcription.^23,24^ To implement this strategy, we first needed to know where to recruit each effector in the region upstream of the promoter. Bacterial CRISPRa systems typically have strict requirements for target site positions,^6,13,14^ so we first systematically mapped preferred target sites for different CRISPRa systems (Figure 2 and Table 1). We used a set of promoters compatible with multiple CRISPRa systems, which was critical because the preferred target sites for each CRISPRa system can vary in different promoter contexts. Despite defining optimal target sites, we did not observe any large synergistic effects on gene activation. When we recruited the potent activator SoxS together with other SoxS-family activators or with RNA polymerase subunits at the same target site, we observed either impaired or unchanged activation compared to SoxS alone (Figure 3). We observed similar behavior when recruiting the two potent activators SoxS and AsiA at distinct target sites (Figure 4). These antagonistic effects could arise if the RNAP interactions are incompatible and cannot form simultaneously. Finally, we observed a modest but potentially significant ∼2-fold synergistic effect when both SoxS and PspF were recruited together (Figure 5). However, this effect arises largely due to simultaneous binding of two CRISPR-Cas complexes, and only a small fraction of the effect is due to the simultaneous action of both SoxS and PspF. We also expect this effect to have limited practical utility at endogenous bacterial gene targets because it is restricted to σ^54^ promoters, which constitute a small fraction of the bacterial genome. Thus, despite careful analysis to identify effective combinations of effectors for synergistic CRISPRa, we remain largely unable to improve upon the gene activation that can be obtained with individual CRISPRa systems.

Our findings emphasize the challenges of bacterial CRISPRa compared to eukaryotic CRISPRa. Prior to the advent of CRISPR-Cas gene regulation, it was well understood that eukaryotic gene activation could be enhanced by recruiting multiple effectors together.^17,19,60,61^ These insights were readily transferred to eukaryotic CRISPRa systems, where recruiting multiple effectors produces enhanced and frequently synergistic activation.^17–21,62,63^ Similar effects in bacterial CRISPRa systems have not emerged, possibly due to fundamental differences in the mechanisms of gene activation. In eukaryotes, transcription factors interact with multi-protein complexes of coactivators and chromatin modifiers that in turn recruit RNAP.^64,65^ Eukaryotic CRISPRa effectors that interact with these complexes can be effective at a broad range of upstream target sites,^66,67^ and it is therefore straightforward to enhance gene activation with multiple CRISPRa effectors. In contrast, bacterial CRISPRa effectors typically directly contact RNAP and are subject to strict positioning requirements for productive gene activation.^6,13,14^ These precise positioning requirements have made it difficult to identify compatible positions to recruit multiple bacterial effectors. Although we systematically identified sites that should be compatible, we were unable to enhance bacterial CRISPRa with multiple effectors, suggesting that other structural constraints or unknown factors remain to be addressed.

In principle, synergistic CRISPRa in bacteria should be possible because native transcriptional activators can produce synergistic effects. The best characterized system is the *E. coli* catabolite activator protein (CAP), also known as the cAMP receptor protein (CRP).^24,31,33,34,68,69^ Two appropriately positioned CAP sites can simultaneously contact RNAP to synergistically activate transcription. Based in part on this precedent, we originally screened CAP as a potential CRISPRa effector in *E. coli*, but we did not detect any significant transcriptional activation.^27^ Given the efficacy of CAP as a synergistic activator, and the failure of existing CRISPRa effectors to achieve synergistic activation, it may be worth revisiting CAP as a potential CRISPRa effector, either with engineered variants or homologs. More broadly, it could be informative to generate structural models of candidate CRISPRa complexes with RNAP and promoter DNA, although the sizes of such complexes are beyond the limits of the current publicly-available Alphafold3 server.^70^ These structural models could help to identify combinations of CRISPRa complexes that could simultaneously and cooperatively engage with RNAP to initiate transcription. Ultimately, we hope these approaches will help us towards the long-term goal to achieve predictable and robust activation of any desired endogenous genes in bacteria.

## Supporting information

The Supporting Information is available free of charge.

Supporting methods, CRISPRa effects with alternative effectors and reporters, bacterial strains, plasmids, guide RNA spacer sequences, and sequences of reporter genes, proteins, and guide RNAs.

## Authors Contribution

C.K., A.V.K., R.C., K.O., J.M.C. and J.G.Z. conceptualized the research; C.K., A.V.K., R.C., K.O., P.L. and S.S.A. performed experiments; C.K., A.V.K., R.C. and K.O. analyzed the data. C.K., A.V.K., R.C., J.M.C., and J.G.Z. wrote the manuscript with input from all of the authors.

## Acknowledgements

We thank members of the Carothers and Zalatan groups for helpful discussion. This work was supported by NSF Award MCB 1817623 (J.G.Z. and J.M.C.), MCB 2032794 (J.M.C. and J.G.Z.), CBET 1844152 (J.M.C.), DOE Awards DE-EE0008927 and DE-SC0023091 (J.M.C, J.G.Z.). This material is based upon work supported by the National Science Foundation Graduate Research Fellowship Program under Grant No. DGE-2140004 (A.V.K.). Any opinions, findings, and conclusions or recommendations expressed in this material are those of the author(s) and do not necessarily reflect the views of the National Science Foundation.

