## Supporting Information for "Systematic Mapping of Bacterial CRISPRa Design Rules and Implications for Synergistic Gene Activation"

a: Molecular Engineering & Sciences Institute
University of Washington
Seattle, WA 98195
United States

b: Center for Synthetic Biology
University of Washington
Seattle, WA 98195
United States

c: Department of Chemical Engineering
University of Washington
Seattle, WA 98195
United States

d: Department of Bioengineering
University of Washington
Seattle, WA 98195
United States

e: Department of Chemistry
University of Washington
Seattle, WA 98195
United States

f: Department of Biochemistry
Mahidol University
Bangkok, 10400
Thailand

g: Current Address:
Institute for Medical Engineering and Science
Massachusetts Institute of Technology
Cambridge, MA, 02139
United States

*: These authors contributed equally
*: Corresponding authors

206-221-4902


206-543-1670

**Table of Contents**


**Supporting Methods** **3**

Multiple-plasmid systems for CRISPRa screening in E. coli 3

AsiA recruitment via engineered gRNA 3

Systematically tiled promoters for Sth1-dCas9 and Lb-dCas12a 3

**Supporting Figures** **5**

Figure S1. Target site screening for SoxS-mediated CRISPRa at alternative σ-factor promoters 5

Figure S2. CRISPRa with MarA and TetD mutants 7

Figure S3. αNTD-mediated CRISPRa with RNA-binding protein recruitment 8

Figure S4. RNAP-σ38 assembly using MCP-Crl 9

Figure S5. CRISPRa with dCas9-AsiA 10

Figure S6. Comparison of different CRISPRa systems at J1, J3, and J1-J3(122) promoters 11

Figure S7. dCas9-AsiA based CRISPRa at target site positions in J1, J3, and J1-J3 promoters 13

Figure S8. CRISPRa with PspF-λN22 at pspAp-G6 and J1 reporters 14

Figure S9. CRISPRa with tandem activator fusions 15

Figure S11. SoxS and AsiA CRISPRa exhibit antagonistic effects 17

Figure S12. CRISPRa with an MS2 scRNA to recruit MCP-AsiA 18

Figure S13. CRISPRa with Sth1-dCas9-AsiA is not effective 19

Figure S14. Modest CRISPRa with dCas12a-AsiA 20

Figure S15. CRISPRa with Sth1-dCas9/MCP-SoxS and MS2 scRNAs. 22

Figure S16. CRISPRa with PspF-SoxS fusions 23

Figure S17. Characterization of PspF CRISPRa with alternative activators 24

Figure S18. Characterization of CRISPRa with PspF+αNTD and PspF+TetD. 25

**Supporting Tables** **26**

Table S1. Bacterial strains used in this work 26

Table S2. List of plasmids used in this work 27

Table S3. sgRNA/scRNA/crRNA spacer sequences 32

Table S4. sgRNA target site positions for alternative promoters 34

Table S5. Plasmids used in each figure 35

Table S6. Design rules of different CRISPRa systems not tested in this work (extended from Table 1) 40

Table S7. Potential cooperativity of effectors investigated in this work 41

**DNA sequences** **42**

CRISPRa construct sequences: dCas proteins and activators 42

Systematic Reporters 56

sgRNA/scRNA/crRNA 60

**Supporting References** **62**

### Supporting Methods

#### Multiple-plasmid systems for CRISPRa screening in *E. coli*

Three plasmids with different replicons and antibiotic markers were used in this study; 1) p15A-CmR, 2) pSC101-AmpR, and 3) ColE1-KmR. For all experiments, dCas9 or dCas9-AsiA were expressed from p15A-CmR. All RBP-activators were expressed on a p15A-CmR plasmid together with dCas9 constructs. Multiple activators were always constructed in a single p15A-CmR plasmid. mRFP reporters with systematically tiled promoters were always on low-copy pSC101-AmpR plasmids. sgRNAs or scRNAs (sgRNAs modified with RNA hairpins) were expressed on ColE1-KmR when delivered as part of a three plasmid system. In some cases, sgRNAs/scRNAs were incorporated into p15A-CmR or pSC101-AmpR for delivery as a two plasmid system. For expression of multiple scRNAs, both gRNA expression cassettes were co-expressed from the same ColE1-KmR plasmid. Details for each plasmid are available in Table S2. Plasmids used for each figure are also listed in Table S5.

#### AsiA recruitment via engineered gRNA

Following previous work, we used an MS2 scRNA to recruit MCP to the CRISPRa complex.[1,2] In the original report by Dong et al.,[1] expression of wild type AsiA fused with MCP was toxic unless coexpressed with the RpoD F563Y mutant. In the evolved AsiA_m2.1 variant developed by Ho et al.,[2] MCP-AsiA was successfully expressed with an aTc-inducible promoter. When we initially attempted to express MCP-AsiA_m2.1 with a medium constitutive promoter (BBa_J23107), we observed multiple mutations similar to that observed previously when attempting to express wt AsiA.[1] We therefore expressed MCP-AsiA with an aTc-inducible promoter. However, this scRNA-based system showed negligible CRISPRa-mediated gene activation (<1.5-fold) (Figure S12), consistent with previous reports.[2]

#### Systematically tiled promoters for Sth1-dCas9 and Lb-dCas12a

For CRISPRa with alternative Cas proteins, new promoter systems were designed for systematic characterization using a strategy previously used for J1 sequence design.[1] For *Streptococcus thermophilus* dCas9 fused with AsiA_m2.1, termed Sth1-dCas9-AsiA in this manuscript, we used custom Python scripts to generate 180 nt tiling arrays with NNAGAAW PAM site every 20 nt with 5'-NNAGAAWNNNNNNNNNNNNN-3' as a 20 base spacer sequence for each sgRNA (each 20 base spacer is followed by the NNAGAAW PAM, so the next 20 base spacer begins with NNAGAAW). We randomly generated 100,000 sequences of 180 nt and filtered for endogenous GC content of *E. coli* (50.8%). We discarded any sequences with four consecutive identical nucleotides and sequences containing known *E. coli* transcription factor binding sites.[3] Finally, we omitted sequences that have common restriction enzyme sites and arbitrarily selected one sequence, referred as K1, for further systematic screening with a corresponding set of sgRNAs (K101-K108). We constructed a new promoter (K1T) by modifying the J1-J3(122bp) promoter to replace the upstream J1 array with the K1 array. The K1T promoter has protospacer sequences on the top strand (Figure S13). We constructed a variant promoter K1B with the 180 nt K1 sequence flipped so that the protospacer sequences were on the bottom strand. We also constructed variants of each promoter shifted by 10 nt (K1T+10 and K1B+10). For MCP-SoxS activation with Sth1-dCas9, we constructed a J3-Sth1 promoter by modifying the J3 promoter to replace the *S. py* dCas9 AGG PAM with the Sth1-dCas9 TGAGAAT PAM at the J306 target site 81 nt upstream of the TSS.

For *Lachnospiraceae bacterium* dCas12a fused with AsiA_m2.1, termed dCas12a-AsiA in this work, we generated 170 nt tiling arrays with TTTV PAM sites every 20 nt with 5'-NNNNNNNNNNNNNNNNTTTV-3' as a 20 base spacer sequence for each sgRNA (each 20 base spacer is preceded by the TTTV PAM, so the preceding 20 base spacer ends with TTTV). We randomly generated 100,000 sequences of 170 nt and applied the same selection criteria as described above to generate the I1 sequence and I101-108 crRNAs. I1T and I1B and their +10 nt shifted promoters were constructed in a similar manner as in K1 promoter systems (Figure S14). For the I1T promoter, we also constructed +5 nt to +15 nt variants.

All tested Sth1-dCas9-AsiA and dCas12a-AsiA conditions exhibited no significant change in gene expression. Therefore, MCP-SoxS recruitment by Sth1-dCas9 was prioritized (Figure S15).

#### Supporting Figures

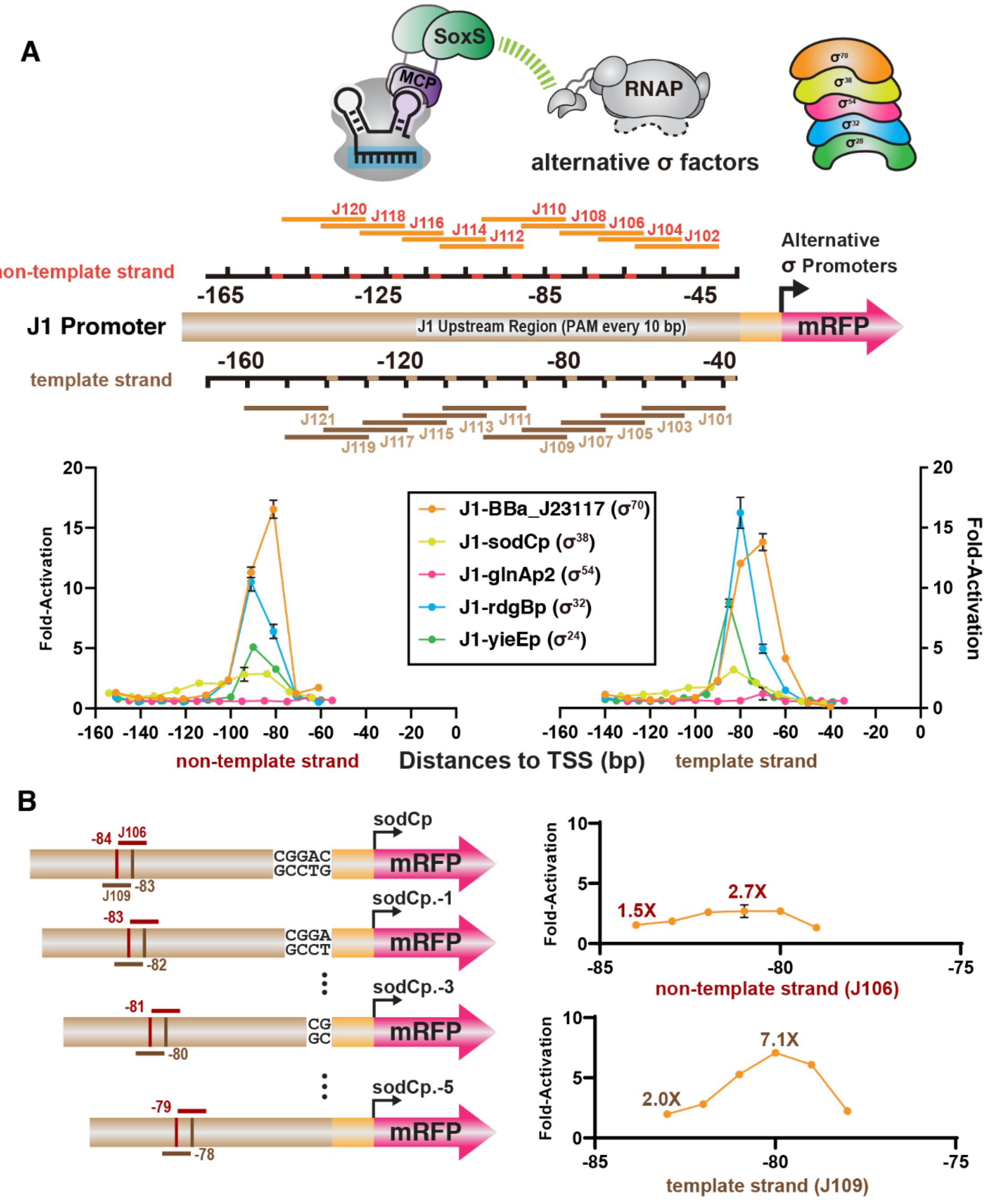


**Figure S1**. Target site screening for SoxS-mediated CRISPRa at alternative σ-factor promoters

**A)** Target site screening from -40 to -170 bp upstream of the TSS in the J1 reporter, following a previously-reported approach.[4] For most of the $\sigma^{70}$ family promoters, the optimal sites are in a narrow range 60-100 bp upstream of TSS. The $\sigma^{54}$ promoter, glnAp2, is the exception as it cannot be activated by SoxS. **B)** CRISPRa screening at phase-shifted sodCp ($\sigma^{54}$) promoters with 1 bp resolution. Shifting the target sites from -83 bp to -80 bp showed optimal CRISPR activity similar to the optimal phase of $\sigma^{70}$ promoters. Values in panel A and B represent the mean ± standard deviation calculated from n = 3.

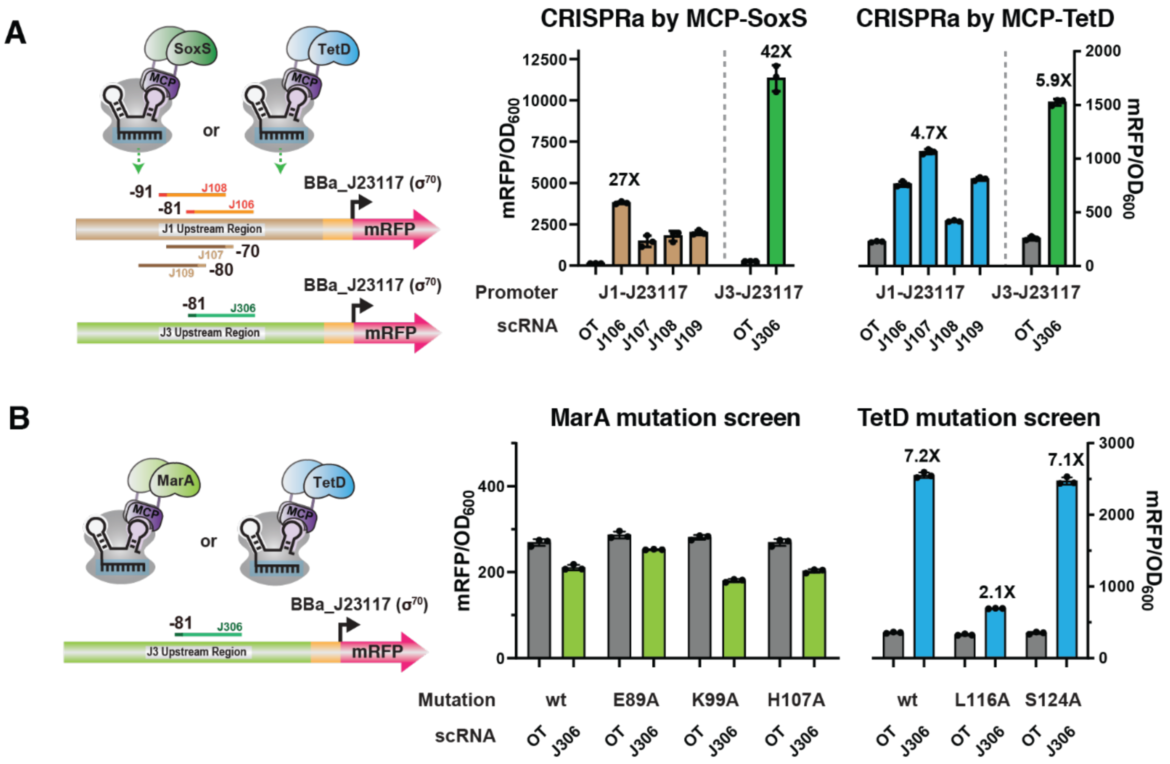


**Figure S2**. CRISPRa with MarA and TetD mutants

**A)** CRISPRa at J1 (J106-J109) and J3 (J306) reporters with MCP-SoxS and MCP-TetD. **B)** CRISPRa with MarA and TetD mutants at a J3 (J306) reporter. MarA mutants showed no improved activation beyond wild-type MarA and TetD mutants showed no improved activation beyond wild-type TetD. hAAVS1 scRNA was used as off-target scRNA in all cases. Values in panel A and B represent the mean ± standard deviation calculated from n = 3.

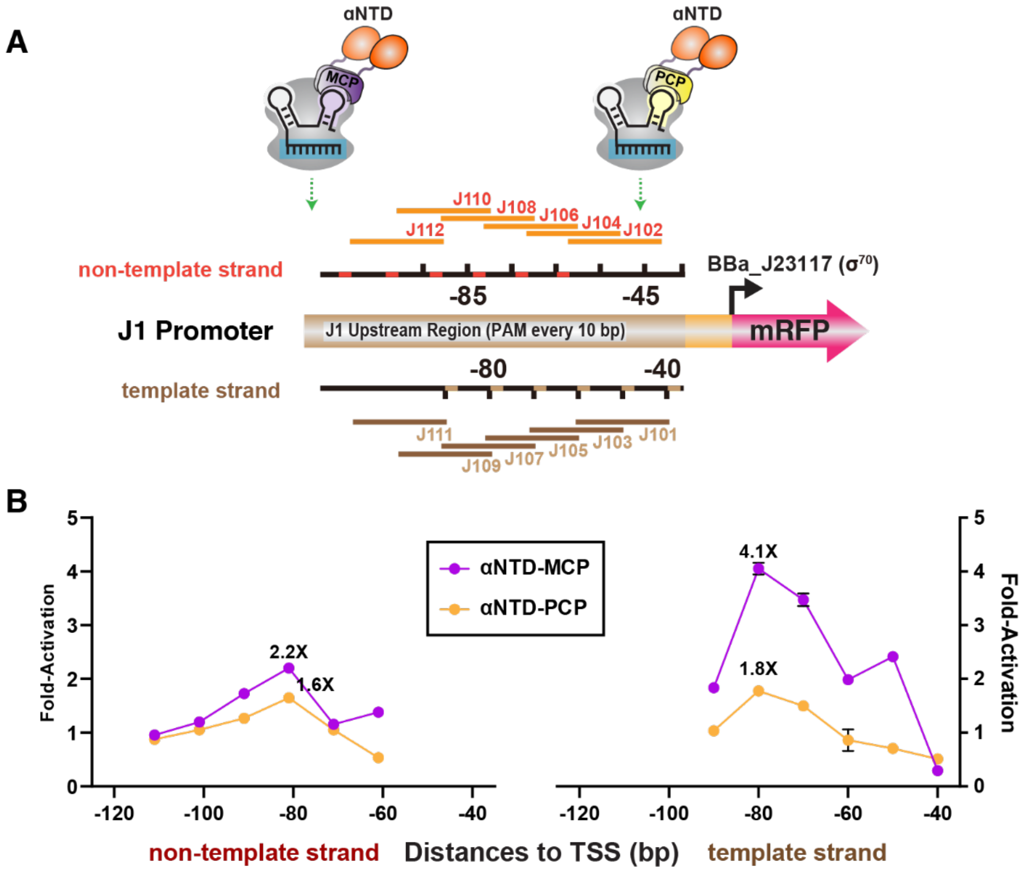


**Figure S3**. αNTD-mediated CRISPRa with RNA-binding protein recruitment

**A)** Conceptual depiction of CRISPRa screening at 40-111 bp (J101-J112) upstream of the TSS by αNTD-MCP and αNTD-PCP. **B)** CRISPRa screening showed that J109 (-80 on the template strand) is the optimal site for αNTD in both MCP- and PCP-based recruitment. Values in panel B represent the mean ± standard deviation calculated from n = 3.

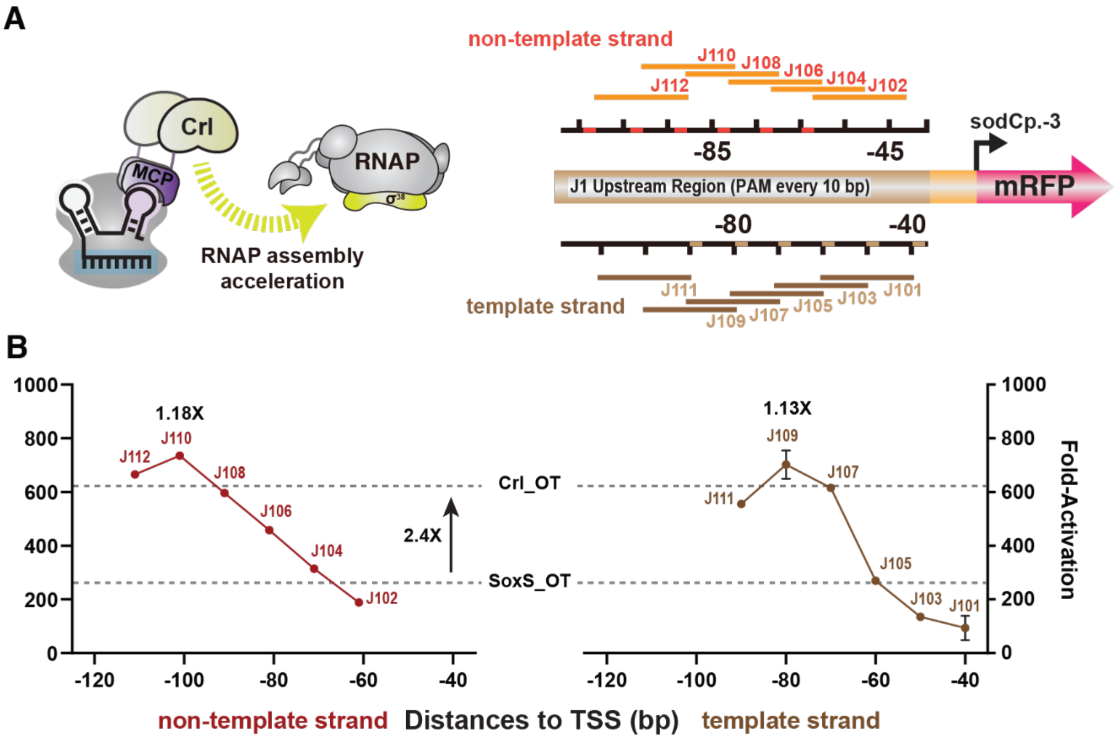


**Figure S4.** RNAP-σ[38] assembly using MCP-Crl

**A)** Schematic of CRISPRa screening at 40-111 bp (J101-J112) upstream of the TSS at the J1-sodCp.-3 (σ[38] promoter) by MCP-Crl. **B)** CRISPRa screening showed weak activity at J109 (1.13x) and J110 (1.18x) relative to the Crl off-target control (Crl_OT). The hAAVS1 sequence was used for the off-target scRNA. Significant non-specific gene activation could be observed by expressing MCP-Crl alone (2.4x) compared to MCP-SoxS expression (SoxS_OT). Since the *crl* gene is silent in *E. coli* K-12 MG1655 used in this study,[5] the presence of Crl alone may upregulate RNAP-σ[38] assembly without localization by CRISPR complex. Values in panel B represent the mean ± standard deviation calculated from n = 3.

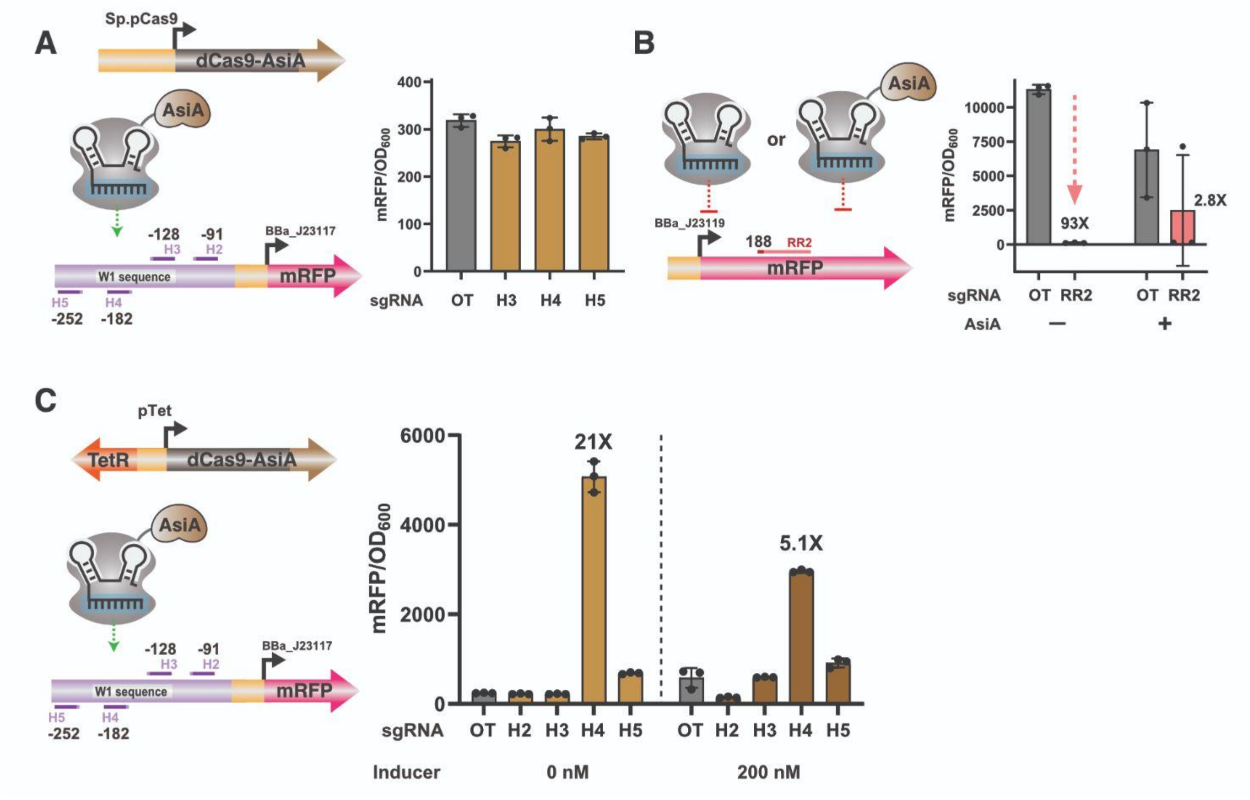


**Figure S5**. CRISPRa with dCas9-AsiA

**A)** CRISPRa screening at the W1 reporter with dCas9-AsiA when expressed under the constitutive Sp.pCas9 promoter showed no significant activation at target sites previously reported to be effective (H4 & H5 at -182 and -252 bp).[2] **B)** CRISPRi using an mRFP-targeting sgRNA (RR2) paired with either dCas9 or dCas9-AsiA showed variable and impaired activity with constitutively-expressed dCas9-AsiA, suggesting unstable expression. **C)** CRISPRa with dCas9-AsiA expressed from an aTc-inducible TetR-Ptet promoter showed 21-fold activation from H4 sgRNA even in an absence of inducer. Induction of dCas9-AsiA led to a 2.4 increase in basal expression (compare OT samples with and without inducer) and a decrease in CRISPRa-mediated expression level. The increase in basal expression may be due to a non-specific effect of dCas9-AsiA.[2] The decrease in CRISPRa-mediated gene expression is unexpected and could be a result of metabolic burden derived from expression of AsiA, as suggested previously.[1] See also Figure S11. Values in panel A, B, and C represent the mean ± standard deviation calculated from n = 3.

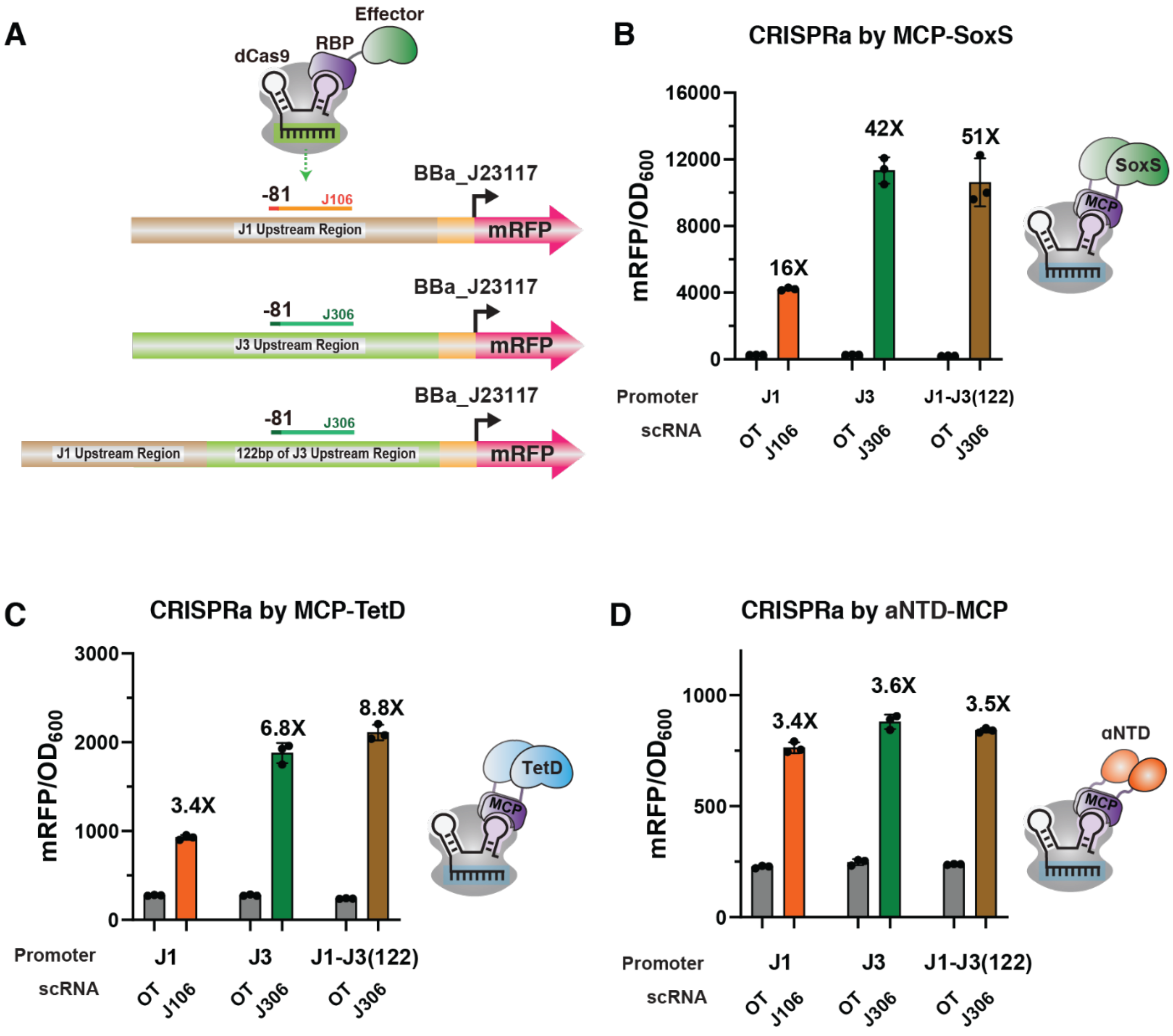


**Figure S6**. Comparison of different CRISPRa systems at J1, J3, and J1-J3(122) promoters

**A)** Schematic of the J1, J3, and J1-J3(122) promoters. **B)** MCP-SoxS activation at J3 showed improved activation with J3 and J1-J3(122) compared to J1. The same experiment was also performed for **C)** MCP-TetD and **D)** αNTD-MCP. MCP-SoxS and MCP-TetD exhibited higher CRISPRa efficiency with the J3 promoter while αNTD-MCP showed similar activity across all three promoters. Values in panel B, C, and D represent the mean ± standard deviation calculated from n = 3.

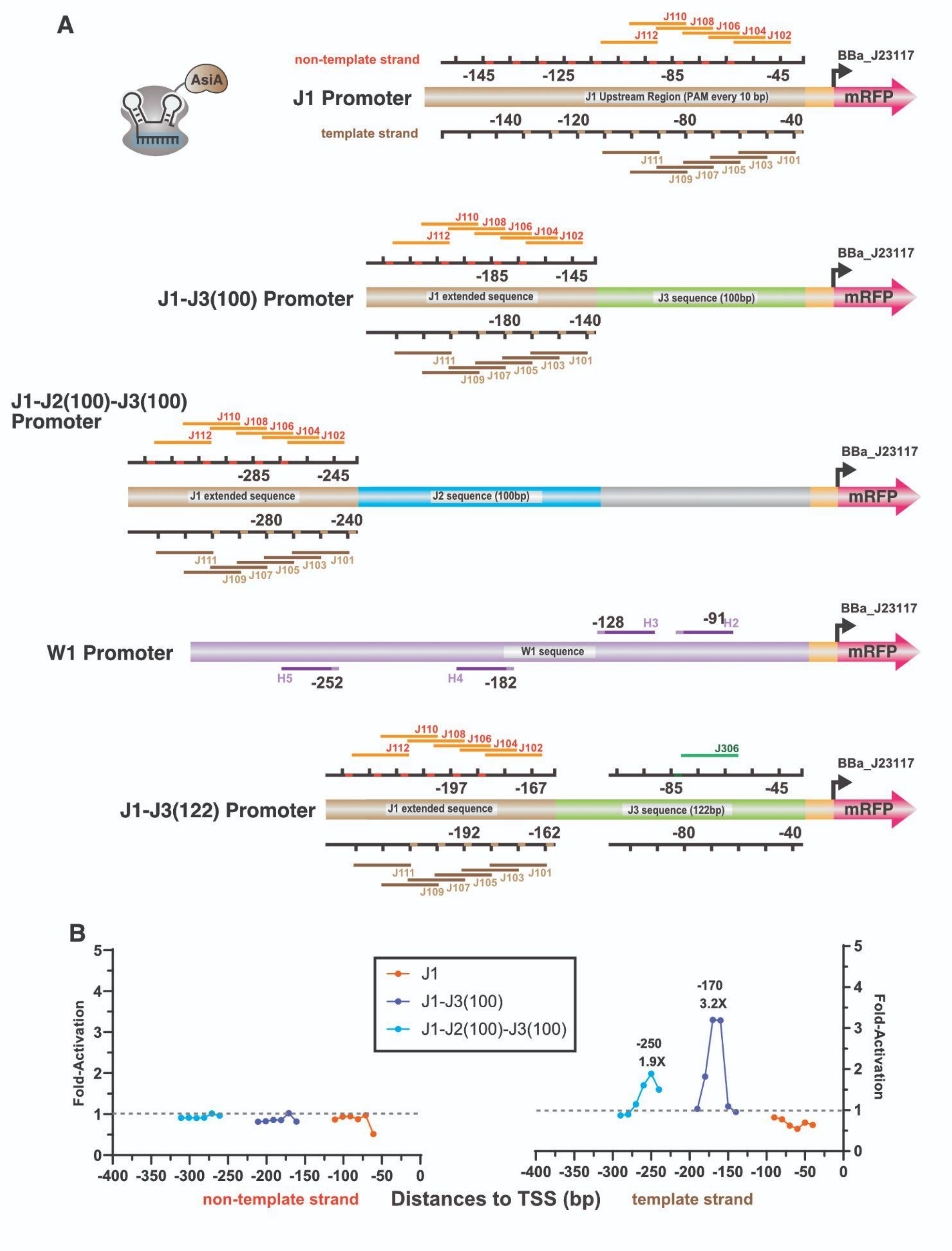
A
AsiA
non-template strand
template strand
J1 Promoter
J1 Upstream Region (PAM every 10 bp)
mRFP
BBa_J23117
-145
-125
-85
-45
-140
-120
-80
-40
J110 J108 J106 J104 J102
J112
J111 J109 J107 J105 J103 J101
J1-J3(100) Promoter
J1 extended sequence
J3 sequence (100bp)
-185
-180
J1-J2(100)-J3(100) Promoter
J2 sequence (100bp)
-285
-245
-280
-240
W1 Promoter
W1 sequence
-128 H3
-91 H2
H5 -252
H4 -182
J1-J3(122) Promoter
J3 sequence (122bp)
J306
-197
-167
-192
-162
B
Fold-Activation
J1
J1-J3(100)
J1-J2(100)-J3(100)
-170
3.2X
-250
1.9X
non-template strand
Distances to TSS (bp)
template strand

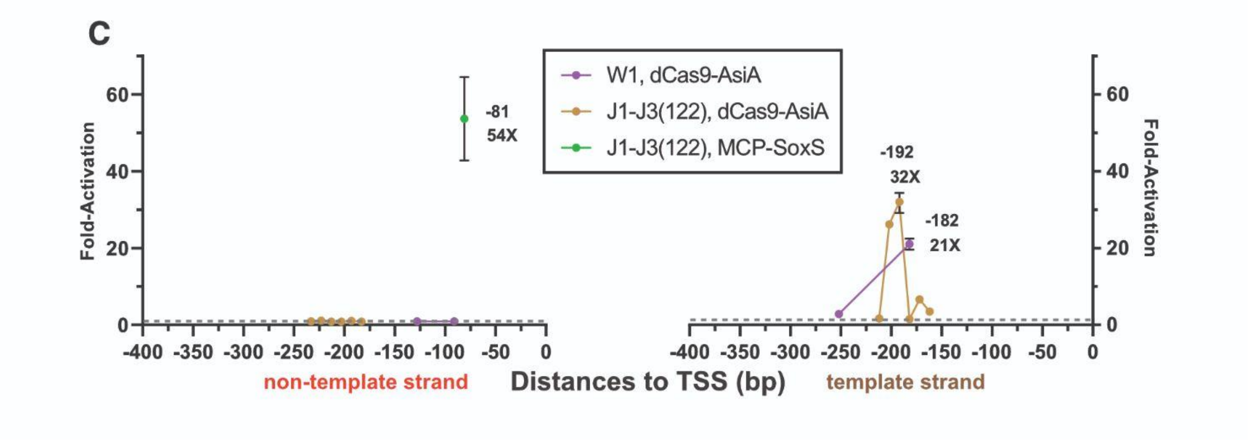


**Figure S7.** dCas9-AsiA based CRISPRa at target site positions in J1, J3, and J1-J3 promoters

**A)** Schematics showing promoters with arrays of sgRNA target sites at different distance ranges upstream of the TSS. The J1-J3(100), J1-J2(100)-J3(100), and J1-J3(122) promoters have 100, 200, and 122 bp added between the J1 array and the minimal promoter, respectively. For promoters with a J3 array, the J306 target site is always maintained at -81 bp for SoxS targeting. Target site positions for each promoter are listed in Table S4. **B)** dCas9-AsiA mediated activation produces relatively small effects at the J1-J3(100) promoter (1.8-fold to 3.2-fold activation between -160 to -180 bp) and the J1-J2(100)-J3(100) promoter (1.5-fold to 1.9-fold activation between -240 to -260 bp). **C)** dCas9-AsiA produces 32-fold activation at -192 bp (J107) in the phase-shifted J1-J3(122) promoter. There is no significant activation at -182 bp (J105) even though 21-fold activation was observed at the same distance with the W1 promoter and the H4 sgRNA (-182 bp). An activation value for dCas9/MCP-SoxS at -81 bp with the J1-J3(122) reporter is included for comparison. Values in panel B and C represent the mean ± standard deviation calculated from n = 3.

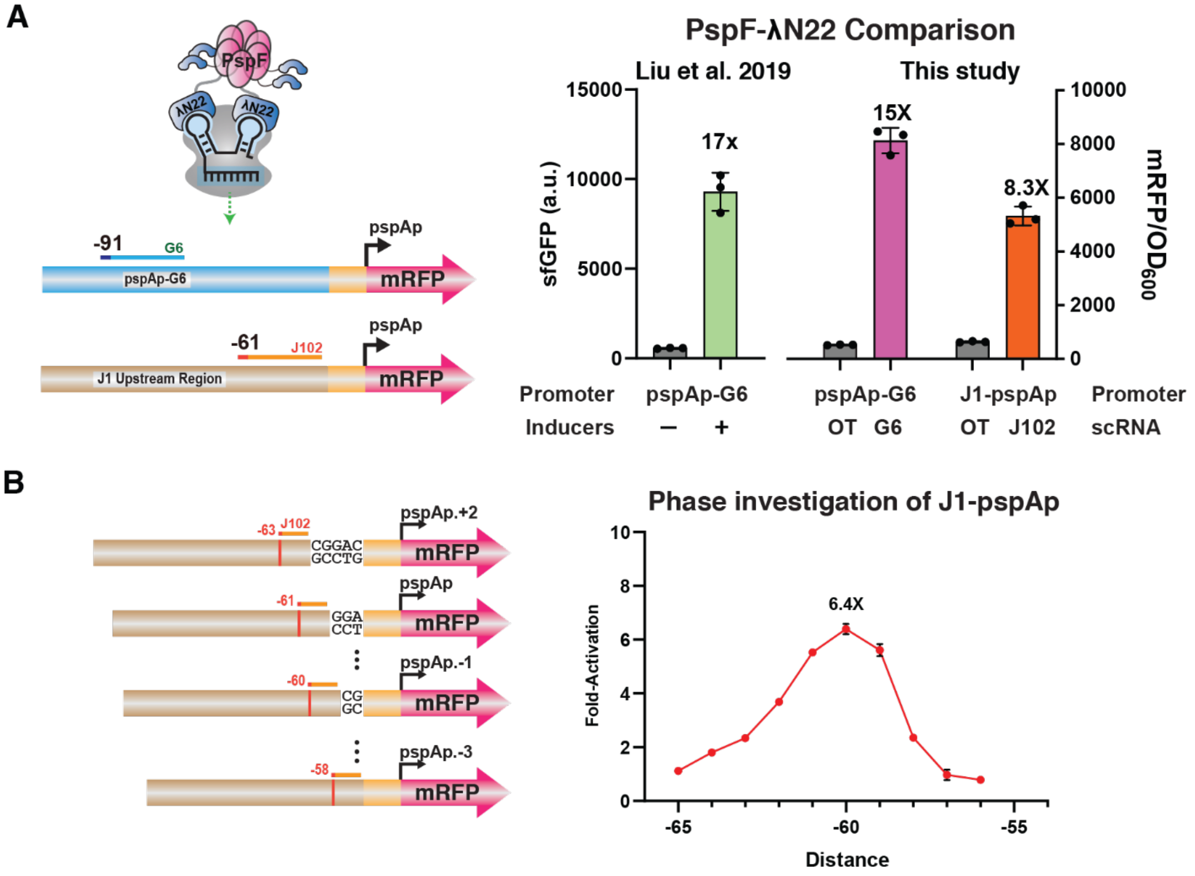


**Figure S8.** CRISPRa with PspF-λN22 at pspAp-G6 and J1 reporters

**A)** CRISPRa by PspF-λN22 at $\sigma^{54}$ promoters. At the pspAp-G6 promoter, we observe gene activation similar to the raw data reported previously.[6] The most effective activation observed from the J1-pspAp reporter is shown for comparison. See Fig 2 for complete target site characterization at J1. **B)** PspF CRISPRa with J1-pspAp shifted by 1 bp to determine target site preferences at single base resolution. In a side-by-side comparison, position -60 produced a small improvement compared to the original -61 target site in J1-pspAp. Values in panel A and B represent the mean ± standard deviation calculated from n = 3.

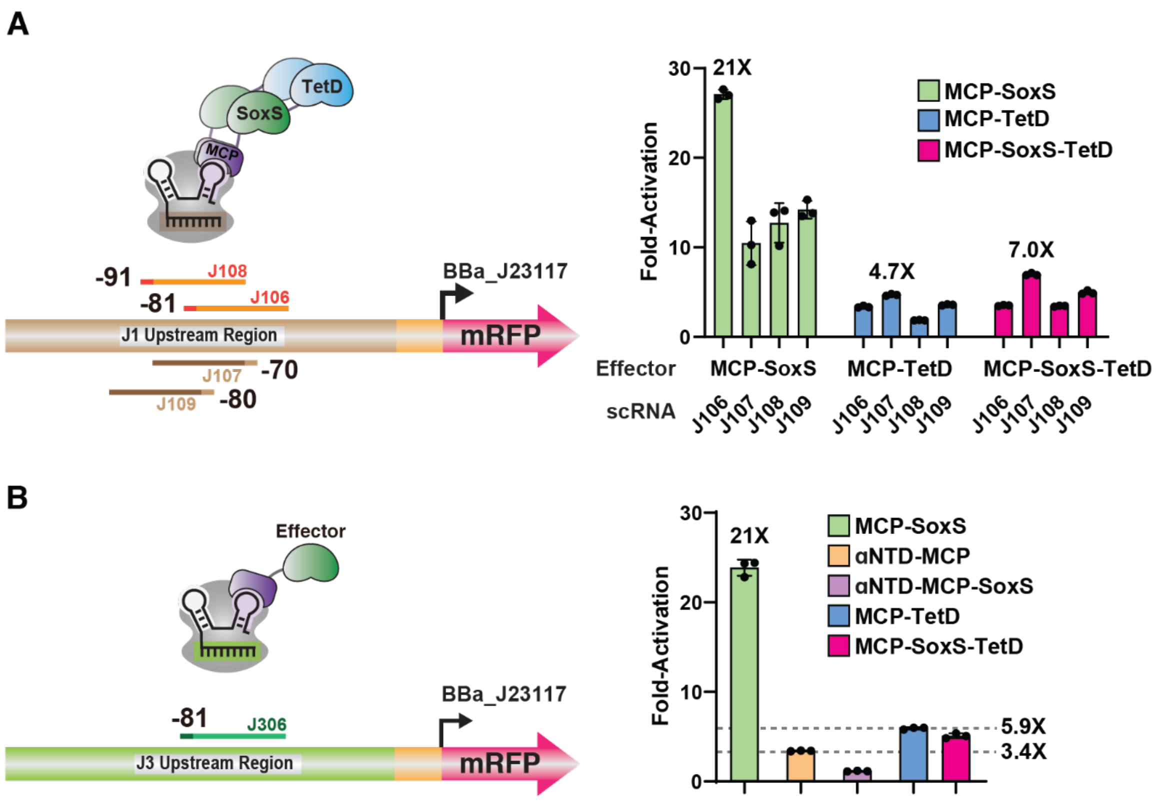


**Figure S9.** CRISPRa with tandem activator fusions

**A)** CRISPRa with MCP-SoxS-TetD compared to MCP-SoxS or MCP-TetD at 60-100 bp (J106-J109) in the J1 reporter. MCP-SoxS-TetD does not improve gene activation relative to MCP-SoxS. **B)** CRISPRa with MCP-SoxS-TetD or αNTD-MCP-SoxS at -81 bp (J306) in the J3 reporter. Neither fusion activator shows improved gene activation relative to MCP-SoxS. Values in panel A and B represent the mean ± standard deviation calculated from n = 3.

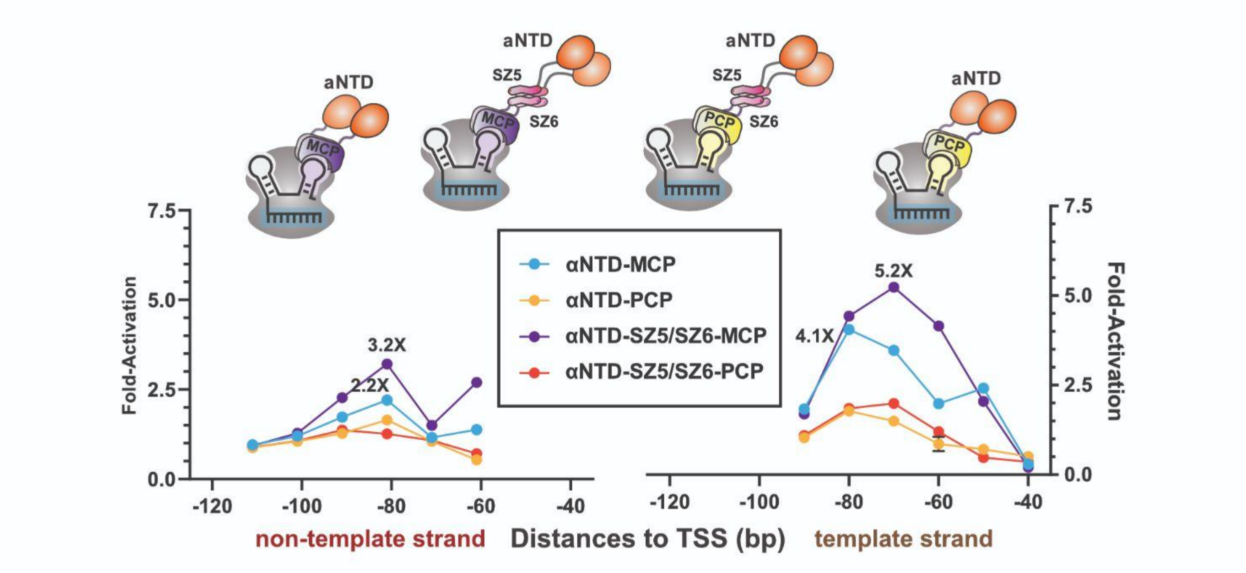


**Figure S10.** CRISPRa with αNTD using direct fusion or SYNZIP recruitment to RNA binding proteins

αNTD-mediated CRISPRa with direct fusion or SYNZIP recruitment to RNA binding proteins MCP or PCP. SYNZIP recruitment is performed with αNTD fused to SYNZIP5 (SZ5) and MCP or PCP fused to SYNZIP6 (SZ6). Gene activation was measured at multiple target sites in a J1 reporter with an appropriate scRNA with MS2 to bind MCP or PP7 to bind PCP. Some SYNZIP-mediated complexes show modest improvements compared to the corresponding direct fusion. In general, MCP-mediated recruitment outperforms PCP-mediated recruitment. Values represent the mean ± standard deviation calculated from n = 3.

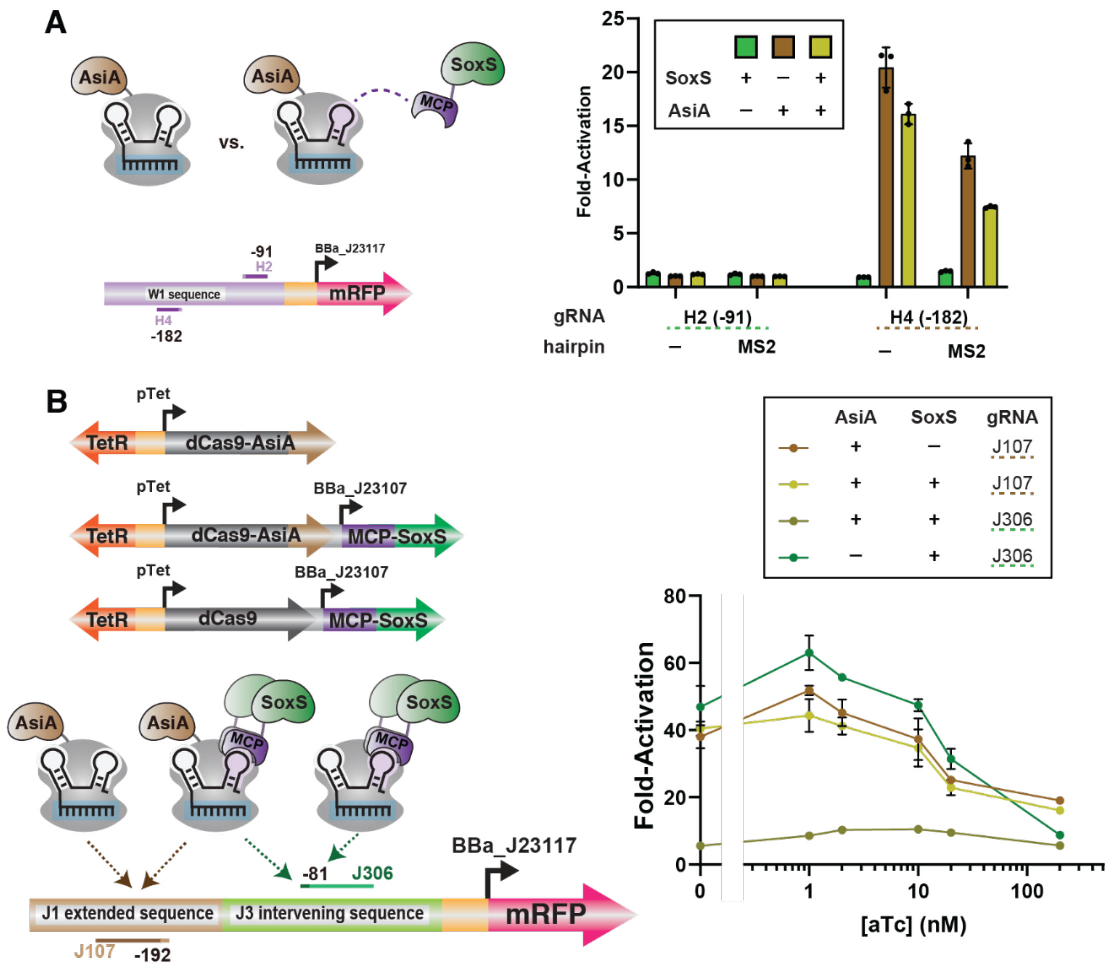


**Figure S11.** SoxS and AsiA CRISPRa exhibit antagonistic effects

**A)** dCas9-AsiA, dCas9/MCP-SoxS, or dCas9-AsiA/MCP-SoxS recruitment to target sites on the W1 promoter, using gRNAs with or without the MS2 hairpin. SoxS recruitment to the CRISPRa complex requires the MS2 hairpin. Only the H4 target site shows significant activation with dCas9-AsiA, and recruitment of SoxS to the dCas9-AsiA complex antagonizes gene activation, similar to effects observed with the J1-J3(122) promoter (Figure 4A). **B)** CRISPRa with dCas9 or dCas9-AsiA expressed from an aTc-inducible TetR-Ptet promoter. Gene activation is optimal at a low induction level with 1 nM aTc, and there is no induction level at which the dCas9-AsiA/MCP-SoxS complex outperforms dCas9-AsiA. Values in panel A and B represent the mean ± standard deviation calculated from n = 3.

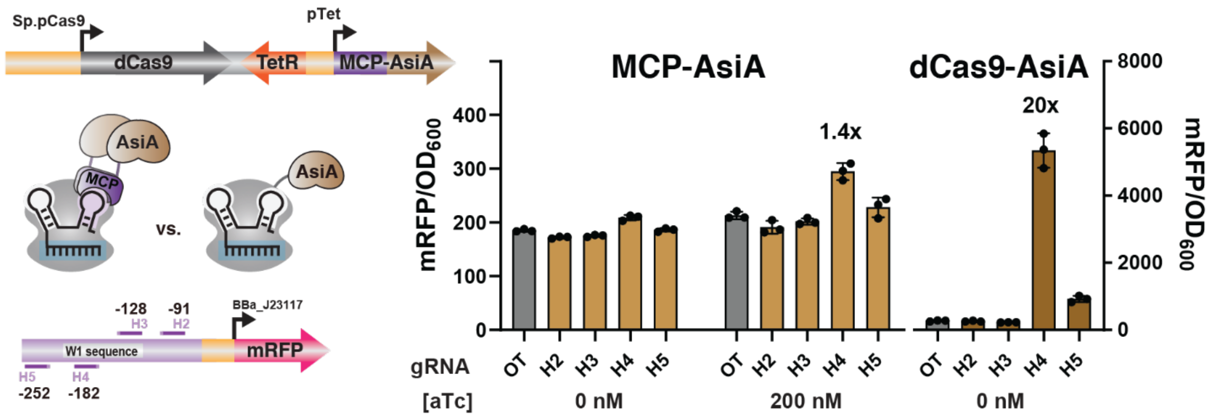


**Figure S12.** CRISPRa with an MS2 scRNA to recruit MCP-AsiA

Comparison of CRISPRa with dCas9-AsiA and dCas9/MCP-AsiA, all using the AsiA_m2.1 evolved variant. MCP-AsiA is expressed from an aTc-inducible promoter and tested at either 0 or 200 nM aTc. dCas9-AsiA is effective at 0 nM aTc (Figure S11). CRISPRa was evaluated at four target sites in the W1 promoter and compared to an off-target (OT) negative control. MCP-AsiA produces high basal activation in the OT control, and at most 1.4-fold activation at the H4 target site with 200 nM aTc. This effect is significantly smaller than the ~20-fold activation observed with dCas9-AsiA. Values represent the mean ± standard deviation calculated from n = 3.

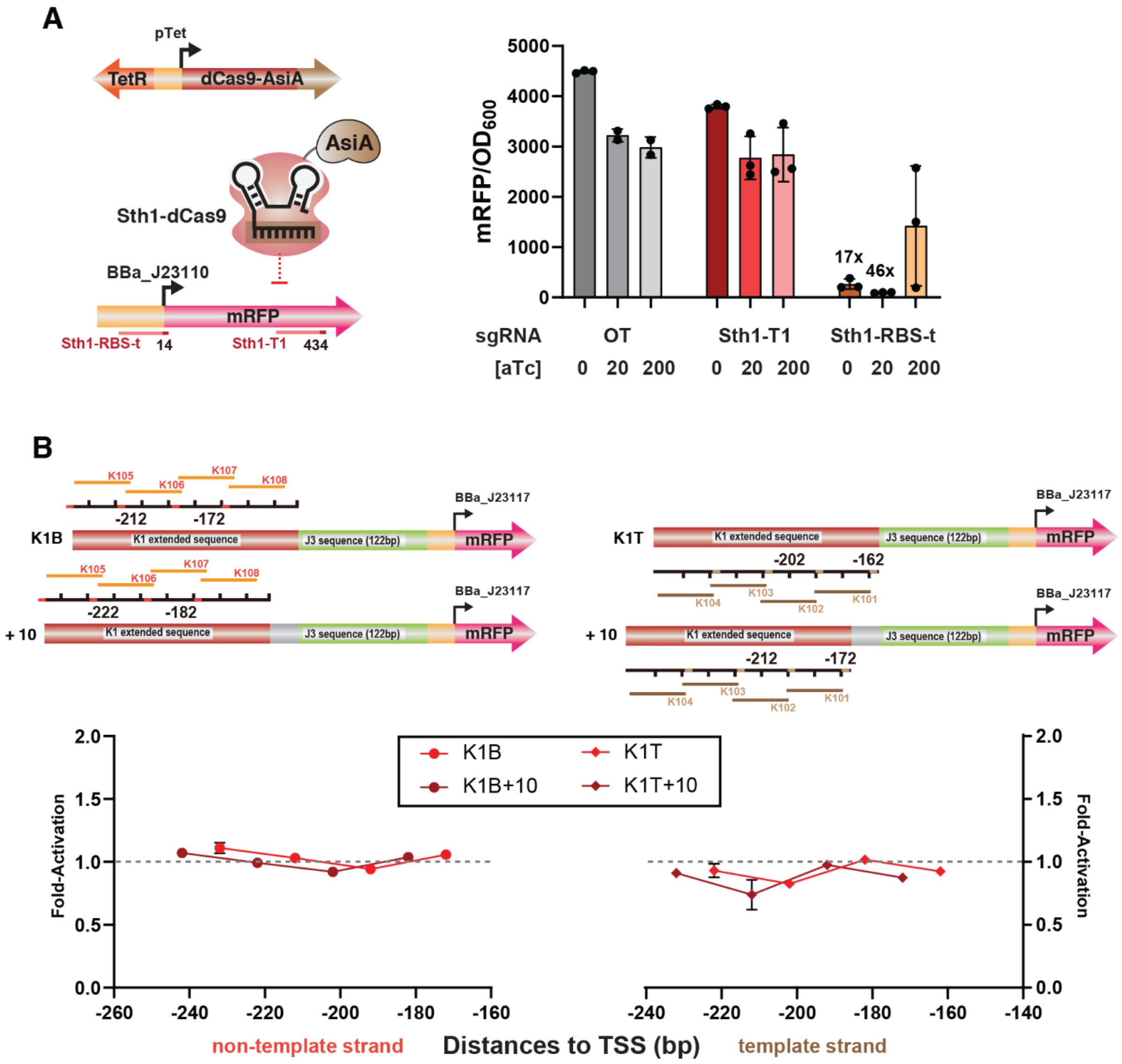


**Figure S13.** CRISPRa with Sth1-dCas9-AsiA is not effective

**A)** CRISPRi gene repression with Sth1-dCas9-AsiA to test expression and DNA-binding function. Sth1-dCas9-AsiA was targeted to target sites in the ribosome binding site (RBS-t) or within the open reading frame (Sth1-T1). Targeting RBS-t produced up to 46-fold repression upon induction with 20 nM aTc. However, 200 nM aTc produced variable CRISPRi effects across the replicates, which could indicate genetic instability. Similar variability in CRISPRi among the individual replicates was observed with Spy-dCas9-AsiA (Figure S5). **B)** CRISPRa with Sth1-dCas9-AsiA on K1 promoters with systematically tiled PAM sites positioned from 162-202 bp from the TSS. There is no significant CRISPRa effect across all tested target sites. Values in panel A and B represent the mean ± standard deviation calculated from n = 3.

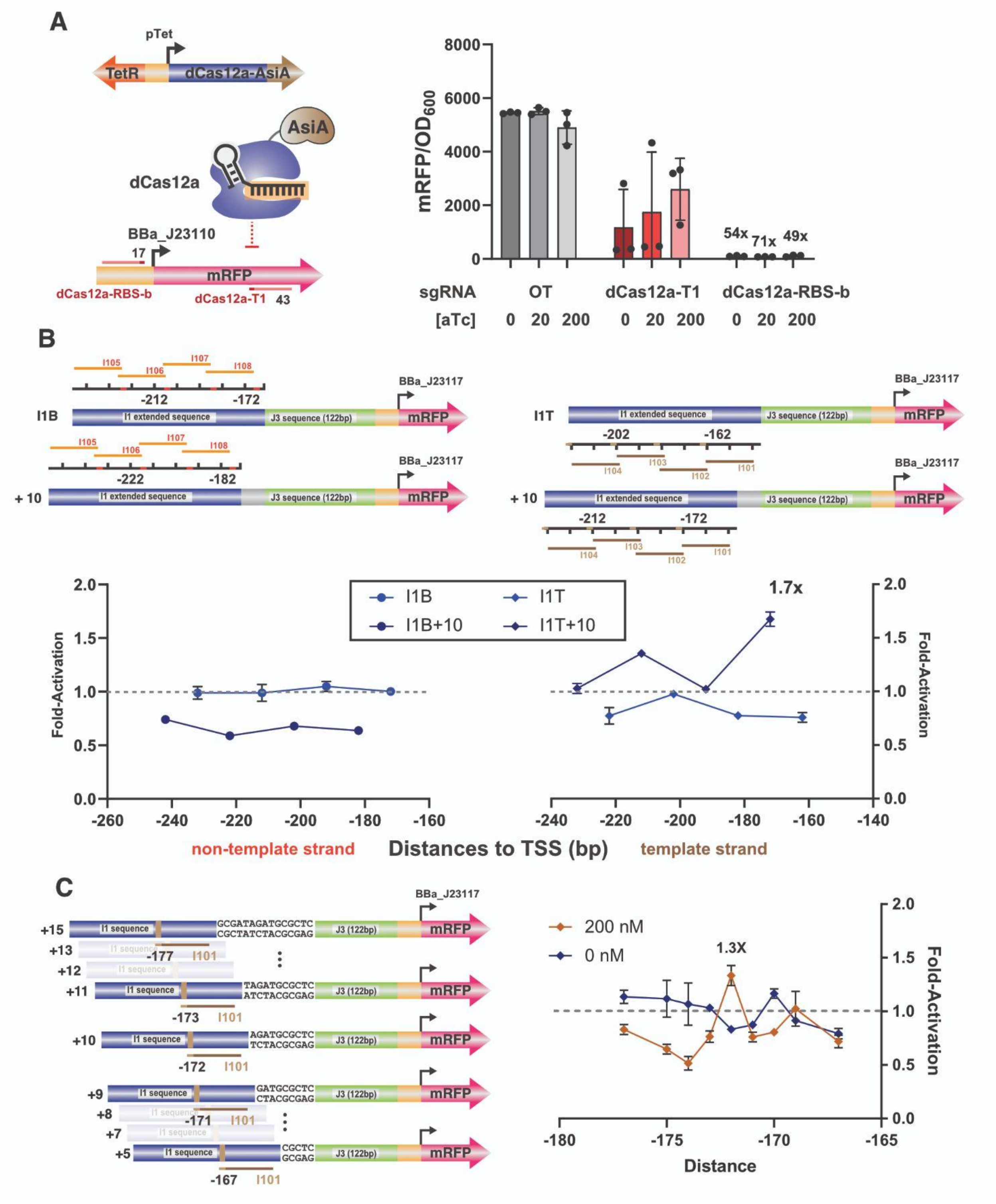


**Figure S14.** Modest CRISPRa with dCas12a-AsiA

**A)** CRISPRi gene repression with Lb-dCas12a-AsiA to test expression and DNA-binding function. Lb-dCas12a-AsiA was targeted to target sites in the ribosome binding site (RBS-b) or within the open reading frame (dCas12a-T1). Targeting RBS-b produced up to 71-fold repression. Targeting dCas12a-T1 also repressed gene expression to a lesser extent and with substantial variability among replicates (see also Figure S5 & S13). **B)** CRISPRa with Lb-dCas12a-AsiA on I1 promoters with systematically tiled PAM sites positioned from 162-202 bp from the TSS. We observed a small 1.7-fold increase in reporter gene expression at I101 crRNA on the I1T+10 reporter. C) CRISPRa with Lb-dCas12a-AsiA on I1 promoters with target sites shifted by 1-2 nt. In this experiment, 1.3-fold activation was observed. Values in panel A, B, and C represent the mean ± standard deviation calculated from n = 3.

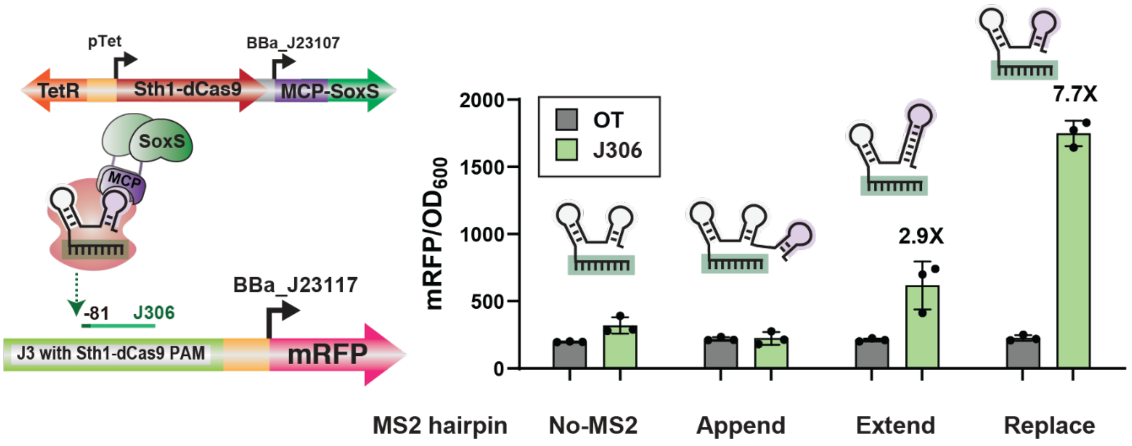


**Figure S15.** CRISPRa with Sth1-dCas9/MCP-SoxS and MS2 scRNAs.

Three alternative MS2 scRNA designs were screened for CRISPRa function with an mRFP reporter. MS2 was either appended to the 3' end of the Sth1 sgRNA, extended on the TrrnB terminator hairpin, or inserted to replace TrrnB. Complete scRNA sequences are included in the Supporting DNA sequences. Unmodified sgRNA lacking an MS2 hairpin was used as a negative control. The mRFP reporter has a modified J3 sequence with the Spy AGG PAM replaced by the Sth1 TGAGAAT PAM. Both the Extend and Replace scRNA designs produce significant CRISPRa effects, with the highest 7.7-fold effect from the Replace scRNA design. This optimal engineered Sth1 scRNA design is similar to our previously-reported optimal Spy scRNA.b2.[1,4] Values represent the mean ± standard deviation calculated from n = 3.

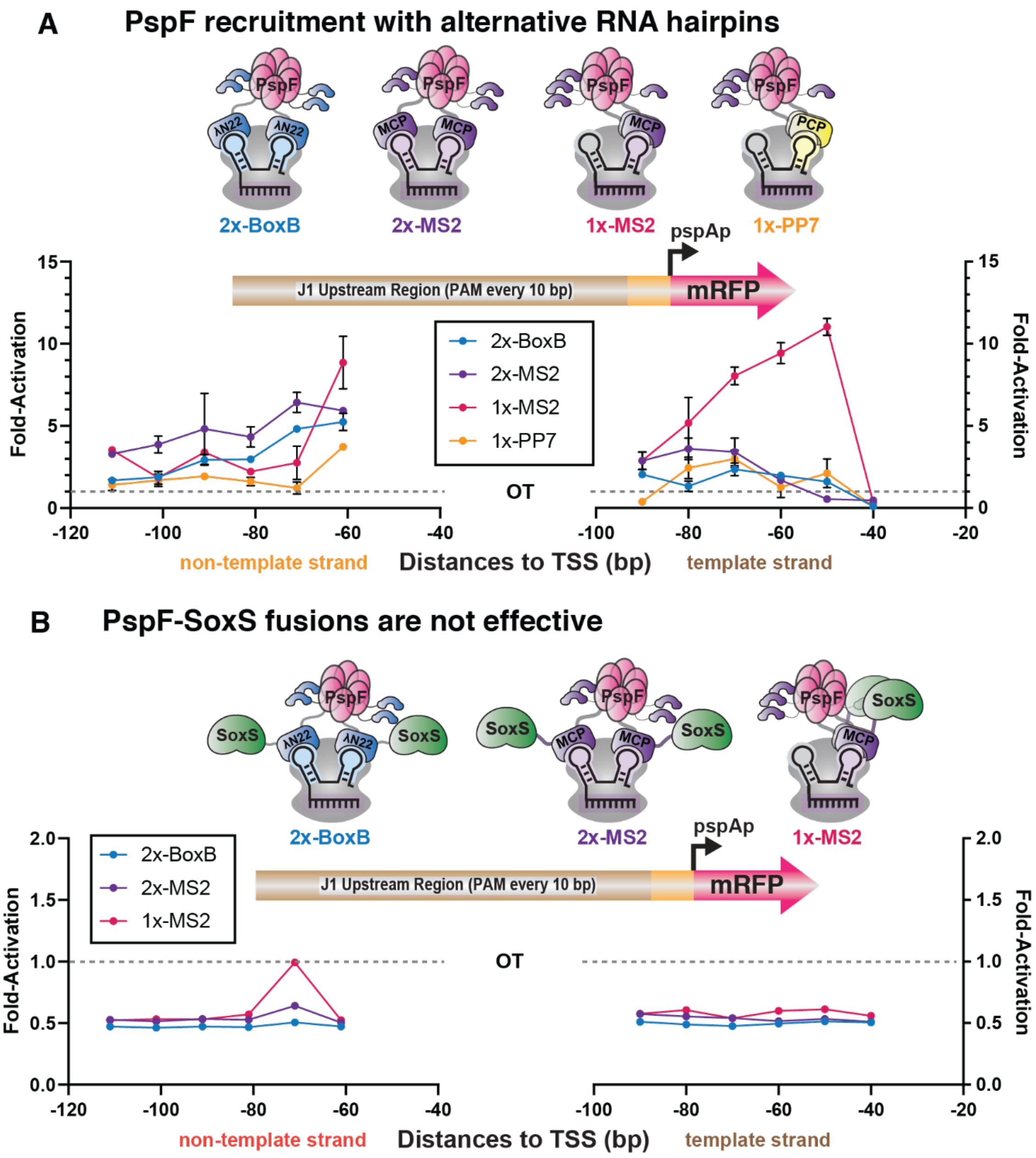


**Figure S16**. CRISPRa with PspF-SoxS fusions

**A)** CRISPRa with PspF recruited via scRNAs with alternative hairpins. 2x-BoxB, 2x-MS2, 1x-MS2, and 1x-PP7 scRNAs recruit PspF-λN22 (BoxB). PspF-MCP (MS2), or PspF-PCP (PP7). 1x-MS2 scRNA yielded the highest performance out of all conditions tested. **B)** CRISPRa with PspF and SoxS recruited together as a PspF-λN22-SoxS or PspF-MCP-SoxS fusion. These constructs were ineffective and exhibited small repression effects. Values in panel A and B represent the mean ± standard deviation calculated from n = 3.

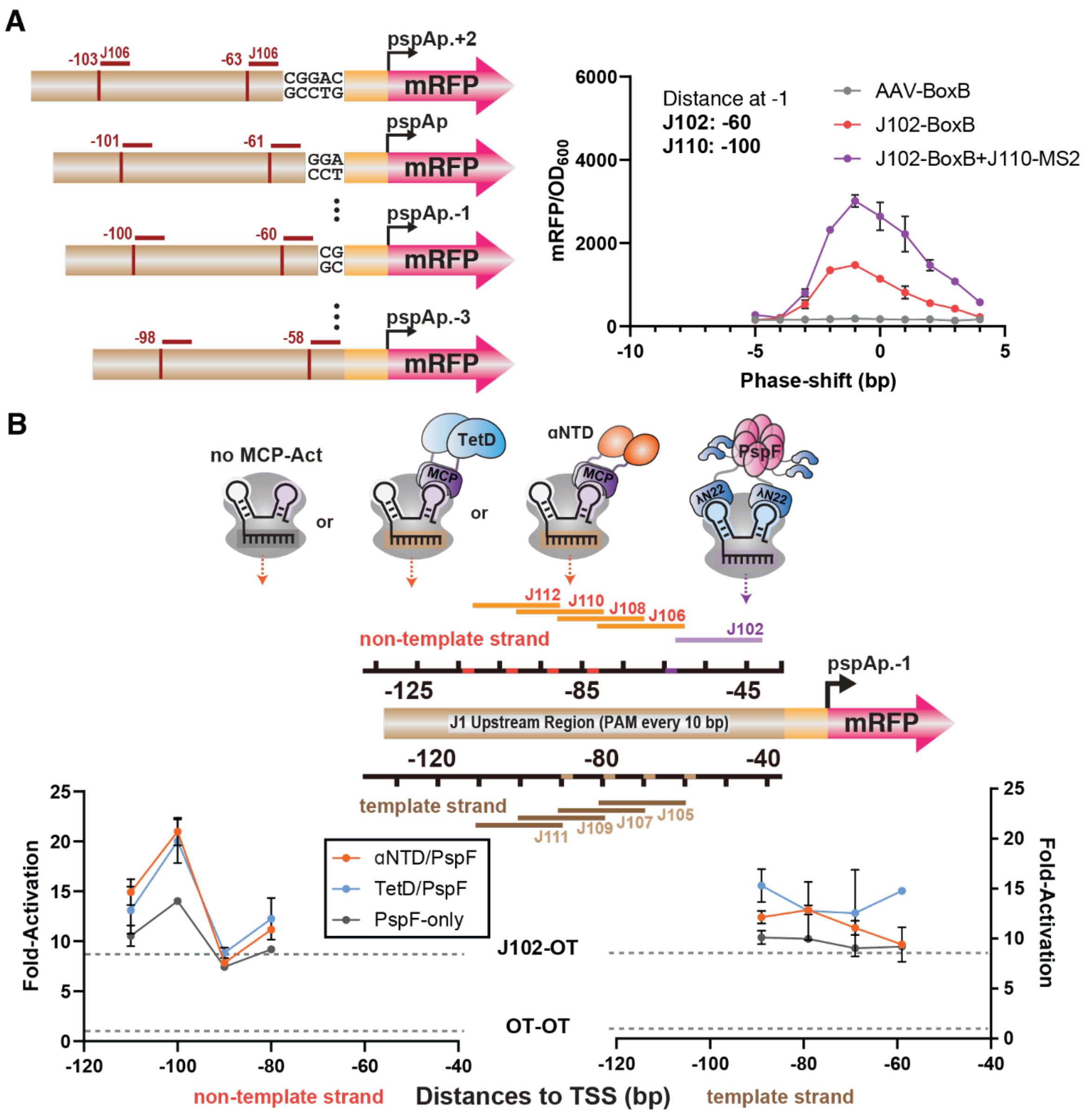


**Figure S17**. Characterization of PspF CRISPRa with alternative activators

**A**) Comparison of PspF-only and combined PspF/SoxS CRISPRa as a function of target site distance at 1 bp resolution, using reporters with different amounts of DNA inserted to shift target site positions. J102-BoxB scRNA recruits PspF-λN22 and J110-MS2 scRNA recruits MCP-SoxS. The optimal -1 position on the x-axis corresponds to J102-BoxB targeting position -60 and J110-MS2 targeting position -100. **B**) CRISPRa with αNTD+PspF, TetD+PspF, or PspF alone. The non-template targeting data is reproduced from Figure 5D for comparison to the template targeting data shown here. For simultaneous recruitment of PspF with αNTD or TetD, PspF was fixed at -60 on the non-template strand and the x-axis refers to the target site for αNTD or TetD. Values in panel A and B represent the mean ± standard deviation calculated from n = 3.

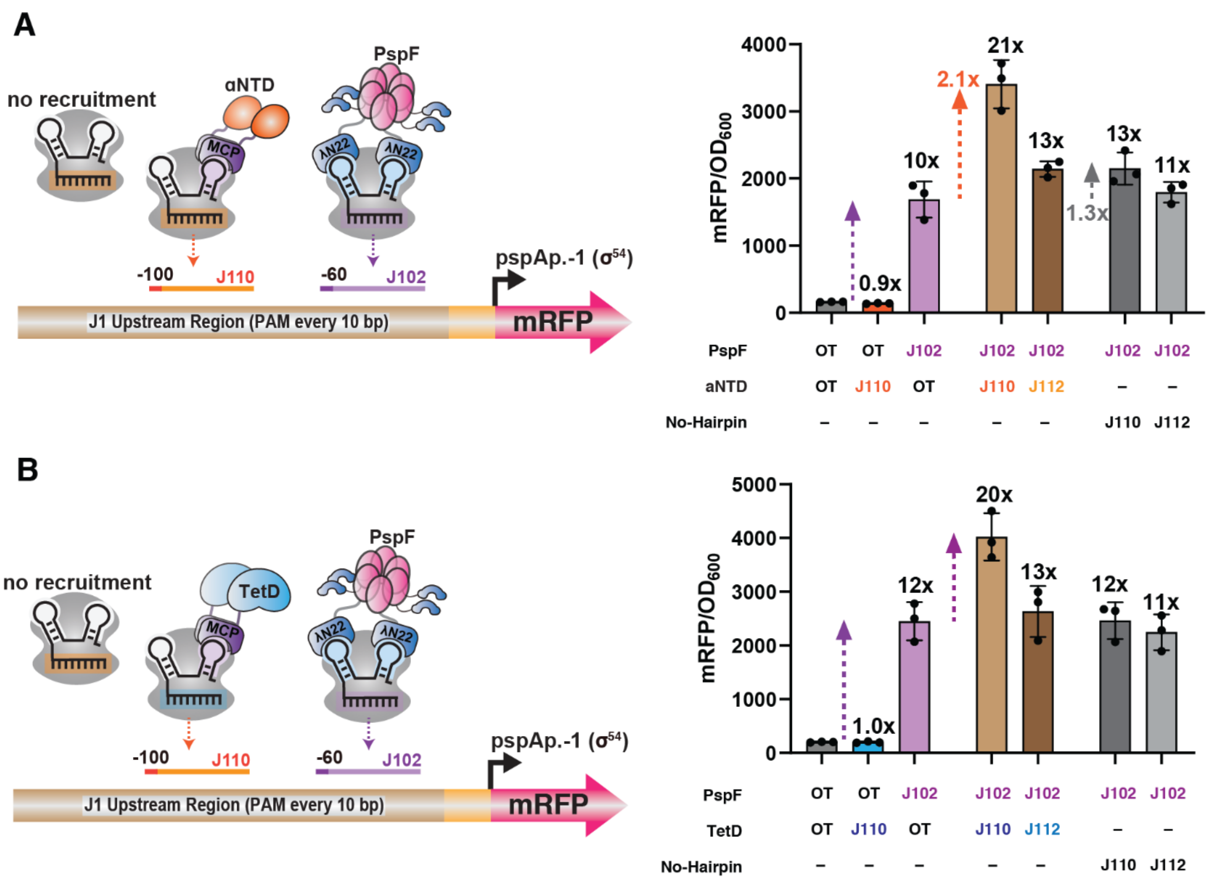


**Figure S18**. Characterization of CRISPRa with PspF+αNTD and PspF+TetD.

CRISPRa with PspF+αNTD **(A)** or PspF+TetD **(B)**. The observed effects are similar to PspF+SoxS (Figure 5). Values in panel A and B represent the mean ± standard deviation calculated from n = 3.

### Supporting Tables

**Table S1.** Bacterial strains used in this work

| Strain | Description | Genotype |
|---|---|---|
| MG1655 | Wildtype *Escherichia coli strain* | F- λ- ilvG- rfb-50 rph-1 |
| DH10B | Cloning strain | F- mcrA Δ(mrr-hsdRMS-mcrBC) φ80lacZΔM15 ΔlacX74 recA1 endA1 araD139 Δ (ara, leu)7697 galU galK λ- rpsL nupG /pMON14272 / pMON7124 |
| NEB turbo | Cloning strain | F' proA+B+ lacIq ΔlacZM15 / fhuA2 Δ(lac-proAB) glnV galK16 galE15 R(zgb-210::Tn10)TetS endA1 thi-1 Δ(hsdS-mcrB)5 |
| CD38[a] | MG1655 strain with integration of mRFP expressed under a strong constitutive promoter | MG1655_rbsAR::BBa_J23119-mRFP |

[a]CD38 was developed previously.[7]

**Table S2.** List of plasmids used in this work

| Plasmid | Marker | Origin | Promoter | Gene | Terminator | Reference |
|---|---|---|---|---|---|---|
| **dCas9 + Activator** | | | | | | |
| pCD442 | CmR | p15A | 1) Sp.pCas9<br>2) BBa_J23107 | 1) dCas9<br>2) MCP-SoxS[a] | 1) BBa_B0015<br>2) BBa_B1002 | Fontana and Dong *et al.* 2020[4] |
| pCD178 | CmR | p15A | 1) Sp.pCas9<br>2) BBa_J23107 | 1) dCas9<br>2) MCP-TetD | 1) BBa_B0015<br>2) BBa_B1002 | Dong *et al.* 2018[1] |
| pCD037 | CmR | p15A | 1) Sp.pCas9<br>2) BBa_J23107 | 1) dCas9<br>2) αNTD-MCP | 1) BBa_B0015<br>2) BBa_B1002 | Dong *et al.* 2018[1] |
| pCD196 | CmR | p15A | 1) Sp.pCas9<br>2) BBa_J23107 | 1) dCas9<br>2) αNTD-PCP | 1) BBa_B0015<br>2) BBa_B1002 | Dong *et al.* 2018[1] |
| pCK068 | CmR | p15A | 1) Sp.pCas9<br>2) BBa_J23107 | 1) dCas9<br>2) MCP-Crl | 1) BBa_B0015<br>2) BBa_B1002 | This work |
| pCK226 | CmR | p15A | 1) Sp.pCas9 | 1) dCas9-AsiA[b] | 1) BBa_B0015 | This work |
| pCK320 | CmR | p15A | 1) TetR-Ptet | 1) dCas9-AsiA[b] | 1) BBa_B0015 | This work |
| pAK007 | CmR | p15A | 1) TetR-Ptet<br>2) BBa_J23107 | 1) dCas9<br>2) MCP-SoxS | 1) BBa_B0015<br>2) BBa_B1002 | This work |
| pCK071 | CmR | p15A | 1) Sp.pCas9<br>2) BBa_J23107 | 1) dCas9<br>2) PspF-λN22 | 1) BBa_B0015<br>2) BBa_B1002 | This work |
| pCK694 | CmR | p15A | 1) Sp.pCas9<br>2) BBa_J23107 | 1) dCas9<br>2) PspF-MCP | 1) BBa_B0015<br>2) BBa_B1002 | This work |
| pCK934 | CmR | p15A | 1) Sp.pCas9<br>2) BBa_J23107 | 1) dCas9<br>2) PspF-PCP | 1) BBa_B0015<br>2) BBa_B1002 | This work |
| pCK157 | CmR | p15A | 1) Sp.pCas9<br>2) BBa_J23107<br>3) BBa_J23107 | 1) dCas9<br>2) MCP-SYNZIP6<br>3) αNTD-SYNZIP5 | 1) BBa_B0015<br>2) BBa_B1002<br>3) BBa_B1002 | This work |
| pCK770 | CmR | p15A | 1) Sp.pCas9<br>2) BBa_J23107<br>3) BBa_J23107 | 1) dCas9<br>2) PCP-SYNZIP6<br>3) αNTD-SYNZIP5 | 1) BBa_B0015<br>2) BBa_B1002<br>3) BBa_B1002 | This work |
| pAK010 | CmR | p15A | 1) Sp.pCas9<br>2) TetR-Ptet | 1) dCas9<br>2) MCP-AsiA | 1) BBa_B0015<br>2) BBa_B1002 | This work |
| pAK140 | CmR | p15A | 1) TetR-Ptet<br>2) BBa_J23107 | 1) Sth1-dCas9<br>2) MCP-SoxS | 1) BBa_B0015<br>2) BBa_B1002 | This work |
| pCK1004 | CmR | p15A | 1) TetR-Ptet | 1) Sth1-dCas9-AsiA | 1) BBa_B0015 | This work |
| pCK1008 | CmR | p15A | 1) TetR-Ptet | 1) Lb-dCas12a-AsiA | 1) BBa_B0015 | This work |

| Plasmid | Marker | Origin | Promoter | Gene | Terminator | Reference |
|---|---|---|---|---|---|---|
| **dCas9 + Activator + gRNA** | | | | | | |
| pCK005.X | CmR | p15A | 1) Sp.pCas9<br>2) BBa_J23107<br>3) BBa_J23119 | 1) dCas9<br>2) MCP-SoxS<br>3) scRNA | 1) BBa_B0015<br>2) BBa_B1002<br>3) TrrnB | Fontana and Dong *et al.* 2020[4] |
| pCD296.X | CmR | p15A | 1) Sp.pCas9<br>2) BBa_J23107<br>3) BBa_J23119 | 1) dCas9<br>2) MCP-TetD<br>3) scRNA | 1) BBa_B0015<br>2) BBa_B1002<br>3) TrrnB | Fontana and Dong *et al.* 2020[4] |
| pCK090.X | CmR | p15A | 1) Sp.pCas9<br>2) BBa_J23107<br>3) BBa_J23119 | 1) dCas9<br>2) MCP-MarA[c]<br>3) scRNA | 1) BBa_B0015<br>2) BBa_B1002<br>3) TrrnB | This work |
| pCK094.X | CmR | p15A | 1) Sp.pCas9<br>2) BBa_J23107<br>3) BBa_J23119 | 1) dCas9<br>2) MCP-TetD[c]<br>3) scRNA | 1) BBa_B0015<br>2) BBa_B1002<br>3) TrrnB | This work |
| pCD297.X | CmR | p15A | 1) Sp.pCas9<br>2) BBa_J23107<br>3) BBa_J23119 | 1) dCas9<br>2) αNTD-MCP<br>3) scRNA | 1) BBa_B0015<br>2) BBa_B1002<br>3) TrrnB | Fontana and Dong *et al.* 2020[4] |
| pRC099.X | CmR | p15A | 1) Sp.pCas9<br>2) BBa_J23107<br>3) BBa_J23119 | 1) dCas9<br>2) αNTD-PCP<br>3) scRNA | 1) BBa_B0015<br>2) BBa_B1002<br>3) TrrnB | This work |
| pCK147.X | CmR | p15A | 1) Sp.pCas9<br>2) BBa_J23107<br>3) BBa_J23119 | 1) dCas9<br>2) PspF-λN22<br>3) scRNA | 1) BBa_B0015<br>2) BBa_B1002<br>3) TrrnB | This work |
| pCK083.X | CmR | p15A | 1) Sp.pCas9<br>2) BBa_J23107<br>3) BBa_J23119 | 1) dCas9<br>2) MCP-Crl<br>3) scRNA | 1) BBa_B0015<br>2) BBa_B1002<br>3) TrrnB | This work |
| pCK309.X | CmR | p15A | 1) Sp.pCas9<br>2) BBa_J23119 | 1) dCas9-AsiA<br>2) scRNA | 1) BBa_B0015<br>2) TrrnB | This work |
| **Multiple activators** | | | | | | |
| pCK692 | CmR | p15A | 1) Sp.pCas9<br>2) BBa_J23107 | 1) dCas9<br>2) αNTD-MCP-SoxS | 1) BBa_B0015<br>2) BBa_B1002 | This work |
| pCK693 | CmR | p15A | 1) Sp.pCas9<br>2) BBa_J23107 | 1) dCas9<br>2) MCP-SoxS-TetD | 1) BBa_B0015<br>2) BBa_B1002 | This work |
| pCK321 | CmR | p15A | 1) TetR-Ptet<br>2) BBa_J23107 | 1) dCas9-AsiA[b]<br>2) MCP-SoxS | 1) BBa_B0015<br>2) BBa_B1002 | This work |
| pKO025 | CmR | p15A | 1) TetR-Ptet<br>2) Ptet<br>3) BBa_J23107 | 1) dCas9-AsiA<br>2) Sth1-dCas9<br>3) MCP-SoxS | 1) BBa_B0015<br>2) BBa_B0015<br>3) BBa_B1002 | This work |
| pCK660 | CmR | p15A | 1) Sp.pCas9<br>2) BBa_J23107 | 1) dCas9<br>2) MCP-SoxS | 1) BBa_B0015<br>2) BBa_B1002 | This work |

| Plasmid | Marker | Origin | Promoter | Gene | Terminator | Reference |
|---|---|---|---|---|---|---|
| | | | 3) BBa_J23107 | 3) PspF-λN22 | 3) BBa_B1002 | |
| pCK761 | CmR | p15A | 1) Sp.pCas9<br>2) BBa_J23107<br>3) BBa_J23107 | 1) dCas9<br>2) αNTD-MCP<br>3) PspF-λN22 | 1) BBa_B0015<br>2) BBa_B1002<br>3) BBa_B1002 | This work |
| pCK948 | CmR | p15A | 1) Sp.pCas9<br>2) BBa_J23107<br>3) BBa_J23107 | 1) dCas9<br>2) MCP-TetD<br>3) PspF-λN22 | 1) BBa_B0015<br>2) BBa_B1002<br>3) BBa_B1002 | This work |
| pCK882 | CmR | p15A | 1) Sp.pCas9<br>2) BBa_J23107 | 1) dCas9<br>2) MCP-SoxS-SYNZIP6 | 1) BBa_B0015<br>2) BBa_B1002 | This work |
| pCK883 | CmR | p15A | 1) Sp.pCas9<br>2) BBa_J23107<br>3) BBa_J23107 | 1) dCas9<br>2) MCP-SoxS-SYNZIP6<br>3) αNTD-SYNZIP5 | 1) BBa_B0015<br>2) BBa_B1002<br>3) BBa_B1002 | This work |
| pCK695 | CmR | p15A | 1) Sp.pCas9<br>2) BBa_J23107 | 1) dCas9<br>2) PspF-MCP-SoxS | 1) BBa_B0015<br>2) BBa_B1002 | This work |
| pCK696 | CmR | p15A | 1) Sp.pCas9<br>2) BBa_J23107 | 1) dCas9<br>2) PspF-λN22-SoxS | 1) BBa_B0015<br>2) BBa_B1002 | This work |
| **Reporter** | | | | | | |
| pJF076 | AmpR | pSC101** | J1-BBa_J23117 | mRFP | BBa_B0015 | Dong *et al.* 2018[1] |
| pJF143.J3 | AmpR | pSC101** | J3-BBa_J23117 | mRFP | BBa_B0015 | Fontana and Dong *et al.* 2020[4] |
| pCK001 | AmpR | pSC101** | J1-sodCp | mRFP | BBa_B0015 | Fontana and Dong *et al.* 2020[4] |
| pCK002 | AmpR | pSC101** | J1-glnAp2 | mRFP | BBa_B0015 | Fontana and Dong *et al.* 2020[4] |
| pCK003 | AmpR | pSC101** | J1-rdgBp | mRFP | BBa_B0015 | Fontana and Dong *et al.* 2020[4] |
| pCK004 | AmpR | pSC101** | J1-yieEp | mRFP | BBa_B0015 | Fontana and Dong *et al.* 2020[4] |
| pCK067.N | AmpR | pSC101** | J1-sodCp_phase-shift | mRFP | BBa_B0015 | Fontana and Dong *et al.* 2020[4] |

| Plasmid | Marker | Origin | Promoter | Gene | Terminator | Reference |
| --- | --- | --- | --- | --- | --- | --- |
| pCD249 | AmpR | pSC101** | W1-BBa_J23117 | mRFP | BBa_B0015 | Dong *et al.* 2018[1] |
| pCK702 | AmpR | pSC101** | J1-J3(100) -BBa_J23117 | mRFP | BBa_B0015 | This work |
| pCK703 | AmpR | pSC101** | J1-J2(100)-J3(100) -BBa_J23117 | mRFP | BBa_B0015 | This work |
| pAK011 | AmpR | pSC101** | J1-J3(122) -BBa_J23117 | mRFP | BBa_B0015 | This work |
| pCK070 | AmpR | pSC101** | J1-pspAp | mRFP | BBa_B0015 | This work |
| pCK069 | AmpR | pSC101** | pspAp-G6 | mRFP | BBa_B0015 | This work |
| pCK662.N | AmpR | pSC101** | J1-pspAp_phase-shift | mRFP | BBa_B0015 | This work |
| pKO021 | AmpR | pSC101** | J3-BBa_J23117 with Sth1 PAM | mRFP | BBa_B0015 | This work |
| pCK1006.N | AmpR | pSC101** | K1T-BBa_J23117 | mRFP | BBa_B0015 | This work |
| pCK1007.N | AmpR | pSC101** | K1B-BBa_J23117 | mRFP | BBa_B0015 | This work |
| pCK1011.N | AmpR | pSC101** | I1T-BBa_J23117 | mRFP | BBa_B0015 | This work |
| pCK1012.N | AmpR | pSC101** | I1B-BBa_J23117 | mRFP | BBa_B0015 | This work |
| pKO023 | AmpR | pSC101** | J1-J3(122)-BBa_J23117 with Sth1 PAM | mRFP | BBa_B0015 | This work |
| **Reporter + gRNA** | | | | | | |
| pCK161.X | AmpR | pSC101** | 1) BBa_J23119 2) J1-BBa_J23117 | 1) scRNA 2) mRFP | 1) TrrnB 2) BBa_B0015 | This work |
| pCK101.X | AmpR | pSC101** | 1) BBa_J23119 2) J3-BBa_J23117 | 1) scRNA 2) mRFP | 1) TrrnB 2) BBa_B0015 | This work |
| **gRNA(s)** | | | | | | |
| pCK303.scRNA | KmR | ColE1 | BBa_J23119 | scRNA_b2.MS2 | TrrnB | This work |
| pAK030.sgRNA | KmR | ColE1 | BBa_J23119 | sgRNA | TrrnB | This work |

| Plasmid | Marker | Origin | Promoter | Gene | Terminator | Reference |
|---|---|---|---|---|---|---|
| pCK676. scRNA | KmR | ColE1 | BBa_J23119 | scRNA_2xBoxB | TrrnB | This work |
| pCK732. scRNA | KmR | ColE1 | BBa_J23119 | scRNA_b1.PP7 (1xPP7) | TrrnB | This work |
| pCK697. scRNA | KmR | ColE1 | BBa_J23119 | scRNA_2xMS2 | TrrnB | This work |
| pKO018. scRNA | KmR | ColE1 | BBa_J23119 | Sth1-dCas9 scRNA with MS2 | TrrnB | This work |
| pCK1005. sgRNA | KmR | ColE1 | BBa_J23119 | Sth1-dCas9 sgRNA | TrrnB | This work |
| pCK1010. crRNA | KmR | ColE1 | BBa_J23119 | Lb-dCas12a crRNA | TrrnB | This work |
| pKO024. scRNAs | KmR | ColE1 | 1) BBa_J23119<br>2) BBa_J23119 | 1) Sth1-dCas9 scRNA with MS2.X<br>2) Spy-dCas9 sgRNA.X | 1) TrrnB<br>2) TrrnB | This work |
| pCK677. scRNAs | KmR | ColE1 | 1) BBa_J23119<br>2) BBa_J23119 | 1) scRNA_2xBoxB_J102<br>2) scRNA_b2.MS2.X | 1) TrrnB<br>2) TrrnB | This work |
| pCK962. gRNAs | KmR | ColE1 | 1) BBa_J23119<br>2) BBa_J23119 | 1) scRNA_2xBoxB_J102<br>2) sgRNA | 1) TrrnB<br>2) TrrnB | This work |

[a]All SoxS genes contain two mutations (R93A and S101A)
[b]All dCas9-AsiA genes in this work are dCas9-AsiA_m2.1 with the same protein sequence as described previously.[2]
[c]MarA and TetD mutants were generated as listed in Figure S2B.

**Table S3.** sgRNA/scRNA/crRNA spacer sequences

| Spacer | DNA sequence | Target strand[a] | Distance to TSS[b] |
|---|---|---|---|
| **sgRNA/scRNA for Spy-dCas9** | | | |
| **hAAVS1** | GGGGCCACTAGGGACAGGAT | Off-target | NA |
| **J306** | TTGTGTCCAGAACGCTCCGT | Non-Template | -81 |
| **RR2** | TGGAACCGTACTGGAACTGC | Non-Template | 188 |
| **H2** | GAAGATCCGGCCTGCAGCCA | Non-Template | -91 |
| **H3** | GGCTCGAGTCGACAGTTCAT | Non-Template | -128 |
| **H4** | CTACGGAACTCTTGTGCGTA | Template | -182 |
| **H5** | GCAAAAGCTCATTTCTGAAG | Template | -252 |
| **G6** | AAAGCTGTTGTGGCAAAAAA | Non-Template | -91 |
| **J101** | TGGGTTCCACCGGATACCTC | Template | -40 |
| **J102** | AGGTATCCGGTGGAACCCAA | Non-Template | -61 |
| **J103** | AGGCGTCCTTTGGGTTCCAC | Template | -50 |
| **J104** | TGGAACCCAAAGGACGCCTT | Non-Template | -71 |
| **J105** | CGGTTACCAAAGGCGTCCTT | Template | -60 |
| **J106** | AGGACGCCTTTGGTAACCGC | Non-Template | -81 |
| **J107** | CGGTGTCCTGCGGTTACCAA | Template | -70 |
| **J108** | TGGTAACCGCAGGACACCGC | Non-Template | -91 |
| **J109** | AGGTATCCTGCGGTGTCCTG | Template | -80 |
| **J110** | AGGACACCGCAGGATACCTG | Non-Template | -101 |
| **J111** | GGGCGACCTCAGGTATCCTG | Template | -90 |
| **J112** | AGGATACCTGAGGTCGCCCG | Non-Template | -111 |
| **J113** | GGGCCACCACGGGCGACCTC | Template | -100 |
| **J114** | AGGTCGCCCGTGGTGGCCCA | Non-Template | -121 |
| **J115** | TGGTGACCATGGGCCACCAC | Template | -110 |
| **J116** | TGGTGGCCCATGGTCACCAT | Non-Template | -131 |
| **J117** | GGGTGACCTATGGTGACCAT | Template | -120 |
| **J118** | TGGTCACCATAGGTCACCCT | Non-Template | -141 |
| **J119** | TGGTTGCCAAGGGTGACCTA | Template | -130 |
| **J120** | AGGTCACCCTTGGCAACCAA | Non-Template | -151 |
| **J121** | AGGACACCTTTGGTTGCCAA | Template | -140 |

| sgRNA/scRNA for Sth1-dCas9[a] | | | |
|---|---|---|---|
| **hAAVS1** | GGGGCCACTAGGGACAGGAT | Off-target | NA |
| **J306** | TTGTGTCCAGAACGCTCCGT | Bottom | -81 |
| **Sth1-T1** | ccgtccgacggtccggttat | Template | 434 from TSS |
| **Sth1-RBS-t** | gctagcGAATTCATTAAAGA | RBS | 14 from TSS |
| **K101** | TGAGAATGGGTTCACAGCGA | Template | -50 |
| **K102** | CCAGAATAAAGTCTCGCAGA | Template | -70 |
| **K103** | AGAGAATGCGGACCACTTGA | Template | -90 |
| **K104** | TGAGAAAGGTGATAGTGCGA | Template | -110 |
| **K105** | CCAGAAACGACCGGCAGACG | Template | -130 |
| **K106** | CGAGAATAGACTTCGACTGC | Template | -150 |
| **K107** | TGAGAATGCCTCCCACCGAG | Template | -170 |
| **K108** | GGAGAAAGACATCCAAGTCT | Template | -190 |
| **crRNA for dCas12a[b]** | | | |
| **hAAVS1** | GGGGCCACTAGGGACAGGAT | Off-target | NA |
| **12a-T1** | AAAGTTCGTATGGAAGGTTC | Template | 42 |
| **12a-RBS-b** | TCCTCTTTAATGAATTCgct | RBS | 17 from TSS |
| **I101** | CGTCGAATCATCGGGCTTTC | Template | -40 |
| **I102** | ATATGGCTCCCGGTCATTTC | Template | -60 |
| **I103** | GTGGCACAACTATAGGTTTA | Template | -80 |
| **I104** | TCTGCGGCCAGCACCATTTC | Template | -100 |
| **I105** | TCGCTATGCGGACGGCTTTA | Template | -120 |
| **I106** | CTGCCAAGCCTGGGCCTTTA | Template | -140 |
| **I107** | GCTTTGAAGTCGGTACTTTA | Template | -160 |
| **I108** | GCTGACACCACAGTCCTTTC | Template | -180 |

[a]gRNA hairpin of Sth1 dCas9 is distinct from that of Spy-dCas9.
[b]crRNA spacer locates at 3'-end different from that of 5'-end in the Cas9 case. See DNA sequences section for further details.

**Table S4.** sgRNA target site positions for alternative promoters

| Spacer | J1 | J1-J3(100) | J1-J2(100)-J3(100) | J1-J3(122) |
|---|---|---|---|---|
| **Non-Template strand**[a] | | | | |
| J101 | -40 | -140 | -240 | -162 |
| J103 | -50 | -150 | -250 | -172 |
| J105 | -60 | -160 | -260 | -182 |
| J107 | -70 | -170 | -270 | -192 |
| J109 | -80 | -180 | -280 | -202 |
| J111 | -90 | -190 | -290 | -212 |
| J113 | -100 | -200 | -300 | -222 |
| J115 | -110 | -210 | -310 | -232 |
| J117 | -120 | -220 | -320 | -242 |
| J119 | -130 | -230 | -330 | -252 |
| **Template strand**[b] | | | | |
| J102 | -61 | -161 | -261 | -183 |
| J104 | -71 | -171 | -271 | -193 |
| J106 | -81 | -181 | -281 | -203 |
| J108 | -91 | -191 | -291 | -213 |
| J110 | -101 | -201 | -301 | -223 |
| J112 | -111 | -211 | -311 | -233 |
| J114 | -121 | -221 | -321 | -243 |
| J116 | -131 | -231 | -331 | -253 |
| J118 | -141 | -241 | -341 | -263 |
| J120 | -151 | -251 | -351 | -273 |

[a]gRNAs targeting Non-Template strand match the DNA sequence on the Top strand.
[b]gRNAs targeting Template strand match the DNA sequence on the Bottom strand.

**Table S5.** Plasmids used in each figure

| Figures (Activators) | p15A-CmR | pSC101-AmpR | ColE1-KmR |
|---|---|---|---|
| 2A (MCP-SoxS) | pCK005.01-21 | pJF076 (J1-BBa_J23117) | |
| 2A (αNTD-MCP) | pCD297.01-21 | pJF076 (J1-BBa_J23117) | |
| 2A (dCas9-AsiA) | pCK320 | pJF076 (J1-BBa_J23117) | pAK030.01-12 |
| 2A (PspF-λN22) | pCK147.01-20 | pJF076 (J1-BBa_J23117) | |
| 2B (MCP-SoxS) | pCK005.01-21 | pAK070 (J1-pspAp) | |
| 2B (PspF-λN22) | pCK147.01-20 | pAK070 (J1-pspAp) | |
| 3A (MCP-SoxS) | pCD442 | pCK161.06-09 (J1-BBa_J23117) | |
| 3A (αNTD-MCP) | pCD037 | pCK161.06-09 (J1-BBa_J23117) | |
| 3A (αNTD-MCP-SoxS) | pCK692 | pCK161.06-09 (J1-BBa_J23117) | |
| 3B (MCP-SoxS) | pCD442 | pCK161.06-09 (J1-BBa_J23117) | |
| 3B (MCP-SoxS-SYNZIP6) | pCK882 | pCK161.06-09 (J1-BBa_J23117) | |
| 3B (MCP-SoxS-SYNZIP6 and αNTD-SYNZIP5) | pCK883 | pCK161.06-09 (J1-BBa_J23117) | |
| 4A (MCP-SoxS) | pAK007 | pAK011 (J1-J3(122)-BBa_J23117) | pCK303.306<br>pCK303.107 |
| 4A (dCas9-AsiA) | pCK320 | pAK011 (J1-J3(122)-BBa_J23117) | pCK303.306<br>pCK303.107 |
| 4A (dCas9-AsiA + MCP-SoxS) | pCK321 | pAK011 (J1-J3(122)-BBa_J23117) | pCK303.306<br>pCK303.107 |
| 4B (dCas9-AsiA + Sth1-dCas9 + MCP-SoxS) | pKO025 | pKO023 (J1-J3(122)-BBa_J23117 with Sth1 PAM) | pKO024.AAV-AAV<br>pKO024.306-AAV<br>pKO024.AAV-107<br>pKO024.306-107 |
| 5B (MCP-SoxS + PspF-λN22) | pCK660 | pCK662.-1 (J1-pspAp.-1) | pCK667.AAV-AAV<br>pCK667.102-AAV<br>pCK667.102-(105-112) |
| 5C (MCP-SoxS + PspF-λN22) | pCK660 | pCK662.-1 (J1-pspAp.-1) | pCK667.AAV-AAV<br>pCK667.102-AAV<br>pCK667.AAV-110<br>pCK667.102-110<br>pCK667.102-112<br>pCK962.102-110<br>pCK962.102-112 |
| 5D (αNTD-MCP + PspF-λN22) | pCK761 | pCK662.-1 (J1-pspAp.-1) | pCK667.AAV-AAV<br>pCK667.102-AAV<br>pCK667.102-(105-112) |
| 5D (MCP-TetD | pCK761 | pCK662.-1 (J1-pspAp.-1) | pCK667.AAV-AAV |

| Figures (Activators) | p15A-CmR | pSC101-AmpR | ColE1-KmR |
|---|---|---|---|
| + PspF-λN22) | | | pCK667.102-AAV<br>pCK667.102-(105-112) |
| 5D (PspF-λN22) | pCK071 | pCK662.-1 (J1-pspAp.-1) | pCK667.AAV-AAV<br>pCK667.102-AAV<br>pCK667.102-(105-112) |
| 5E (PspF-λN22) | pCK071 | pCK662.-1 (J1-pspAp.-1) | pCK667.AAV-AAV<br>pCK667.102-AAV<br>pCK667.AAV-110<br>pCK667.102-110<br>pCK667.102-112<br>pCK962.102-110<br>pCK962.102-112 |
| S1A (MCP-SoxS) | pCK005.01-21 | pJF076 (J1-BBa_J23117) | |
| S1A (MCP-SoxS) | pCK005.01-21 | pCK001 (J1-sodCp) | |
| S1A (MCP-SoxS) | pCK005.01-21 | pCK002 (J1-glnAp2) | |
| S1A (MCP-SoxS) | pCK005.01-21 | pCK003 (J1-rdgBp) | |
| S1A (MCP-SoxS) | pCK005.01-21 | pCK004 (J1-yieEp) | |
| S1B (MCP-SoxS) | pCK005.06/09/AA<br>V | pCK001 (J1-sodCp),<br>pCK067.N (J1-sodCp_phase-shift) | |
| S2A (MCP-SoxS) | pCD442 | pCK161.X (J1-BBa_J23117),<br>pCK101.X (J3-BBa_J23117) | |
| S2A (MCP-TetD) | pCD178 | pJF076 (J1-BBa_J23117),<br>pJF143.J3 (J3-BBa_J23117) | |
| S2B (MCP-MarA-mut) | pCK090.X | pJF143.J3 (J3-BBa_J23117) | |
| S2B (MCP-TetD-mut) | pCK094.X | pJF143.J3 (J3-BBa_J23117) | |
| S3B (αNTD-MCP) | pCD297.X | pJF076 (J1-BBa_J23117) | |
| S3B (αNTD-PCP) | pRC099.X | pJF076 (J1-BBa_J23117) | |
| S4B (MCP-Crl) | pCK083.X | pCK067.3 (J1-sodCp.-3) | |
| S5A (dCas9-AsiA) | pCK309.X | pCD249 (W1-BBa_J23117) | |
| S5B (dCas9-AsiA) | pCD442,<br>pCK226 | CD38 (BBa_J23119-mRFP) | pAK030.gAAV<br>pAK030.RR2 |
| S5C (dCas9-AsiA) | pCK320 | pCD249 (W1-BBa_J23117) | pAK030.gAAV<br>pAK030.H2-H5 |
| S6B (MCP-SoxS) | pCK005.X | pJF076 (J1-BBa_J23117),<br>pJF143.J3 (J3-BBa_J23117),<br>pAK011 (J1-J3(122)-BBa_J23117) | |
| S6C (MCP-TetD) | pCD296.X | pJF076 (J1-BBa_J23117),<br>pJF143.J3 (J3-BBa_J23117), | |

| Figures (Activators) | p15A-CmR | pSC101-AmpR | ColE1-KmR |
|---|---|---|---|
| | | pAK011 (J1-J3(122)-BBa_J23117) | |
| S6D (αNTD-MCP) | pCD297.X | pJF076 (J1-BBa_J23117),<br>pJF143.J3 (J3-BBa_J23117),<br>pAK011 (J1-J3(122)-BBa_J23117) | |
| S7B (dCas9-AsiA) | pCK320 | pJF076 (J1-BBa_J23117),<br>pCK702 (J1-J3(100)-<br>BBa_J23117),<br>pCK703 (J1-J2(100)-J3(100)-<br>BBa_J23117) | pAK030.gAAV<br>pAK030.01-12 |
| S7C (dCas9-AsiA) | pCK320 | pCD249 (W1-BBa_J23117),<br>pAK011 (J1-J3(122)-BBa_J23117) | pAK030.gAAV<br>pAK030.01-12<br>pAK030.H2-H5 |
| S7C (MCP-SoxS) | pCD442 | pJF076 (J1-BBa_J23117) | pCK303.AAV<br>pCK303.306 |
| S8A (PspF-λN22) | pCK147.X | pCK069 (pspAp-G6),<br>pCK070 (J1-pspAp) | |
| S8B (PspF-λN22) | pCK147.X | pCK002 (J1-glnAp2),<br>pCK299 (J1-glnAp2*) | |
| S8C (PspF-λN22) | pCK147.X | pCK070 (J1-pspAp),<br>pCK662.N (J1-pspAp_phase-shift) | |
| S9A (MCP-SoxS) | pCD442 | pCK161.X (J1-BBa_J23117) | |
| S9A (MCP-TetD) | pCD178 | pCK161.X (J1-BBa_J23117) | |
| S9A (MCP-SoxS-TetD) | pCK693 | pCK161.X (J1-BBa_J23117) | |
| S9B (MCP-SoxS) | pCD442 | pCK101.X (J3-BBa_J23117) | |
| S9B (αNTD-MCP) | pCD037 | pCK101.X (J3-BBa_J23117) | |
| S9B (αNTD-MCP-SoxS) | pCK692 | pCK101.X (J3-BBa_J23117) | |
| S9B (MCP-TetD) | pCD178 | pCK101.X (J3-BBa_J23117) | |
| S9B (MCP-SoxS-TetD) | pCK693 | pCK101.X (J3-BBa_J23117) | |
| S10 (αNTD-MCP) | pCD297.X | pJF076 (J1-BBa_J23117) | |
| S10 (αNTD-PCP) | pRC099.X | pJF076 (J1-BBa_J23117) | |
| S10 (αNTD-SYNZIP5 +<br>MCP-SYNZIP6) | pCK157 | pCK161.X (J1-BBa_J23117) | |
| S10 (αNTD-SYNZIP5 +<br>PCP-SYNZIP6) | pCK770 | pJF076 (J1-BBa_J23117) | pCK732.01-12 |
| S11A (MCP-SoxS) | pAK007 | pCD249 (W1-BBa_J23117) | pCK303.X<br>pAK030.X |
| S11A (dCas9-AsiA) | pCK320 | pCD249 (W1-BBa_J23117) | pCK303.X |

| Figures (Activators) | p15A-CmR | pSC101-AmpR | ColE1-KmR |
| --- | --- | --- | --- |
| | | | pAK030.X |
| S11A (dCas9-AsiA<br>+ MCP-SoxS) | pCK321 | pCD249 (W1-BBa_J23117) | pCK303.X<br>pAK030.X |
| S11B (MCP-SoxS) | pAK007 | pAK011 (J1-J3(122)-BBa_J23117) | pCK303.AAV<br>pCK303.306 |
| S11B (dCas9-AsiA) | pCK320 | pAK011 (J1-J3(122)-BBa_J23117) | pAK030.gAAV<br>pAK030.107 |
| S11B (dCas9-AsiA<br>+ MCP-SoxS) | pCK321 | pAK011 (J1-J3(122)-BBa_J23117) | pAK030.gAAV<br>pAK030.107<br>pCK303.AAV<br>pCK303.306 |
| S12 (MCP-AsiA) | pAK010 | pCD249 (W1-BBa_J23117) | pAK030.gAAV<br>pAK030.H2-H5 |
| S12 (dCas9-AsiA) | pCK320 | pCD249 (W1-BBa_J23117) | pAK030.gAAV<br>pAK030.H2-H5 |
| S13A (Sth1-dCas9-AsiA) | pCK1004 | pJF143.J3.110 (J3-BBa_J23110) | pCK1005.AAV<br>pCK1005.Sth1-T1<br>pCK1005.RBS-t |
| S13B (Sth1-dCas9-AsiA) | pCK1004 | pCK1006.N (K1T-BBa_J23117),<br>pCK1007.N (K1B-BBa_J23117) | pCK1005.01-08 |
| S14A (Lb-dCas12a-AsiA) | pCK1008 | pJF143.J3.110 (J3-BBa_J23110) | pCK1010.AAV<br>pCK1010.12a-T1<br>pCK1010.RBS-b |
| S14B (Lb-dCas12a-AsiA) | pCK1008 | pCK1010.N (I1T-BBa_J23117),<br>pCK1011.N (I1B-BBa_J23117) | pCK1010.01-08 |
| S14C (Lb-dCas12a-AsiA) | pCK1008 | pCK1011.N (I1B-BBa_J23117) | pCK1010.01 |
| S15 (Sth1-dCas9 + MCP-SoxS) | pAK140 | pAK021 (J3-BBa_J23117 with Sth1 PAM) | pKO024.X |
| S16A (PspF-λN22) | pCK071 | pCK070 (J1-pspAp) | pCK676.01-12 |
| S16A (PspF-MCP) | pCK694 | pCK070 (J1-pspAp) | pCK303.01-12<br>pCK697.01-12 |
| S16A (PspF-PCP) | pCK934 | pCK070 (J1-pspAp) | pCK732.01-12 |
| S16B (PspF-λN22) | pCK696 | pCK070 (J1-pspAp) | pCK676.01-12 |
| S16B (PspF-MCP) | pCK695 | pCK070 (J1-pspAp) | pCK303.01-12<br>pCK697.01-12 |
| S17A (MCP-SoxS<br>+ PspF-λN22) | pCK660 | pCK070 (J1-pspAp),<br>pCK662.N (J1-pspAp.N) | pCK667.AAV-AAV<br>pCK667.102-AAV<br>pCK667.102-110 |
| S17B (αNTD-MCP | pCK761 | pCK662.-1 (J1-pspAp.-1) | pCK667.AAV-AAV |

| Figures (Activators) | p15A-CmR | pSC101-AmpR | ColE1-KmR |
|---|---|---|---|
| + PspF-λN22) | | | pCK667.102-AAV<br>pCK667.102-(105-112) |
| S17B (MCP-TetD<br>+ PspF-λN22) | pCK761 | pCK662.-1 (J1-pspAp.-1) | pCK667.AAV-AAV<br>pCK667.102-AAV<br>pCK667.102-(105-112) |
| S17B (PspF-λN22) | pCK071 | pCK662.-1 (J1-pspAp.-1) | pCK667.AAV-AAV<br>pCK667.102-AAV<br>pCK667.102-(105-112) |
| S18A (αNTD-MCP<br>+ PspF-λN22) | pCK071 | pCK662.-1 (J1-pspAp.-1) | pCK667.AAV-AAV<br>pCK667.102-AAV<br>pCK667.AAV-110<br>pCK667.102-110<br>pCK667.102-112<br>pCK962.102-110<br>pCK962.102-112 |
| S18B (MCP-TetD<br>+ PspF-λN22) | pCK071 | pCK662.-1 (J1-pspAp.-1) | pCK667.AAV-AAV<br>pCK667.102-AAV<br>pCK667.AAV-110<br>pCK667.102-110<br>pCK667.102-112<br>pCK962.102-110<br>pCK962.102-112 |

**Table S6**. Design rules of different CRISPRa systems not tested in this work (extended from Table 1)

| **Activator** | **Mechanism group** | **Reported working distance (Strand preference)[a]** | **Organisms** |
|---|---|---|---|
| λcII (Dong et al., 2018)[1] | Interact with RpoA | -71 (T) | *Escherichia coli* |
| RemA (Wu et al., 2020)[8] | Unknown | -90 (NT) | *Bacillus subtilis* |
| RpoZ (Bikard et al., 2013; Dong et al., 2018)[1,9] | RNAP assembly | -60 to -100 (NT) | *Escherichia coli, Bacillus subtilis, Lysobacter enzymogenes, Myxococcus xanthus* |
| RpoA (Lu et al., 2019)[10] | RNAP assembly | -267 to -415 (T) | *Bacillus subtilis* |
| RpoD (Chen et al., 2022)[11] | RNAP assembly | -199 to -216 (NT) | *Shewanella oneidensis* |
| RpoN (Peng et al., 2018)[12] | RNAP assembly | -53 to -103 (NT) | *Myxococcus xanthus* |
| NorR (Liu et al., 2019)[6] | $\sigma^{54}$ activation | -91 to -131 (NT) | *Escherichia coli* |
| WtsA (Liu et al., 2019)[6] | $\sigma^{54}$ activation | -91 to -131 (NT) | *Escherichia coli* |

[a]T stands for Template strand and NT stands for Non-Template strand.

**Table S7.** Potential cooperativity of effectors investigated in this work

| Effectors | SoxS-family | RNAP-subunit | AsiA | PspF |
|---|---|---|---|---|
| SoxS-family | **S** | | | |
| RNAP-subunit | **D** | **S** | | |
| AsiA | **O** | **O** | **S** | |
| PspF | **O** | **O** | **U** | **S** |

Representative labels are listed below for each activator pair.

**S**: Same activator, does not fit different mechanism criteria

**U**: Unlikely as only one sigma factor will be present

**D**: Same target site, prefer direct fusion.

**O**: Distinct target sites, prefer orthogonal recruitment.

### DNA sequences

CRISPRa construct sequences: dCas proteins and activators

Sequence features are color-coded.
Start and Stop codons are **bolded**.

>Sp.pCas9-Spy-dCas9-dblT
TTACGAAATCATCCTGTGGAGCTTAGTAGGTTTAGCAAGATGGCAGCGCCTAAATGTAGAATGATAAAAG
GATTAAGAGATTAATTTCCCTAAAAATGATAAAACAAGCGTTTTGAAAGCGCTTGTTTTTTTGGTTTGCA
GTCAGAGTAGAATAGAAGTATCAAAAAAAGCACCGACTCGGTGCCACTTTTTCAAGTTGATAACGGACTA
GCCTTATTTTAACTTGCTATGCTGTTTTGAATGGTTCCAACAAGATTATTTTATAACTTTTATAACAAAT
AATCAAGGAGAAATTCAAAGAAATTTATCAGCCATAAAACAATACTTAATACTATAGAATGATAACAAAA
TAAACTACTTTTTAAAAGAATTTTGTGTTATAATCTATTTATTATTAAGTATTGGGTAATATTTTTTGAA
GAGATATTTTGAAAAAGAAAAATTAAAGCATATTAAACTAATTTCGGAGGTCATTAAAACTATTATTGAA
ATCATCAAACTCATTATGGATTTAATTTAAACTTTTTATTTTAGGAGGCAAAA**ATG**GATAAGAAATACTC
AATAGGCTTAGCTATCGGCACAAATAGCGTCGGATGGGCGGTGATCACTGATGAATATAAGGTTCCGTCT
AAAAAGTTCAAGGTTCTGGGAAATACAGACCGCCACAGTATCAAAAAAAATCTTATAGGGGCTCTTTTAT
TTGACAGTGGAGAGACAGCGGAAGCGACTCGTCTCAAACGGACAGCTCGTAGAAGGTATACACGTCGGAA
GAATCGTATTTGTTATCTACAGGAGATTTTTTCAAATGAGATGGCGAAAGTAGATGATAGTTTCTTTCAT
CGACTTGAAGAGTCTTTTTTGGTGGAAGAAGACAAGAAGCATGAACGTCATCCTATTTTTGGAAATATAG
TAGATGAAGTTGCTTATCATGAGAAATATCCAACTATCTATCATCTGCGAAAAAAATTGGTAGATTCTAC
TGATAAAGCGGATTTGCGCTTAATCTATTTGGCCTTAGCGCATATGATTAAGTTTCGTGGTCATTTTTTG
ATTGAGGGAGATTTAAATCCTGATAATAGTGATGTGGACAAACTATTTATCCAGTTGGTACAAACCTACA
ATCAATTATTTGAAGAAAACCCTATTAACGCAAGTGGAGTAGATGCTAAAGCGATTCTTTCTGCACGATT
GAGTAAATCAAGACGATTAGAAAATCTCATTGCTCAGCTCCCCGGTGAGAAGAAAAATGGCTTATTTGGG
AATCTCATTGCTTTGTCATTGGGTTTGACCCCTAATTTTAAATCAAATTTTGATTTGGCAGAAGATGCTA
AATTACAGCTTTCAAAAGATACTTACGATGATGATTTAGATAATTTATTGGCGCAAATTGGAGATCAATA
TGCTGATTTGTTTTTGGCAGCTAAGAATTTATCAGATGCTATTTTACTTTCAGATATCCTAAGAGTAAAT
ACTGAAATAACTAAGGCTCCCCTATCAGCTTCAATGATTAAACGCTACGATGAACATCATCAAGACTTGA
CTCTTTTAAAAGCTTTAGTTCGACAACAACTTCCAGAAAAGTATAAAGAAATCTTTTTTGATCAATCAAA
AAACGGATATGCAGGTTATATTGATGGGGGAGCTAGCCAAGAAGAATTTTATAAATTTATCAAACCAATT
TTAGAAAAAATGGATGGTACTGAGGAATTATTGGTGAAACTAAATCGTGAAGATTTGCTGCGCAAGCAAC
GGACCTTTGACAACGGCTCTATTCCCCATCAAATTCACTTGGGTGAGCTGCATGCTATTTTGAGAAGACA
AGAAGACTTTTATCCATTTTTAAAAGACAATCGTGAGAAGATTGAAAAAATCTTGACTTTTCGAATTCCT
TATTATGTTGGTCCATTGGCGCGTGGCAATAGTCGTTTTGCATGGATGACTCGGAAGTCTGAAGAAACAA
TTACCCCATGGAATTTTGAAGAAGTTGTCGATAAAGGTGCTTCAGCTCAATCATTTATTGAACGCATGAC
AAACTTTGATAAAAATCTTCCAAATGAAAAAGTACTACCAAAACATAGTTTGCTTTATGAGTATTTTACG
GTTTATAACGAATTGACAAAGGTCAAATATGTTACTGAAGGAATGCGAAAACCAGCATTTCTTTCAGGTG
AACAGAAGAAAGCCATTGTTGATTTACTCTTCAAAACAAATCGAAAAGTAACCGTTAAGCAATTAAAAGA
AGATTATTTCAAAAAAATAGAATGTTTTGATAGTGTTGAAATTTCAGGAGTTGAAGATAGATTTAATGCT
TCATTAGGTACCTACCATGATTTGCTAAAAATTATTAAAGATAAAGATTTTTTGGATAATGAAGAAAATG
AAGATATCTTAGAGGATATTGTTTTAACATTGACCTTATTTGAAGATAGGGAGATGATTGAGGAAAGACT
TAAAACATATGCTCACCTCTTTGATGATAAGGTGATGAAACAGCTTAAACGTCGCCGTTATACTGGTTGG
GGACGTTTGTCTCGAAAATTGATTAATGGTATTAGGGATAAGCAATCTGGCAAAACAATATTAGATTTTT

TGAAATCAGATGGTTTTGCCAATCGCAATTTTATGCAGCTGATCCATGATGATAGTTTGACATTTAAAGA
AGACATTCAAAAAGCACAAGTGTCTGGACAAGGCGATAGTTTACATGAACATATTGCAAATTTAGCTGGT
AGCCCTGCTATTAAAAAAGGTATTTTACAGACTGTAAAAGTTGTTGATGAATTGGTCAAAGTAATGGGGC
GGCATAAGCCAGAAAATATCGTTATTGAAATGGCACGTGAAAATCAGACAACTCAAAAGGGCCAGAAAAA
TTCGCGAGAGCGTATGAAACGAATCGAAGAAGGTATCAAAGAATTAGGAAGTCAGATTCTTAAAGAGCAT
CCTGTTGAAAATACTCAATTGCAAAATGAAAAGCTCTATCTCTATTATCTCCAAAATGGAAGAGACATGT
ATGTGGACCAAGAATTAGATATTAATCGTTTAAGTGATTATGATGTCGATGCCATTGTTCCACAAAGTTT
CCTTAAAGACGATTCAATAGACAATAAGGTCTTAACGCGTTCTGATAAAAATCGTGGTAAATCGGATAAC
GTTCCAAGTGAAGAAGTAGTCAAAAAGATGAAAAACTATTGGAGACAACTTCTAAACGCCAAGTTAATCA
CTCAACGTAAGTTTGATAATTTAACGAAAGCTGAACGTGGAGGTTTGAGTGAACTTGATAAAGCTGGTTT
TATCAAACGCCAATTGGTTGAAACTCGCCAAATCACTAAGCATGTGGCACAAATTTTGGATAGTCGCATG
AATACTAAATACGATGAAAATGATAAACTTATTCGAGAGGTTAAAGTGATTACCTTAAAATCTAAATTAG
TTTCTGACTTCCGAAAAGATTTCCAATTCTATAAAGTACGTGAGATTAACAATTACCATCATGCCCATGA
TGCGTATCTAAATGCCGTCGTTGGAACTGCTTTGATTAAGAAATATCCAAAACTTGAATCGGAGTTTGTC
TATGGTGATTATAAAGTTTATGATGTTCGTAAAATGATTGCTAAGTCTGAGCAAGAAATAGGCAAAGCAA
CCGCAAAATATTTCTTTTACTCTAATATCATGAACTTCTTCAAAACAGAAATTACACTTGCAAATGGAGA
GATTCGCAAACGCCCTCTAATCGAAACTAATGGGGAAACTGGAGAAATTGTCTGGGATAAAGGGCGAGAT
TTTGCCACAGTGCGCAAAGTATTGTCCATGCCCCAAGTCAATATTGTCAAGAAAACAGAAGTACAGACAG
GCGGATTCTCCAAGGAGTCAATTTTACCAAAAAGAAATTCGGACAAGCTTATTGCTCGTAAAAAAGACTG
GGATCCAAAAAAATATGGTGGTTTTGATAGTCCAACGGTAGCTTATTCAGTCCTAGTGGTTGCTAAGGTG
GAAAAAGGGAAATCGAAGAAGTTAAAATCCGTTAAAGAGTTACTAGGGATCACAATTATGGAAAGAAGTT
CCTTTGAAAAAAATCCGATTGACTTTTTAGAAGCTAAAGGATATAAGGAAGTTAAAAAAGACTTAATCAT
TAAACTACCTAAATATAGTCTTTTTGAGTTAGAAAACGGTCGTAAACGGATGCTGGCTAGTGCCGGAGAA
TTACAAAAAGGAAATGAGCTGGCTCTGCCAAGCAAATATGTGAATTTTTTATATTTAGCTAGTCATTATG
AAAAGTTGAAGGGTAGTCCAGAAGATAACGAACAAAAACAATTGTTTGTGGAGCAGCATAAGCATTATTT
AGATGAGATTATTGAGCAAATCAGTGAATTTTCTAAGCGTGTTATTTTAGCAGATGCCAATTTAGATAAA
GTTCTTAGTGCATATAACAAACATAGAGACAAACCAATACGTGAACAAGCAGAAAATATTATTCATTTAT
TTACGTTGACGAATCTTGGAGCTCCCGCTGCTTTTAAATATTTTGATACAACAATTGATCGTAAACGATA
TACGTCTACAAAAGAAGTTTTAGATGCCACTCTTATCCATCAATCCATCACTGGTCTTTATGAAACACGC
ATTGATTTGAGTCAGCTAGGAGGTGAC**TAA**CTCGAGTAAGGATCTCCAGGCATCAAATAAAACGAAAGGC
TCAGTCGAAAGACTGGGCCTTTCGTTTTATCTGTTGTTTGTCGGTGAACGCTCTCTACTAGAGTCACACT
GGCTCACCTTCGGGTGGGCCTTTCTGCGTTTATA

>BBa_J23107-RBS-MCP-SoxS(2mut)-BBa_B1002
TTTACGGCTAGCTCAGCCCTAGGTATTATGCTAGCGAATTCATTAAAGAGGAGAAAGGTACC**ATG**GGGCC
CGCTTCTAACTTTACTCAGTTCGTTCTCGTCGACAATGGCGGAACTGGCGACGTGACTGTCGCCCCAAGC
AACTTCGCTAACGGGATCGCTGAATGGATCAGCTCTAACTCGCGTTCACAGGCTTACAAAGTAACCTGTA
GCGTTCGTCAGAGCTCTGCGCAGAATCGCAAATACACCATCAAAGTCGAGGTGCCTAAAGGCGCCTGGCG
TTCGTACTTAAATATGGAACTAACCATTCCAATTTTCGCCACGAATTCCGACTGCGAGCTTATTGTTAAG
GCAATGCAAGGTCTCCTAAAAGATGGAAACCCGATTCCCTCAGCAATCGCAGCAAACTCCGGCATCTACG
GTGGCGGAGGTAGCATGTCCCATCAGAAAATTATTCAGGATCTTATCGCATGGATTGACGAGCATATTGA
CCAGCCGCTTAACATTGATGTAGTCGCAAAAAAATCAGGCTATTCAAAGTGGTACTTGCAACGAATGTTC
CGCACGGTGACGCATCAGACGCTTGGCGATTACATTCGCCAACGCCGCCTGTTACTGGCCGCCGTTGAGT
TGCGCACCACCGAGCGTCCGATTTTTGATATCGCAATGGACCTGGGTTATGTCTCGCAGCAGACCTTCTC

CCGCGTTTTCGCGCGGCAGTTTGATCGCACTCCCGCGGATTATCGCCACCGCCTG**TAA**GCGGCCGCCACG
CAAAAAACCCCGCTTCGGCGGGGTTTTTTCGC

>BBa_J23107-RBS-αNTD-MCP-BBa_B1002
TTTACGGCTAGCTCAGCCCTAGGTATTATGCTAGCGAATTCATTAAAGAGGAGAAAGGTACC**ATG**GGGCC
CATGCAGGGTTCTGTGACAGAGTTTCTAAAACCGCGCCTGGTTGATATCGAGCAAGTGAGTTCGACGCAC
GCCAAGGTGACCCTTGAGCCTTTAGAGCGTGGCTTTGGCCATACTCTGGGTAACGCACTGCGCCGTATTC
TGCTCTCATCGATGCCGGGTTGCGCGGTGACCGAGGTTGAGATTGATGGTGTACTACATGAGTACAGCAC
CAAAGAAGGCGTTCAGGAAGATATCCTGGAAATCCTGCTCAACCTGAAAGGGCTGGCGGTGAGAGTTCAG
GGCAAAGATGAAGTTATTCTTACCTTGAATAAATCTGGCATTGGCCCTGTGACTGCAGCCGATATCACCC
ACGACGGTGATGTCGAAATCGTCAAGCCGCAGCACGTGATCTGCCACCTGACCGATGAGAACGCGTCTAT
TAGCATGCGTATCAAAGTTCAGCGCGGTCGTGGTTATGTGCCGGCTTCTACCCGAATTCATTCGGAAGAA
GATGAGCGCCCAATCGGCCGTCTGCTGGTCGACGCATGCTACAGCCCTGTGGAGCGTATTGCCTACAATG
TTGAAGCAGCGCGTGTAGAACAGCGTACCGACCTGGACAAGCTGGTCATCGAAATGGAAACCAACGGCAC
AATCGATCCTGAAGAGGCGATTCGTCGTGCGGCAACCATTCTGGCTGAACAACTGGAAGCTTTCGTTGAC
TTACGTGATGTACGTCAGCCTGAAGTGAAAGAAGAGAAACCAGAGGCTTCTAACTTTACTCAGTTCGTTC
TCGTCGACAATGGCGGAACTGGCGACGTGACTGTCGCCCCAAGCAACTTCGCTAACGGGATCGCTGAATG
GATCAGCTCTAACTCGCGTTCACAGGCTTACAAAGTAACCTGTAGCGTTCGTCAGAGCTCTGCGCAGAAT
CGCAAATACACCATCAAAGTCGAGGTGCCTAAAGGCGCCTGGCGTTCGTACTTAAATATGGAACTAACCA
TTCCAATTTTCGCCACGAATTCCGACTGCGAGCTTATTGTTAAGGCAATGCAAGGTCTCCTAAAAGATGG
AAACCCGATTCCCTCAGCAATCGCAGCAAACTCCGGCATCTAC**TAA**GCGGCCGCCACGCAAAAAACCCCG
CTTCGGCGGGGTTTTTTCGC

>BBa_J23107-RBS-αNTD-PCP-BBa_B1002
TTTACGGCTAGCTCAGCCCTAGGTATTATGCTAGCGAATTCATTAAAGAGGAGAAAGGTACC**ATG**GGGCC
CATGCAGGGTTCTGTGACAGAGTTTCTAAAACCGCGCCTGGTTGATATCGAGCAAGTGAGTTCGACGCAC
GCCAAGGTGACCCTTGAGCCTTTAGAGCGTGGCTTTGGCCATACTCTGGGTAACGCACTGCGCCGTATTC
TGCTCTCATCGATGCCGGGTTGCGCGGTGACCGAGGTTGAGATTGATGGTGTACTACATGAGTACAGCAC
CAAAGAAGGCGTTCAGGAAGATATCCTGGAAATCCTGCTCAACCTGAAAGGGCTGGCGGTGAGAGTTCAG
GGCAAAGATGAAGTTATTCTTACCTTGAATAAATCTGGCATTGGCCCTGTGACTGCAGCCGATATCACCC
ACGACGGTGATGTCGAAATCGTCAAGCCGCAGCACGTGATCTGCCACCTGACCGATGAGAACGCGTCTAT
TAGCATGCGTATCAAAGTTCAGCGCGGTCGTGGTTATGTGCCGGCTTCTACCCGAATTCATTCGGAAGAA
GATGAGCGCCCAATCGGCCGTCTGCTGGTCGACGCATGCTACAGCCCTGTGGAGCGTATTGCCTACAATG
TTGAAGCAGCGCGTGTAGAACAGCGTACCGACCTGGACAAGCTGGTCATCGAAATGGAAACCAACGGCAC
AATCGATCCTGAAGAGGCGATTCGTCGTGCGGCAACCATTCTGGCTGAACAACTGGAAGCTTTCGTTGAC
TTACGTGATGTACGTCAGCCTGAAGTGAAAGAAGAGAAACCAGAGTCCAAAACCATCGTTCTTTCGGTCG
GCGAGGCTACTCGCACTCTGACTGAGATCCAGTCCACCGCAGACCGTCAGATCTTCGAAGAGAAGGTCGG
GCCTCTGGTGGGTCGGCTGCGCCTCACGGCTTCGCTCCGTCAAAACGGAGCCAAGACCGCGTATCGCGTC
AACCTAAAACTGGATCAGGCGGACGTCGTTGATTCCGGACTTCCGAAAGTGCGCTACACTCAGGTATGGT
CGCACGACGTGACAATCGTTGCGAATAGCACCGAGGCCTCGCGCAAATCGTTGTACGATTTGACCAAGTC
CCTCGTCGCGACCTCGCAGGTCGAAGATCTTGTCGTCAACCTTGTGCCGCTGGGCCGT**TAA**GCGGCCGCC
ACGCAAAAAACCCCGCTTCGGCGGGGTTTTTTCGC

>TetR-Ptet-Spy-dCas9-AsiA_m2.1

TTAAGACCCACTTTCACATTTAAGTTGTTTTTCTAATCCGCATATGATCAATTCAAGGCCGAATAAGAAG
GCTGGCTCTGCACCTTGGTGATCAAATAATTCGATAGCTTGTCGTAATAATGGCGGCATACTATCAGTAG
TAGGTGTTTCCCTTTCTTCTTTAGCGACTTGATGCTCTTGATCTTCCAATACGCAACCTAAAGTAAAATG
CCCCACAGCGCTGAGTGCATATAATGCATTCTCTAGTGAAAAACCTTGTTGGCATAAAAAGGCTAATTGA
TTTTCGAGAGTTTCATACTGTTTTTCTGTAGGCCGTGTACCTAAATGTACTTTTGCTCCATCGCGATGAC
TTAGTAAAGCACATCTAAAACTTTTAGCGTTATTACGTAAAAAATCTTGCCAGCTTTCCCCTTCTAAAGG
GCAAAAGTGAGTATGGTGCCTATCTAACATCTCAATGGCTAAGGCGTCGAGCAAAGCCCGCTTATTTTTT
ACATGCCAATACAATGTAGGCTGCTCTACACCTAGCTTCTGGGCGAGTTTACGGGTTGTTAAACCTTCGA
TTCCGACCTCATTAAGCAGCTCTAATGCGCTGTTAATCACTTTACTTTTATCTAATCTAGACATCATTAA
TTCCTAATTTTTGTTGACACTCTATCGTTGATAGAGTTATTTTACCACTCCCTATCAGTGATAGAGAAAA
GAATTCAAAAGATCTAAAGAGGAGAAAGGATCT**ATG**GATAAGAAATACTCAATAGGCTTAGCTATCGGCA
CAAATAGCGTCGGATGGGCGGTGATCACTGATGAATATAAGGTTCCGTCTAAAAAGTTCAAGGTTCTGGG
AAATACAGACCGCCACAGTATCAAAAAAAATCTTATAGGGGCTCTTTTATTTGACAGTGGAGAGACAGCG
GAAGCGACTCGTCTCAAACGGACAGCTCGTAGAAGGTATACACGTCGGAAGAATCGTATTTGTTATCTAC
AGGAGATTTTTTCAAATGAGATGGCGAAAGTAGATGATAGTTTCTTTCATCGACTTGAAGAGTCTTTTTT
GGTGGAAGAAGACAAGAAGCATGAACGTCATCCTATTTTTGGAAATATAGTAGATGAAGTTGCTTATCAT
GAGAAATATCCAACTATCTATCATCTGCGAAAAAAATTGGTAGATTCTACTGATAAAGCGGATTTGCGCT
TAATCTATTTGGCCTTAGCGCATATGATTAAGTTTCGTGGTCATTTTTTGATTGAGGGAGATTTAAATCC
TGATAATAGTGATGTGGACAAACTATTTATCCAGTTGGTACAAACCTACAATCAATTATTTGAAGAAAAC
CCTATTAACGCAAGTGGAGTAGATGCTAAAGCGATTCTTTCTGCACGATTGAGTAAATCAAGACGATTAG
AAAATCTCATTGCTCAGCTCCCCGGTGAGAAGAAAAATGGCTTATTTGGGAATCTCATTGCTTTGTCATT
GGGTTTGACCCCTAATTTTAAATCAAATTTTGATTTGGCAGAAGATGCTAAATTACAGCTTTCAAAAGAT
ACTTACGATGATGATTTAGATAATTTATTGGCGCAAATTGGAGATCAATATGCTGATTTGTTTTTGGCAG
CTAAGAATTTATCAGATGCTATTTTACTTTCAGATATCCTAAGAGTAAATACTGAAATAACTAAGGCTCC
CCTATCAGCTTCAATGATTAAACGCTACGATGAACATCATCAAGACTTGACTCTTTTAAAAGCTTTAGTT
CGACAACAACTTCCAGAAAAGTATAAAGAAATCTTTTTTGATCAATCAAAAAACGGATATGCAGGTTATA
TTGATGGGGGAGCTAGCCAAGAAGAATTTTATAAATTTATCAAACCAATTTTAGAAAAAATGGATGGTAC
TGAGGAATTATTGGTGAAACTAAATCGTGAAGATTTGCTGCGCAAGCAACGGACCTTTGACAACGGCTCT
ATTCCCCATCAAATTCACTTGGGTGAGCTGCATGCTATTTTGAGAAGACAAGAAGACTTTTATCCATTTT
TAAAAGACAATCGTGAGAAGATTGAAAAAATCTTGACTTTTCGAATTCCTTATTATGTTGGTCCATTGGC
GCGTGGCAATAGTCGTTTTGCATGGATGACTCGGAAGTCTGAAGAAACAATTACCCCATGGAATTTTGAA
GAAGTTGTCGATAAAGGTGCTTCAGCTCAATCATTTATTGAACGCATGACAAACTTTGATAAAAATCTTC
CAAATGAAAAAGTACTACCAAAACATAGTTTGCTTTATGAGTATTTTACGGTTTATAACGAATTGACAAA
GGTCAAATATGTTACTGAAGGAATGCGAAAACCAGCATTTCTTTCAGGTGAACAGAAGAAAGCCATTGTT
GATTTACTCTTCAAAACAAATCGAAAAGTAACCGTTAAGCAATTAAAAGAAGATTATTTCAAAAAAATAG
AATGTTTTGATAGTGTTGAAATTTCAGGAGTTGAAGATAGATTTAATGCTTCATTAGGTACCTACCATGA
TTTGCTAAAAATTATTAAAGATAAAGATTTTTTGGATAATGAAGAAAATGAAGATATCTTAGAGGATATT
GTTTTAACATTGACCTTATTTGAAGATAGGGAGATGATTGAGGAAAGACTTAAAACATATGCTCACCTCT
TTGATGATAAGGTGATGAAACAGCTTAAACGTCGCCGTTATACTGGTTGGGGACGTTTGTCTCGAAAATT
GATTAATGGTATTAGGGATAAGCAATCTGGCAAAACAATATTAGATTTTTTGAAATCAGATGGTTTTGCC
AATCGCAATTTTATGCAGCTGATCCATGATGATAGTTTGACATTTAAAGAAGACATTCAAAAAGCACAAG
TGTCTGGACAAGGCGATAGTTTACATGAACATATTGCAAATTTAGCTGGTAGCCCTGCTATTAAAAAAGG
TATTTTACAGACTGTAAAAGTTGTTGATGAATTGGTCAAAGTAATGGGGCGGCATAAGCCAGAAAATATC
GTTATTGAAATGGCACGTGAAAATCAGACAACTCAAAAGGGCCAGAAAAATTCGCGAGAGCGTATGAAAC
GAATCGAAGAAGGTATCAAAGAATTAGGAAGTCAGATTCTTAAAGAGCATCCTGTTGAAAATACTCAATT

GCAAAATGAAAAGCTCTATCTCTATTATCTCCAAAATGGAAGAGACATGTATGTGGACCAAGAATTAGAT
ATTAATCGTTTAAGTGATTATGATGTCGATGCCATTGTTCCACAAAGTTTCCTTAAAGACGATTCAATAG
ACAATAAGGTCTTAACGCGTTCTGATAAAAATCGTGGTAAATCGGATAACGTTCCAAGTGAAGAAGTAGT
CAAAAAGATGAAAAACTATTGGAGACAACTTCTAAACGCCAAGTTAATCACTCAACGTAAGTTTGATAAT
TTAACGAAAGCTGAACGTGGAGGTTTGAGTGAACTTGATAAAGCTGGTTTTATCAAACGCCAATTGGTTG
AAACTCGCCAAATCACTAAGCATGTGGCACAAATTTTGGATAGTCGCATGAATACTAAATACGATGAAAA
TGATAAACTTATTCGAGAGGTTAAAGTGATTACCTTAAAATCTAAATTAGTTTCTGACTTCCGAAAAGAT
TTCCAATTCTATAAAGTACGTGAGATTAACAATTACCATCATGCCCATGATGCGTATCTAAATGCCGTCG
TTGGAACTGCTTTGATTAAGAAATATCCAAAACTTGAATCGGAGTTTGTCTATGGTGATTATAAAGTTTA
TGATGTTCGTAAAATGATTGCTAAGTCTGAGCAAGAAATAGGCAAAGCAACCGCAAAATATTTCTTTTAC
TCTAATATCATGAACTTCTTCAAAACAGAAATTACACTTGCAAATGGAGAGATTCGCAAACGCCCTCTAA
TCGAAACTAATGGGGAAACTGGAGAAATTGTCTGGGATAAAGGGCGAGATTTTGCCACAGTGCGCAAAGT
ATTGTCCATGCCCCAAGTCAATATTGTCAAGAAAACAGAAGTACAGACAGGCGGATTCTCCAAGGAGTCA
ATTTTACCAAAAAGAAATTCGGACAAGCTTATTGCTCGTAAAAAAGACTGGGATCCAAAAAAATATGGTG
GTTTTGATAGTCCAACGGTAGCTTATTCAGTCCTAGTGGTTGCTAAGGTGGAAAAAGGGAAATCGAAGAA
GTTAAAATCCGTTAAAGAGTTACTAGGGATCACAATTATGGAAAGAAGTTCCTTTGAAAAAAATCCGATT
GACTTTTTAGAAGCTAAAGGATATAAGGAAGTTAAAAAAGACTTAATCATTAAACTACCTAAATATAGTC
TTTTTGAGTTAGAAAACGGTCGTAAACGGATGCTGGCTAGTGCCGGAGAATTACAAAAAGGAAATGAGCT
GGCTCTGCCAAGCAAATATGTGAATTTTTTATATTTAGCTAGTCATTATGAAAAGTTGAAGGGTAGTCCA
GAAGATAACGAACAAAAACAATTGTTTGTGGAGCAGCATAAGCATTATTTAGATGAGATTATTGAGCAAA
TCAGTGAATTTTCTAAGCGTGTTATTTTAGCAGATGCCAATTTAGATAAAGTTCTTAGTGCATATAACAA
ACATAGAGACAAACCAATACGTGAACAAGCAGAAAATATTATTCATTTATTTACGTTGACGAATCTTGGA
GCTCCCGCTGCTTTTAAATATTTTGATACAACAATTGATCGTAAACGATATACGTCTACAAAAGAAGTTT
TAGATGCCACTCTTATCCATCAATCCATCACTGGTCTTTATGAAACACGCATTGATTTGAGTCAGCTAGG
AGGTGACTGCGCGGGTGGCGGAGGAAGTGGGGGCGGTGGTAGCATGAATAAAAACATTGATACAGTTCGT
GAAATTATTACTGTTGCATCTATTTTGATTAAATTTTCCAGAGAAGATATTGTTGAGAATCGTGCTAATT
TTATTGCATTTCTAAATGAGATTGGAGTAACGCATGAAGGTAGAAAGTTAAATCGCAATTCATTCCGTAA
AATTATTTCTAAATTAACTCAAGAAGATAAGAAAACCCTCATCGACGAATTCAACGAGGGTTTTGAGGGT
GTATATCGATATCTAGAGATGTATACGAACAAA**TAA**CTCGAGTAAGGATCTCCAGGCATCAAATAAAACG
AAAGGCTCAGTCGAAAGACTGGGCCTTTCGTTTTATCTGTTGTTTGTCGGTGAACGCTCTCTACTAGAGT
CACACTGGCTCACCTTCGGGTGGGCCTTTCTGCGTTTATA

>BBa_J23107-RBS-PspF-λN22-BBa_B1002
TTTACGGCTAGCTCAGCCCTAGGTATTATGCTAGCGAATTCATTAAAGAGGAGAAAGGTACC**ATG**GGGCC
CATGGCAGAATACAAAGATAATTTACTTGGTGAGGCGAACAGCTTTCTCGAAGTGCTGGAACAGGTTTCG
CATCTCGCACCGCTGGACAAACCGGTGCTCATCATCGGCGAACGCGGCACCGGTAAAGAGCTGATTGCCA
GCCGCCTGCATTATCTCTCCTCCCGTTGGCAAGGGCCGTTTATTTCCCTTAACTGCGCGGCGTTAAATGA
AAATCTGCTGGATTCCGAACTGTTTGGTCACGAAGCGGGGGCGTTTACCGGTGCGCAAAAACGTCATCCA
GGGAGATTTGAACGTGCCGACGGCGGTACGCTATTTCTTGATGAACTCGCTACGGCACCGATGATGGTGC
AGGAGAAATTATTGCGCGTGATTGAGTACGGTGAACTGGAGCGCGTTGGCGGCAGCCAACCATTGCAGGT
GAATGTGCGGTTGGTATGCGCGACGAATGCCGATCTCCCGGCGATGGTCAATGAAGGCACTTTTCGCGCT
GACCTGCTCGACCGACTGGCTTTTGATGTTGTACAACTGCCACCACTGCGCGAGCGCGAAAGCGACATAA
TGTTGATGGCAGAATACTTTGCCATCCAGATGTGTCGGGAAATCAAGCTGCCTCTGTTCCCGGGGTTTAC
GGAGCGCGCCAGAGAAACATTGCTGAATTATCGTTGGCCGGGAAATATTCGTGAATTGAAAAACGTGGTG
GAACGTTCAGTGTATCGCCACGGCACCAGCGATTATCCGCTTGATGACATCATTATTGATCCCTTTAAAC

GGCGTCCGCCTGAAGACGCTATCGCCGTTTCAGAAACCACCTCGCTTCCAACACTGCCGCTGGATTTACG
TGAGTTTCAGATGCAGCAGGAAAAAGAGTTGCTGCAACTCAGTTTGCAAATGAATGCACGCACACGCCGC
CGCGAACGTCGCGCAGAGAAACAGGCTCAATGGAAAGCAGCAAAT**TAA**GCGGCCGCCACGCAAAAAACCC
CGCTTCGGCGGGGTTTTTTCGC

>BBa_J23107-RBS-MCP-TetD-BBa_B1002
TTTACGGCTAGCTCAGCCCTAGGTATTATGCTAGCGAATTCATTAAAGAGGAGAAAGGTACC**ATG**GGGCC
CGCTTCTAACTTTACTCAGTTCGTTCTCGTCGACAATGGCGGAACTGGCGACGTGACTGTCGCCCCAAGC
AACTTCGCTAACGGGATCGCTGAATGGATCAGCTCTAACTCGCGTTCACAGGCTTACAAAGTAACCTGTA
GCGTTCGTCAGAGCTCTGCGCAGAATCGCAAATACACCATCAAAGTCGAGGTGCCTAAAGGCGCCTGGCG
TTCGTACTTAAATATGGAACTAACCATTCCAATTTTCGCCACGAATTCCGACTGCGAGCTTATTGTTAAG
GCAATGCAAGGTCTCCTAAAAGATGGAAACCCGATTCCCTCAGCAATCGCAGCAAACTCCGGCATCTACG
GTGGCGGAGGTAGCATGTATATTGAACAGCATTCTCGCTATCAAAATAAAGCTAATAACATCCAATTAAG
ATATGATGATAAGCAGTTTCATACAACGGTTATCAAAGATGTTCTATTATGGATTGAACATAATTTAGAT
CAGTCTTTACTGCTTGATGATGTGGCGAATAAAGCGGGTTATACCAAGTGGTATTTTCAGCGGCTGTTCA
AAAAAGTAACAGGGGTCACACTGGCTAGCTATATTCGTGCTCGTCGTTTGACGAAAGCGGCTGTTGAGTT
GAGGTTGACGAAAAAAACTATCCTTGAGATCGCATTAAAATATCAATTTGATTCCCAACAATCTTTTACA
CGTCGATTTAAGTACATTTTTAAGGTTACACCAAGTTATTATCGGCGTAATAAATTATGGGAATTGGAGG
CAATGCAC**TAA**GCGGCCGCCACGCAAAAAACCCCGCTTCGGCGGGGTTTTTTCGC

>BBa_J23107-RBS-MCP-Crl-BBa_B1002
TTTACGGCTAGCTCAGCCCTAGGTATTATGCTAGCGAATTCATTAAAGAGGAGAAAGGTACC**ATG**GGGCC
CGCTTCTAACTTTACTCAGTTCGTTCTCGTCGACAATGGCGGAACTGGCGACGTGACTGTCGCCCCAAGC
AACTTCGCTAACGGGATCGCTGAATGGATCAGCTCTAACTCGCGTTCACAGGCTTACAAAGTAACCTGTA
GCGTTCGTCAGAGCTCTGCGCAGAATCGCAAATACACCATCAAAGTCGAGGTGCCTAAAGGCGCCTGGCG
TTCGTACTTAAATATGGAACTAACCATTCCAATTTTCGCCACGAATTCCGACTGCGAGCTTATTGTTAAG
GCAATGCAAGGTCTCCTAAAAGATGGAAACCCGATTCCCTCAGCAATCGCAGCAAACTCCGGCATCTACG
GTGGCGGAGGTAGCATGACGTTACCGAGTGGACACCCGAAGAGCAGATTGATCAAAAAATTTACCGCACT
AGGCCCGTATATTCGTGAAGGTAAGTGCAAAGATAATCGATTCTTTTTCGATTGTCTGGCTGTATGCGTC
AACGTGAAACCGGCACCGGAAGTGCGTGAATTCTGGGGCTGGTGGATGGAGCTGGAAGCGCAGGAATCCC
GTTTTACCTACAGTTACCAGTTTGGTCTGTTCGATAAAGCAGGCGACTGGAAGAGTGTTCCGGTAAAAGA
CACTGAAGTGGTTGAACGACTGGAGCACACCCTGCGTGAGTTTCACGAGAAGCTGCGTGAACTGCTGACG
ACGCTGAATCTGAAGCTGGAACCGGCGGATGATTTTCGTGACGAGCCGGTGAAGTTAACGGCG**TAA**GCGG
CCGCCACGCAAAAAACCCCGCTTCGGCGGGGTTTTTTCGC

>BBa_J23107-RBS-MCP-SoxS-TetD-BBa_B1002
TTTACGGCTAGCTCAGCCCTAGGTATTATGCTAGCGAATTCATTAAAGAGGAGAAAGGTACC**ATG**GGGCC
CGCTTCTAACTTTACTCAGTTCGTTCTCGTCGACAATGGCGGAACTGGCGACGTGACTGTCGCCCCAAGC
AACTTCGCTAACGGGATCGCTGAATGGATCAGCTCTAACTCGCGTTCACAGGCTTACAAAGTAACCTGTA
GCGTTCGTCAGAGCTCTGCGCAGAATCGCAAATACACCATCAAAGTCGAGGTGCCTAAAGGCGCCTGGCG
TTCGTACTTAAATATGGAACTAACCATTCCAATTTTCGCCACGAATTCCGACTGCGAGCTTATTGTTAAG
GCAATGCAAGGTCTCCTAAAAGATGGAAACCCGATTCCCTCAGCAATCGCAGCAAACTCCGGCATCTACG
GTGGCGGAGGTAGCATGTCCCATCAGAAAATTATTCAGGATCTTATCGCATGGATTGACGAGCATATTGA
CCAGCCGCTTAACATTGATGTAGTCGCAAAAAAATCAGGCTATTCAAAGTGGTACTTGCAACGAATGTTC
CGCACGGTGACGCATCAGACGCTTGGCGATTACATTCGCCAACGCCGCCTGTTACTGGCCGCCGTTGAGT

TGCGCACCACCGAGCGTCCGATTTTTGATATCGCAATGGACCTGGGTTATGTCTCGCAGCAGACCTTCTC
CCGCGTTTTCGCGCGGCAGTTTGATCGCACTCCCGCGGATTATCGCCACCGCCTGGGTGGCGGAGGTAGC
ATGTATATTGAACAGCATTCTCGCTATCAAAATAAAGCTAATAACATCCAATTAAGATATGATGATAAGC
AGTTTCATACAACGGTTATCAAAGATGTTCTATTATGGATTGAACATAATTTAGATCAGTCTTTACTGCT
TGATGATGTGGCGAATAAAGCGGGTTATACCAAGTGGTATTTTCAGCGGCTGTTCAAAAAAGTAACAGGG
GTCACACTGGCTAGCTATATTCGTGCTCGTCGTTTGACGAAAGCGGCTGTTGAGTTGAGGTTGACGAAAA
AAACTATCCTTGAGATCGCATTAAAATATCAATTTGATTCCCAACAATCTTTTACACGTCGATTTAAGTA
CATTTTTAAGGTTACACCAAGTTATTATCGGCGTAATAAATTATGGGAATTGGAGGCAATGCAC**TAA**GCG
GCCGCCACGCAAAAAACCCCGCTTCGGCGGGTTTTTCGC

>BBa_J23107-RBS-αNTD-MCP-SoxS-BBa_B1002
TTTACGGCTAGCTCAGCCCTAGGTATTATGCTAGCGAATTCATTAAAGAGGAGAAAGGTACC**ATG**GGGCC
CATGCAGGGTTCTGTGACAGAGTTTCTAAAACCGCGCCTGGTTGATATCGAGCAAGTGAGTTCGACGCAC
GCCAAGGTGACCCTTGAGCCTTTAGAGCGTGGCTTTGGCCATACTCTGGGTAACGCACTGCGCCGTATTC
TGCTCTCATCGATGCCGGGTTGCGCGGTGACCGAGGTTGAGATTGATGGTGTACTACATGAGTACAGCAC
CAAAGAAGGCGTTCAGGAAGATATCCTGGAAATCCTGCTCAACCTGAAAGGGCTGGCGGTGAGAGTTCAG
GGCAAAGATGAAGTTATTCTTACCTTGAATAAATCTGGCATTGGCCCTGTGACTGCAGCCGATATCACCC
ACGACGGTGATGTCGAAATCGTCAAGCCGCAGCACGTGATCTGCCACCTGACCGATGAGAACGCGTCTAT
TAGCATGCGTATCAAAGTTCAGCGCGGTCGTGGTTATGTGCCGGCTTCTACCCGAATTCATTCGGAAGAA
GATGAGCGCCCAATCGGCCGTCTGCTGGTCGACGCATGCTACAGCCCTGTGGAGCGTATTGCCTACAATG
TTGAAGCAGCGCGTGTAGAACAGCGTACCGACCTGGACAAGCTGGTCATCGAAATGGAAACCAACGGCAC
AATCGATCCTGAAGAGGCGATTCGTCGTGCGGCAACCATTCTGGCTGAACAACTGGAAGCTTTCGTTGAC
TTACGTGATGTACGTCAGCCTGAAGTGAAAGAAGAGAAACCAGAGGCTTCTAACTTTACTCAGTTCGTTC
TCGTCGACAATGGCGGAACTGGCGACGTGACTGTCGCCCCAAGCAACTTCGCTAACGGGATCGCTGAATG
GATCAGCTCTAACTCGCGTTCACAGGCTTACAAAGTAACCTGTAGCGTTCGTCAGAGCTCTGCGCAGAAT
CGCAAATACACCATCAAAGTCGAGGTGCCTAAAGGCGCCTGGCGTTCGTACTTAAATATGGAACTAACCA
TTCCAATTTTCGCCACGAATTCCGACTGCGAGCTTATTGTTAAGGCAATGCAAGGTCTCCTAAAAGATGG
AAACCCGATTCCCTCAGCAATCGCAGCAAACTCCGGCATCTACGGTGGCGGAGGTAGCATGTCCCATCAG
AAAATTATTCAGGATCTTATCGCATGGATTGACGAGCATATTGACCAGCCGCTTAACATTGATGTAGTCG
CAAAAAAATCAGGCTATTCAAAGTGGTACTTGCAACGAATGTTCCGCACGGTGACGCATCAGACGCTTGG
CGATTACATTCGCCAACGCCGCCTGTTACTGGCCGCCGTTGAGTTGCGCACCACCGAGCGTCCGATTTTT
GATATCGCAATGGACCTGGGTTATGTCTCGCAGCAGACCTTCTCCCGCGTTTTCGCGCGGCAGTTTGATC
GCACTCCCGCGGATTATCGCCACCGCCTG**TAA**GCGGCCGCCACGCAAAAAACCCCGCTTCGGCGGGTTT
TTTCGC

>BBa_J23107-RBS-MCP-SYNZIP6-BBa_B1002_
BBa_1002-SYNZIP5-αNTD-RBS-BBa_J23107
TTTACGGCTAGCTCAGCCCTAGGTATTATGCTAGCGAATTCATTAAAGAGGAGAAAGGTACC**ATG**GGGCC
CGCTTCTAACTTTACTCAGTTCGTTCTCGTCGACAATGGCGGAACTGGCGACGTGACTGTCGCCCCAAGC
AACTTCGCTAACGGGATCGCTGAATGGATCAGCTCTAACTCGCGTTCACAGGCTTACAAAGTAACCTGTA
GCGTTCGTCAGAGCTCTGCGCAGAATCGCAAATACACCATCAAAGTCGAGGTGCCTAAAGGCGCCTGGCG
TTCGTACTTAAATATGGAACTAACCATTCCAATTTTCGCCACGAATTCCGACTGCGAGCTTATTGTTAAG
GCAATGCAAGGTCTCCTAAAAGATGGAAACCCGATTCCCTCAGCAATCGCAGCAAACTCCGGCATCTACG
GTGGCGGAGGTAGCCAAAAAGTTGCGCAGCTGAAAAACCGTGTTGCGTACAAACTGAAAGAAAACGCGAA
GCTGGAGAACATCGTGGCGCGTCTGGAAAACGACAATGCGAACCTGGAGAAAGACATTGCGAATCTCGAA

AAGGACATCGCAAATCTGGAACGTGACGTTGCGCGT**TAA**GCGGCCGCCACGCAAAAAACCCCGCTTCGGC
GGGGTTTTTTCGCCCTAGGGATATATTCCGCTTCCTCGCTCACTGACTCGCTACGCTCGGTCGTTCGACT
GCGGCGAGCGGAAATGGCTTACGAACGGGGCGGAGATTTCCTGCCCGGGGCGAAAAAACCCCGCCGAAGC
GGGGTTTTTTGCGTGGGCGCGCC**TTA**CTCGAATTTGTGAGCCGCCAGTTCGAATTCCAGTTCCGCTTTTG
CGAATTTCAGGTGTTCCTTCAGGTTTTTGAGTTCAGCGTTACGCTCTTCCAGCTCCTGGATGTAGTTTTT
CAGTTCTTTAACGGTGTTCTCTGGTTTCTCTTCTTTCACTTCAGGCTGACGTACATCACGTAAGTCAACG
AAAGCTTCCAGTTGTTCAGCCAGAATGGTTGCCGCACGACGAATCGCCTCTTCAGGATCGATTGTGCCGT
TGGTTTCCATTTCGATGACCAGCTTGTCCAGGTCGGTACGCTGTTCTACACGCGCTGCTTCAACATTGTA
GGCAATACGCTCCACAGGGCTGTAGCATGCGTCGACCAGCAGACGGCCGATTGGGCGCTCATCTTCTTCC
GAATGAATTCGGGTAGAAGCCGGCACATAACCACGACCGCGCTGAACTTTGATACGCATGCTAATAGACG
CGTTCTCATCGGTCAGGTGGCAGATCACGTGCTGCGGCTTGACGATTTCGACATCACCGTCGTGGGTGAT
ATCGGCTGCAGTCACAGGGCCAATGCCAGATTTATTCAAGGTAAGAATAACTTCATCTTTGCCCTGAACT
CTCACCGCCAGCCCTTTCAGGTTGAGCAGGATTTCCAGGATATCTTCCTGAACGCCTTCTTTGGTGCTGT
ACTCATGTAGTACACCATCAATCTCAACCTCGGTCACCGCGCAACCCGGCATCGATGAGAGCAGAATACG
GCGCAGTGCGTTACCCAGAGTATGGCCAAAGCCACGCTCTAAAGGCTCAAGGGTCACCTTGGCGTGCGTC
GAACTCACTTGCTCGATATCAACCAGGCGCGGTTTTAGAAACTCTGTCACAGAACCCTG**CAT**GGTACCTT
TCTCCTCTTTAATGAATTCGCTAGCATAATACCTAGGGCTGAGCTAGCCGTAAA

>BBa_J23107-RBS-MCP-SoxS-SYNZIP6-BBa_B1002_
BBa_1002-SYNZIP5-αNTD-RBS-BBa_J23107
TTTACGGCTAGCTCAGCCCTAGGTATTATGCTAGCGAATTCATTAAAGAGGAGAAAGGTACC**ATG**GGGCC
CGCTTCTAACTTTACTCAGTTCGTTCTCGTCGACAATGGCGGAACTGGCGACGTGACTGTCGCCCCAAGC
AACTTCGCTAACGGGATCGCTGAATGGATCAGCTCTAACTCGCGTTCACAGGCTTACAAAGTAACCTGTA
GCGTTCGTCAGAGCTCTGCGCAGAATCGCAAATACACCATCAAAGTCGAGGTGCCTAAAGGCGCCTGGCG
TTCGTACTTAAATATGGAACTAACCATTCCAATTTTCGCCACGAATTCCGACTGCGAGCTTATTGTTAAG
GCAATGCAAGGTCTCCTAAAAGATGGAAACCCGATTCCCTCAGCAATCGCAGCAAACTCCGGCATCTACG
GTGGCGGAGGTAGCATGTCCCATCAGAAAATTATTCAGGATCTTATCGCATGGATTGACGAGCATATTGA
CCAGCCGCTTAACATTGATGTAGTCGCAAAAAAATCAGGCTATTCAAAGTGGTACTTGCAACGAATGTTC
CGCACGGTGACGCATCAGACGCTTGGCGATTACATTCGCCAACGCCGCCTGTTACTGGCCGCCGTTGAGT
TGCGCACCACCGAGCGTCCGATTTTTGATATCGCAATGGACCTGGGTTATGTCTCGCAGCAGACCTTCTC
CCGCGTTTTCGCGCGGCAGTTTGATCGCACTCCCGCGGATTATCGCCACCGCCTGGGTGGCGGAGGTAGC
CAAAAAGTTGCGCAGCTGAAAAACCGTGTTGCGTACAAACTGAAAGAAAACGCGAAGCTGGAGAACATCG
TGGCGCGTCTGGAAAACGACAATGCGAACCTGGAGAAAGACATTGCGAATCTCGAAAAGGACATCGCAAA
TCTGGAACGTGACGTTGCGCGT**TAA**GCGGCCGCCACGCAAAAAACCCCGCTTCGGCGGGGTTTTTTCGCC
CTAGGGATATATTCCGCTTCCTCGCTCACTGACTCGCTACGCTCGGTCGTTCGACTGCGGCGAGCGGAAA
TGGCTTACGAACGGGGCGGAGATTTCCTGCCCGGGGCGAAAAAACCCCGCCGAAGCGGGGTTTTTTGCGT
GGGCGCGCC**TTA**CTCGAATTTGTGAGCCGCCAGTTCGAATTCCAGTTCCGCTTTTGCGAATTTCAGGTGT
TCCTTCAGGTTTTTGAGTTCAGCGTTACGCTCTTCCAGCTCCTGGATGTAGTTTTTCAGTTCTTTAACGG
TGTTCTCTGGTTTCTCTTCTTTCACTTCAGGCTGACGTACATCACGTAAGTCAACGAAAGCTTCCAGTTG
TTCAGCCAGAATGGTTGCCGCACGACGAATCGCCTCTTCAGGATCGATTGTGCCGTTGGTTTCCATTTCG
ATGACCAGCTTGTCCAGGTCGGTACGCTGTTCTACACGCGCTGCTTCAACATTGTAGGCAATACGCTCCA
CAGGGCTGTAGCATGCGTCGACCAGCAGACGGCCGATTGGGCGCTCATCTTCTTCCGAATGAATTCGGGT
AGAAGCCGGCACATAACCACGACCGCGCTGAACTTTGATACGCATGCTAATAGACGCGTTCTCATCGGTC
AGGTGGCAGATCACGTGCTGCGGCTTGACGATTTCGACATCACCGTCGTGGGTGATATCGGCTGCAGTCA
CAGGGCCAATGCCAGATTTATTCAAGGTAAGAATAACTTCATCTTTGCCCTGAACTCTCACCGCCAGCCC

TTTCAGGTTGAGCAGGATTTCCAGGATATCTTCCTGAACGCCTTCTTTGGTGCTGTACTCATGTAGTACA
CCATCAATCTCAACCTCGGTCACCGCGCAACCCGGCATCGATGAGAGCAGAATACGGCGCAGTGCGTTAC
CCAGAGTATGGCCAAAGCCACGCTCTAAAGGCTCAAGGGTCACCTTGGCGTGCGTCGAACTCACTTGCTC
GATATCAACCAGGCGCGGTTTTAGAAACTCTGTCACAGAACCCTG**CAT**GGTACCTTTCTCCTCTTTAATG
AATTCGCTAGCATAATACCTAGGGCTGAGCTAGCCGTAAA

>TetR-Ptet-Sth1-dCas9-AsiA_m2.1
TTAAGACCCACTTTCACATTTAAGTTGTTTTTCTAATCCGCATATGATCAATTCAAGGCCGAATAAGAAG
GCTGGCTCTGCACCTTGGTGATCAAATAATTCGATAGCTTGTCGTAATAATGGCGGCATACTATCAGTAG
TAGGTGTTTCCCTTTCTTCTTTAGCGACTTGATGCTCTTGATCTTCCAATACGCAACCTAAAGTAAAATG
CCCCACAGCGCTGAGTGCATATAATGCATTCTCTAGTGAAAAACCTTGTTGGCATAAAAAGGCTAATTGA
TTTTCGAGAGTTTCATACTGTTTTTCTGTAGGCCGTGTACCTAAATGTACTTTTGCTCCATCGCGATGAC
TTAGTAAAGCACATCTAAAACTTTTAGCGTTATTACGTAAAAAATCTTGCCAGCTTTCCCCTTCTAAAGG
GCAAAAGTGAGTATGGTGCCTATCTAACATCTCAATGGCTAAGGCGTCGAGCAAAGCCCGCTTATTTTTT
ACATGCCAATACAATGTAGGCTGCTCTACACCTAGCTTCTGGGCGAGTTTACGGGTTGTTAAACCTTCGA
TTCCGACCTCATTAAGCAGCTCTAATGCGCTGTTAATCACTTTACTTTTATCTAATCTAGACATCATTAA
TTCCTAATTTTTGTTGACACTCTATCGTTGATAGAGTTATTTTACCACTCCCTATCAGTGATAGAGAAAA
GAATTCAAAAGATCTAAAGAGGAGAAAGGATCT**ATG**AGTGACTTAGTTTTAGGACTTGCTATCGGTATAG
GTTCTGTTGGTGTGGGTATCCTTAACAAAGTGACAGGAGAAATTATCCATAAAAACTCACGCATCTTCCC
AGCAGCTCAAGCGGAAAATAACCTAGTACGTAGAACGAATCGTCAAGGAAGACGCTTGGCACGACGTAAA
AAACATCGTAGAGTTCGTTTAAATCGTCTATTTGAGGAAAGTGGATTAATCACTGATTTTACGAAGATTT
CAATTAATCTTAACCCATATCAATTACGAGTTAAGGGCTTGACCGATGAATTGTCTAATGAAGAACTGTT
TATCGCTCTTAAAAATATGGTGAAACACCGTGGGATTAGTTACCTCGATGATGCTAGTGATGACGGAAAT
TCATCAGTAGGAGACTATGCACAAATTGTTAAGGAAAATAGTAAACAATTAGAAACTAAGACACCGGGAC
AGATACAGTTGGAACGCTACCAAACATATGGTCAATTACGTGGTGATTTTACTGTTGAGAAAGATGGCAA
AAAACATCGCTTGATTAATGTCTTTCCAACATCAGCTTATCGTTCAGAAGCCTTAAGGATACTGCAAACT
CAACAAGAATTTAATCCACAGATTACAGATGAATTTATTAATCGTTATCTCGAAATTTTAACTGGAAAAC
GGAAATATTATCATGGACCCGGAAATGAAAAGTCACGGACTGATTATGGTCGTTACAGAACGAGTGGAGA
AACTTTAGACAATATTTTTGGAATTCTAATTGGGAAATGTACATTTTATCCAGACGAGTTTAGAGCAGCA
AAAGCTTCCTACACGGCTCAAGAATTCAATTTGCTAAATGATTTGAACAATCTAACAGTTCCTACTGAAA
CCAAAAAGTTGAGCAAAGAACAGAAGAATCAAATCATTAATTATGTCAAAAATGAAAAGGCAATGGGGCC
AGCGAAACTTTTTAAATATATCGCTAAGTTACTTTCTTGTGATGTTGCAGATATCAAGGGATACCGTATC
GACAAATCAGGTAAGGCTGAGATTCATACTTTCGAAGCCTATCGAAAAATGAAAACGCTTGAAACCTTAG
ATATTGAGCAAATGGATAGAGAAACGCTTGATAAATTAGCCTATGTCTTAACATTAAACACTGAGAGGGA
AGGTATTCAAGAAGCCTTAGAACATGAATTTGCTGATGGTAGCTTTAGCCAGAAGCAAGTTGACGAATTG
GTTCAATTCCGCAAAGCAAATAGTTCCATTTTTGGAAAAGGATGGCATAATTTTTCTGTCAAACTGATGA
TGGAGTTAATTCCAGAATTGTATGAGACGTCAGAAGAGCAAATGACTATCCTAACACGACTTGGAAAACA
AAAAACGACTTCGTCTTCAAATAAAACAAAATATATAGATGAAAAACTATTAACTGAAGAAATCTATAAT
CCTGTTGTTGCTAAGTCTGTTCGCCAGGCTATAAAAATCGTAAATGCGGCGATTAAAGAATACGGAGACT
TTGACAATATTGTCATCGAAATGGCTCGTGAAACAAATGAAGATGATGAAAAGAAAGCTATTCAAAAGAT
TCAAAAAGCCAACAAAGATGAAAAAGATGCAGCAATGCTTAAGGCTGCTAACCAATATAATGGAAAGGCT
GAATTACCACATAGTGTTTTCCACGGTCATAAGCAATTAGCGACTAAAATCCGCCTTTGGCATCAGCAAG
GAGAACGTTGCCTTTATACTGGTAAGACAATCTCAATCCATGATTTGATAAATAATTCTAATCAGTTTGA
AGTAGATGCTATTTTACCTCTTTCTATCACATTCGATGATAGCCTTGCAAATAAGGTTTTGGTTTATGCA
ACTGCTAACCAAGAAAAAGGACAACGAACACCTTATCAGGCTTTAGATAGTATGGATGATGCGTGGTCTT

TCCGTGAATTAAAAGCTTTTGTACGTGAGTCAAAAACACTTTCGAACAAGAAAAAAGAATACCTCCTTAC
AGAAGAAGATATTTCAAAGTTTGATGTTCGAAAGAAATTTATTGAACGAAATCTTGTAGATACAAGATAC
GCTTCAAGAGTTGTCCTCAATGCCCTTCAAGAACACTTTAGAGCTCACAAGATTGATACAAAAGTTTCCG
TGGTTCGTGGCCAATTTACATCTCAATTGAGACGCCATTGGGGAATTGAGAAGACTCGTGATACTTATCA
TCACCATGCTGTCGATGCATTGATTATTGCCGCCTCAAGTCAGTTGAATTTGTGGAAAAAACAAAAGAAT
ACCCTTGTAAGTTATTCAGAAGACCAACTCCTTGATATTGAAACAGGTGAACTTATTAGTGATGATGAGT
ACAAGGAATCTGTGTTCAAAGCCCCTTATCAACATTTTGTTGATACATTGAAGAGTAAAGAATTTGAAGA
CAGTATCTTATTCTCTTATCAAGTGGATTCTAAGTTTAATCGTAAAATATCAGATGCCACTATTTATGCG
ACAAGACAGGCTAAAGTAGGAAAAGATAAGGCGGATGAAACTTATGTCTTAGGGAAAATCAAAGATATCT
ATACTCAGGATGGTTATGATGCCTTTATGAAGATTTATAAGAAGGATAAGTCAAAATTCCTCATGTATCG
TCACGACCCACAAACCTTTGAGAAAGTTATCGAGCCAATTTTAGAGAACTATCCTAATAAGCAAATAAAT
GAAAAAGGGAAAGAGGTACCATGTAATCCTTTCCTAAAATATAAAGAAGAACATGGCTATATTCGTAAAT
ATAGTAAAAAAGGCAATGGTCCTGAAATCAAGAGTCTTAAATACTATGATAGTAAGCTAGGCAATCATAT
TGATATTACTCCGAAGGATAGTAACAATAAAGTTGTCTTACAGTCAGTTTCTCCATGGAGAGCGGATGTC
TATTTCAATAAGACTACTGGAAAATACGAAATCCTCGGATTGAAATATGCTGATTTACAATTTGAAAAAG
GGACAGGAACATATAAGATTTCCCAGGAAAAATACAATGACATTAAGAAAAAAGAGGGTGTAGATTCTGA
TTCAGAATTCAAGTTTACACTTTATAAAAATGATTTGTTACTCGTTAAAGATACAGAAACAAAAGAACAA
CAGCTTTTCCGTTTTCTTTCTCGAACTATGCCTAAACAAAAGCATTATGTTGAATTAAAACCTTATGATA
AACAGAAATTTGAAGGAGGTGAGGCGTTAATTAAAGTGTTGGGTAACGTTGCTAATAGTGGTCAATGCAA
AAAAGGGCTAGGAAAATCAAATATTTCTATTTATAAGGTAAGAACAGATGTCCTAGGAAATCAGCATATC
ATCAAAAATGAGGGTGATAAGCCTAAGCTAGATTTTTGCGCGGGTGGCGGAGGAAGTGGGGGCGGTGGTA
GCATGAATAAAAACATTGATACAGTTCGTGAAATTATTACTGTTGCATCTATTTTGATTAAATTTTCCAG
AGAAGATATTGTTGAGAATCGTGCTAATTTTATTGCATTTCTAAATGAGATTGGAGTAACGCATGAAGGT
AGAAAGTTAAATCGCAATTCATTCCGTAAAATTATTTCTAAATTAACTCAAGAAGATAAGAAAACCCTCA
TCGACGAATTCAACGAGGGTTTTGAGGGTGTATATCGATATCTAGAGATGTATACGAACAAA**TAA**CTCGA
GTAAGGATCTCCAGGCATCAAATAAAACGAAAGGCTCAGTCGAAAGACTGGGCCTTTCGTTTTATCTGTT
GTTTGTCGGTGAACGCTCTCTACTAGAGTCACACTGGCTCACCTTCGGGTGGGCCTTTCTGCGTTTATA

>TetR-Ptet-Lb-dCas12a-AsiA_m2.1
TTAAGACCCACTTTCACATTTAAGTTGTTTTTCTAATCCGCATATGATCAATTCAAGGCCGAATAAGAAG
GCTGGCTCTGCACCTTGGTGATCAAATAATTCGATAGCTTGTCGTAATAATGGCGGCATACTATCAGTAG
TAGGTGTTTCCCTTTCTTCTTTAGCGACTTGATGCTCTTGATCTTCCAATACGCAACCTAAAGTAAAATG
CCCCACAGCGCTGAGTGCATATAATGCATTCTCTAGTGAAAAACCTTGTTGGCATAAAAAGGCTAATTGA
TTTTCGAGAGTTTCATACTGTTTTTCTGTAGGCCGTGTACCTAAATGTACTTTTGCTCCATCGCGATGAC
TTAGTAAAGCACATCTAAAACTTTTAGCGTTATTACGTAAAAAATCTTGCCAGCTTTCCCCTTCTAAAGG
GCAAAAGTGAGTATGGTGCCTATCTAACATCTCAATGGCTAAGGCGTCGAGCAAAGCCCGCTTATTTTTT
ACATGCCAATACAATGTAGGCTGCTCTACACCTAGCTTCTGGGCGAGTTTACGGGTTGTTAAACCTTCGA
TTCCGACCTCATTAAGCAGCTCTAATGCGCTGTTAATCACTTTACTTTTATCTAATCTAGACATCATTAA
TTCCTAATTTTTGTTGACACTCTATCGTTGATAGAGTTATTTTACCACTCCCTATCAGTGATAGAGAAAA
GAATTCAAAAGATCTAAAGAGGAGAAAGGATCT**ATG**AGCAAGCTGGAGAAGTTTACAAACTGCTACTCCC
TGTCTAAGACCCTGAGGTTCAAGGCCATCCCTGTGGGCAAGACCCAGGAGAACATCGACAATAAGCGGCT
GCTGGTGGAGGACGAGAAGAGAGCCGAGGATTATAAGGGCGTGAAGAAGCTGCTGGATCGCTACTATCTG
TCTTTTATCAACGACGTGCTGCACAGCATCAAGCTGAAGAATCTGAACAATTACATCAGCCTGTTCCGGA
AGAAAACCAGAACCGAGAAGGAGAATAAGGAGCTGGAGAACCTGGAGATCAATCTGCGGAAGGAGATCGC
CAAGGCCTTCAAGGGCAACGAGGGCTACAAGTCCCTGTTTAAGAAGGATATCATCGAGACAATCCTGCCA

GAGTTCCTGGACGATAAGGACGAGATCGCCCTGGTGAACAGCTTCAATGGCTTTACCACAGCCTTCACCG
GCTTCTTTGATAACAGAGAGAATATGTTTTCCGAGGAGGCCAAGAGCACATCCATCGCCTTCAGGTGTAT
CAACGAGAATCTGACCCGCTACATCTCTAATATGGACATCTTCGAGAAGGTGGACGCCATCTTTGATAAG
CACGAGGTGCAGGAGATCAAGGAGAAGATCCTGAACAGCGACTATGATGTGGAGGATTTCTTTGAGGGCG
AGTTCTTTAACTTTGTGCTGACACAGGAGGGCATCGACGTGTATAACGCCATCATCGGCGGCTTCGTGAC
CGAGAGCGGCGAGAAGATCAAGGGCCTGAACGAGTACATCAACCTGTATAATCAGAAAACCAAGCAGAAG
CTGCCTAAGTTTAAGCCACTGTATAAGCAGGTGCTGAGCGATCGGGAGTCTCTGAGCTTCTACGGCGAGG
GCTATACATCCGATGAGGAGGTGCTGGAGGTGTTTAGAAACACCCTGAACAAGAACAGCGAGATCTTCAG
CTCCATCAAGAAGCTGGAGAAGCTGTTCAAGAATTTTGACGAGTACTCTAGCGCCGGCATCTTTGTGAAG
AACGGCCCCGCCATCAGCACAATCTCCAAGGATATCTTCGGCGAGTGGAACGTGATCCGGGACAAGTGGA
ATGCCGAGTATGACGATATCCACCTGAAGAAGAAGGCCGTGGTGACCGAGAAGTACGAGGACGATCGGAG
AAAGTCCTTCAAGAAGATCGGCTCCTTTTCTCTGGAGCAGCTGCAGGAGTACGCCGACGCCGATCTGTCT
GTGGTGGAGAAGCTGAAGGAGATCATCATCCAGAAGGTGGATGAGATCTACAAGGTGTATGGCTCCTCTG
AGAAGCTGTTCGACGCCGATTTTGTGCTGGAGAAGAGCCTGAAGAAGAACGACGCCGTGGTGGCCATCAT
GAAGGACCTGCTGGATTCTGTGAAGAGCTTCGAGAATTACATCAAGGCCTTCTTTGGCGAGGGCAAGGAG
ACAAACAGGGACGAGTCCTTCTATGGCGATTTTGTGCTGGCCTACGACATCCTGCTGAAGGTGGACCACA
TCTACGATGCCATCCGCAATTATGTGACCCAGAAGCCCTACTCTAAGGATAAGTTCAAGCTGTATTTTCA
GAACCCTCAGTTCATGGGCGGCTGGGACAAGGATAAGGAGACAGACTATCGGGCCACCATCCTGAGATAC
GGCTCCAAGTACTATCTGGCCATCATGGATAAGAAGTACGCCAAGTGCCTGCAGAAGATCGACAAGGACG
ATGTGAACGGCAATTACGAGAAGATCAACTATAAGCTGCTGCCCGGCCCTAATAAGATGCTGCCAAAGGT
GTTCTTTTCTAAGAAGTGGATGGCCTACTATAACCCCAGCGAGGACATCCAGAAGATCTACAAGAATGGC
ACATTCAAGAAGGGCGATATGTTTAACCTGAATGACTGTCACAAGCTGATCGACTTCTTTAAGGATAGCA
TCTCCCGGTATCCAAAGTGGTCCAATGCCTACGATTTCAACTTTTCTGAGACAGAGAAGTATAAGGACAT
CGCCGGCTTTTACAGAGAGGTGGAGGAGCAGGGCTATAAGGTGAGCTTCGAGTCTGCCAGCAAGAAGGAG
GTGGATAAGCTGGTGGAGGAGGGCAAGCTGTATATGTTCCAGATCTATAACAAGGACTTTTCCGATAAGT
CTCACGGCACACCCAATCTGCACACCATGTACTTCAAGCTGCTGTTTGACGAGAACAATCACGGACAGAT
CAGGCTGAGCGGAGGAGCAGAGCTGTTCATGAGGCGCGCCTCCCTGAAGAAGGAGGAGCTGGTGGTGCAC
CCAGCCAACTCCCCTATCGCCAACAAGAATCCAGATAATCCCAAGAAAACCACAACCCTGTCCTACGACG
TGTATAAGGATAAGAGGTTTTCTGAGGACCAGTACGAGCTGCACATCCCAATCGCCATCAATAAGTGCCC
CAAGAACATCTTCAAGATCAATACAGAGGTGCGCGTGCTGCTGAAGCACGACGATAACCCCTATGTGATC
GGCATCGCCAGGGGCGAGCGCAATCTGCTGTATATCGTGGTGGTGGACGGCAAGGGCAACATCGTGGAGC
AGTATTCCCTGAACGAGATCATCAACAACTTCAACGGCATCAGGATCAAGACAGATTACCACTCTCTGCT
GGACAAGAAGGAGAAGGAGAGGTTCGAGGCCCGCCAGAACTGGACCTCCATCGAGAATATCAAGGAGCTG
AAGGCCGGCTATATCTCTCAGGTGGTGCACAAGATCTGCGAGCTGGTGGAGAAGTACGATGCCGTGATCG
CCCTGGAGGACCTGAACTCTGGCTTTAAGAATAGCCGCGTGAAGGTGGAGAAGCAGGTGTATCAGAAGTT
CGAGAAGATGCTGATCGATAAGCTGAACTACATGGTGGACAAGAAGTCTAATCCTTGTGCAACAGGCGGC
GCCCTGAAGGGCTATCAGATCACCAATAAGTTCGAGAGCTTTAAGTCCATGTCTACCCAGAACGGCTTCA
TCTTTTACATCCCTGCCTGGCTGACATCCAAGATCGATCCATCTACCGGCTTTGTGAACCTGCTGAAAAC
CAAGTATACCAGCATCGCCGATTCCAAGAAGTTCATCAGCTCCTTTGACAGGATCATGTACGTGCCCGAG
GAGGATCTGTTCGAGTTTGCCCTGGACTATAAGAACTTCTCTCGCACAGACGCCGATTACATCAAGAAGT
GGAAGCTGTACTCCTACGGCAACCGGATCAGAATCTTCCGGAATCCTAAGAAGAACAACGTGTTCGACTG
GGAGGAGGTGTGCCTGACCAGCGCCTATAAGGAGCTGTTCAACAAGTACGGCATCAATTATCAGCAGGGC
GATATCAGAGCCCTGCTGTGCGAGCAGTCCGACAAGGCCTTCTACTCTAGCTTTATGGCCCTGATGAGCC
TGATGCTGCAGATGCGGAACAGCATCACAGGCCGCACCGACGTGGATTTTCTGATCAGCCCTGTGAAGAA
CTCCGACGGCATCTTCTACGATAGCCGGAACTATGAGGCCCAGGAGAATGCCATCCTGCCAAAGAACGCC

GACGCCAATGGCGCCTATAACATCGCCAGAAAGGTGCTGTGGGCCATCGGCCAGTTCAAGAAGGCCGAGG
ACGAGAAGCTGGATAAGGTGAAGATCGCCATCTCTAACAAGGAGTGGCTGGAGTACGCCCAGACCAGCGT
GAAGCACTGCGCGGGTGGCGGAGGAAGTGGGGGCGGTGGTAGCATGAATAAAAACATTGATACAGTTCGT
GAAATTATTACTGTTGCATCTATTTTGATTAAATTTTCCAGAGAAGATATTGTTGAGAATCGTGCTAATT
TTATTGCATTTCTAAATGAGATTGGAGTAACGCATGAAGGTAGAAAGTTAAATCGCAATTCATTCCGTAA
AATTATTTCTAAATTAACTCAAGAAGATAAGAAAACCCTCATCGACGAATTCAACGAGGGTTTTGAGGGT
GTATATCGATATCTAGAGATGTATACGAACAAA**TAA**CTCGAGTAAGGATCTCCAGGCATCAAATAAAACG
AAAGGCTCAGTCGAAAGACTGGGCCTTTCGTTTTATCTGTTGTTTGTCGGTGAACGCTCTCTACTAGAGT
CACACTGGCTCACCTTCGGGTGGGCCTTTCTGCGTTTATA

>BBa_J23107-RBS-MCP-SoxS-BBa_B1002_
BBa_1002-λN22-PspF-RBS-BBa_J23107
TTTACGGCTAGCTCAGCCCTAGGTATTATGCTAGCGAATTCATTAAAGAGGAGAAAGGTACC**ATG**GGGCC
CGCTTCTAACTTTACTCAGTTCGTTCTCGTCGACAATGGCGGAACTGGCGACGTGACTGTCGCCCCAAGC
AACTTCGCTAACGGGATCGCTGAATGGATCAGCTCTAACTCGCGTTCACAGGCTTACAAAGTAACCTGTA
GCGTTCGTCAGAGCTCTGCGCAGAATCGCAAATACACCATCAAAGTCGAGGTGCCTAAAGGCGCCTGGCG
TTCGTACTTAAATATGGAACTAACCATTCCAATTTTCGCCACGAATTCCGACTGCGAGCTTATTGTTAAG
GCAATGCAAGGTCTCCTAAAAGATGGAAACCCGATTCCCTCAGCAATCGCAGCAAACTCCGGCATCTACG
GTGGCGGAGGTAGCATGTCCCATCAGAAAATTATTCAGGATCTTATCGCATGGATTGACGAGCATATTGA
CCAGCCGCTTAACATTGATGTAGTCGCAAAAAAATCAGGCTATTCAAAGTGGTACTTGCAACGAATGTTC
CGCACGGTGACGCATCAGACGCTTGGCGATTACATTCGCCAACGCCGCCTGTTACTGGCCGCCGTTGAGT
TGCGCACCACCGAGCGTCCGATTTTTGATATCGCAATGGACCTGGGTTATGTCTCGCAGCAGACCTTCTC
CCGCGTTTTCGCGCGGCAGTTTGATCGCACTCCCGCGGATTATCGCCACCGCCTG**TAA**GCGGCCGCCACG
CAAAAAACCCCGCTTCGGCGGGGTTTTTTCGCCCTAGGGATATATTCCGCTTCCTCGCTCACTGACTCGC
TACGCTCGGTCGTTCGACTGCGGCGAGCGGAAATGGCTTACGAACGGGGCGGAGATTTCCTGCCCGGGGC
GAAAAAACCCCGCCGAAGCGGGGTTTTTTGCGTGGGCGCGCC**TTA**ATTTGCTGCTTTCCATTGAGCCTGT
TTCTCTGCGCGACGTTCGCGGCGGCGTGTGCGTGCATTCATTTGCAAACTGAGTTGCAGCAACTCTTTTT
CCTGCTGCATCTGAAACTCACGTAAATCCAGCGGCAGTGTTGGAAGCGAGGTGGTTTCTGAAACGGCGAT
AGCGTCTTCAGGCGGACGCCGTTTAAAGGGATCAATAATGATGTCATCAAGCGGATAATCGCTGGTGCCG
TGGCGATACACTGAACGTTCCACCACGTTTTTCAATTCACGAATATTTCCCGGCCAACGATAATTCAGCA
ATGTTTCTCTGGCGCGCTCCGTAAACCCCGGGAACAGAGGCAGCTTGATTTCCCGACACATCTGGATGGC
AAAGTATTCTGCCATCAACATTATGTCGCTTTCGCGCTCGCGCAGTGGTGGCAGTTGTACAACATCAAAA
GCCAGTCGGTCGAGCAGGTCAGCGCGAAAAGTGCCTTCATTGACCATCGCCGGGAGATCGGCATTCGTCG
CGCATACCAACCGCACATTCACCTGCAATGGTTGGCTGCCGCCAACGCGCTCCAGTTCACCGTACTCAAT
CACGCGCAATAATTTCTCCTGCACCATCATCGGTGCCGTAGCGAGTTCATCAAGAAATAGCGTACCGCCG
TCGGCACGTTCAAATCTCCCTGGATGACGTTTTTGCGCACCGGTAAACGCCCCCGCTTCGTGACCAAACA
GTTCGGAATCCAGCAGATTTTCATTTAACGCCGCGCAGTTAAGGGAAATAAACGGCCCTTGCCAACGGGA
GGAGAGATAATGCAGGCGGCTGGCAATCAGCTCTTTACCGGTGCCGCGTTCGCCGATGATGAGCACCGGT
TTGTCCAGCGGTGCGAGATGCGAAACCTGTTCCAGCACTTCGAGAAAGCTGTTCGCCTCACCAAGTAAAT
TATCTTTGTATTCTGCCATGGGCCC**CAT**GGTACCTTTCTCCTCTTTAATGAATTCGCTAGCATAATACCT
AGGGCTGAGCTAGCCGTAAA

>TetR-Ptet-RBS-MCP-SoxS-BBa_B1002_
BBa_1002-λN22-PspF-RBS-PrhaB-RhaS

TTAAGACCCACTTTCACATTTAAGTTGTTTTTCTAATCCGCATATGATCAATTCAAGGCCGAATAAGAAG
GCTGGCTCTGCACCTTGGTGATCAAATAATTCGATAGCTTGTCGTAATAATGGCGGCATACTATCAGTAG
TAGGTGTTTCCCTTTCTTCTTTAGCGACTTGATGCTCTTGATCTTCCAATACGCAACCTAAAGTAAAATG
CCCCACAGCGCTGAGTGCATATAATGCATTCTCTAGTGAAAAACCTTGTTGGCATAAAAAGGCTAATTGA
TTTTCGAGAGTTTCATACTGTTTTTCTGTAGGCCGTGTACCTAAATGTACTTTTGCTCCATCGCGATGAC
TTAGTAAAGCACATCTAAAACTTTTAGCGTTATTACGTAAAAAATCTTGCCAGCTTTCCCCTTCTAAAGG
GCAAAAGTGAGTATGGTGCCTATCTAACATCTCAATGGCTAAGGCGTCGAGCAAAGCCCGCTTATTTTTT
ACATGCCAATACAATGTAGGCTGCTCTACACCTAGCTTCTGGGCGAGTTTACGGGTTGTTAAACCTTCGA
TTCCGACCTCATTAAGCAGCTCTAATGCGCTGTTAATCACTTTACTTTTATCTAATCTAGACATCATTAA
TTCCTAATTTTTGTTGACACTCTATCGTTGATAGAGTTATTTTACCACTCCCTATCAGTGATAGAGAAAA
GAATTCATTAAAGAGGAGAAAGGTACC**ATG**GGCCCGCTTCTAACTTTACTCAGTTCGTTCTCGTCGACA
ATGGCGGAACTGGCGACGTGACTGTCGCCCCAAGCAACTTCGCTAACGGGATCGCTGAATGGATCAGCTC
TAACTCGCGTTCACAGGCTTACAAAGTAACCTGTAGCGTTCGTCAGAGCTCTGCGCAGAATCGCAAATAC
ACCATCAAAGTCGAGGTGCCTAAAGGCGCCTGGCGTTCGTACTTAAATATGGAACTAACCATTCCAATTT
TCGCCACGAATTCCGACTGCGAGCTTATTGTTAAGGCAATGCAAGGTCTCCTAAAAGATGGAAACCCGAT
TCCCTCAGCAATCGCAGCAAACTCCGGCATCTACGGTGGCGGAGGTAGCATGTCCCATCAGAAAATTATT
CAGGATCTTATCGCATGGATTGACGAGCATATTGACCAGCCGCTTAACATTGATGTAGTCGCAAAAAAAT
CAGGCTATTCAAAGTGGTACTTGCAACGAATGTTCCGCACGGTGACGCATCAGACGCTTGGCGATTACAT
TCGCCAACGCCGCCTGTTACTGGCCGCCGTTGAGTTGCGCACCACCGAGCGTCCGATTTTTGATATCGCA
ATGGACCTGGGTTATGTCTCGCAGCAGACCTTCTCCCGCGTTTTCGCGCGGCAGTTTGATCGCACTCCCG
CGGATTATCGCCACCGCCTG**TAA**GCGGCCGCCACGCAAAAAACCCCGCTTCGGCGGGGTTTTTTCGCCCT
AGGGATATATTCCGCTTCCTCGCTCACTGACTCGCTACGCTCGGTCGTTCGACTGCGGCGAGCGGAAATG
GCTTACGAACGGGGCGGAGATTTCCTGCCCGGGGCGAAAAACCCCGCCGAAGCGGGGTTTTTTGCGTGG
GCGCGCC**TTA**ATTTGCTGCTTTCCATTGAGCCTGTTTCTCTGCGCGACGTTCGCGGCGGCGTGTGCGTGC
ATTCATTTGCAAACTGAGTTGCAGCAACTCTTTTTCCTGCTGCATCTGAAACTCACGTAAATCCAGCGGC
AGTGTTGGAAGCGAGGTGGTTTCTGAAACGGCGATAGCGTCTTCAGGCGGACGCCGTTTAAAGGGATCAA
TAATGATGTCATCAAGCGGATAATCGCTGGTGCCGTGGCGATACACTGAACGTTCCACCACGTTTTTCAA
TTCACGAATATTTCCCGGCCAACGATAATTCAGCAATGTTTCTCTGGCGCGCTCCGTAAACCCCGGGAAC
AGAGGCAGCTTGATTTCCCGACACATCTGGATGGCAAAGTATTCTGCCATCAACATTATGTCGCTTTCGC
GCTCGCGCAGTGGTGGCAGTTGTACAACATCAAAAGCCAGTCGGTCGAGCAGGTCAGCGCGAAAAGTGCC
TTCATTGACCATCGCCGGGAGATCGGCATTCGTCGCGCATACCAACCGCACATTCACCTGCAATGGTTGG
CTGCCGCCAACGCGCTCCAGTTCACCGTACTCAATCACGCGCAATAATTTCTCCTGCACCATCATCGGTG
CCGTAGCGAGTTCATCAAGAAATAGCGTACCGCCGTCGGCACGTTCAAATCTCCCTGGATGACGTTTTTG
CGCACCGGTAAACGCCCCCGCTTCGTGACCAAACAGTTCGGAATCCAGCAGATTTTCATTTAACGCCGCG
CAGTTAAGGGAAATAAACGGCCCTTGCCAACGGGAGGAGAGATAATGCAGGCGGCTGGCAATCAGCTCTT
TACCGGTGCCGCGTTCGCCGATGATGAGCACCGGTTTGTCCAGCGGTGCGAGATGCGAAACCTGTTCCAG
CACTTCGAGAAAGCTGTTCGCCTCACCAAGTAAATTATCTTTGTATTCTGCCATGGGCCC**CAT**GGTACCT
TTCTCCTCTTTAATGAATTCTTACGACCAGTCTAAAAAGCGCCTCAATTCGCGACCTTCTCGTTACTGAC
AGGAAAATGGGCCATTGGCAACCAGGGAAAGATGAACGTGATGATGTTCACAATTTGCTGAATTGTGGTG
ATGTGATGCTCACCGCATTCCTGCTCCAGCGGCGTAAAGGTCTCTCTAGTATATAAACGCAGAAAGGCCC
ACCCGAAGGTGAGCCAGTGTGATACTAGAGTTTACAGCTAGCTCAGTCCTAGGTATTATGCTAGCTACTA
GAGGGAGCCCAAATGACCGTATTACATAGTGTGGATTTTTTTCCGTCCGGTAACGCGTCCGTGGCGATAG
AACCCCGGCTCCCGCAGGCGGATTTTCCTGAACATCATCATGATTTTCATGAAATTGTGATTGTCGAACA
TGGCACGGGTATTCATGTGTTTAATGGGCAGCCCTATACCATCACCGGTGGCACGGTCTGTTTCGTACGC
GATCATGATCGGCATCTGTATGAACATACCGATAATCTGTGTCTGACCAATGTGCTGTATCGCTCGCCGG

ATCGATTTCAGTTTCTCGCCGGGCTGAATCAGTTGCTGCCACAAGAGCTGGATGGGCAGTATCCGTCTCA
CTGGCGCGTTAACCACAGCGTATTGCAGCAGGTGCGACAGCTGGTTGCACAGATGGAACAGCAGGAAGGG
GAAAATGATTTACCCTCGACCGCCAGTCGCGAGATCTTGTTTATGCAATTACTGCTCTTGCTGCGTAAAA
GCAGTTTGCAGGAGAACCTGGAAAACAGCGCATCACGTCTCAACTTGCTTCTGGCCTGGCTGGAGGACCA
TTTTGCCGATGAGGTGAATTGGGATGCCGTGGCGGATCAATTTTCTCTTTCACTGCGTACGCTACATCGG
CAGCTTAAGCAGCAAACGGGACTGACGCCTCAGCGATACCTGAACCGCCTGCGACTGATGAAAGCCCGAC
ATCTGCTACGCCACAGCGAGGCCAGCGTTACTGACATCGCCTATCGCTGTGGATTCAGCGACAGTAACCA
CTTTTCGACGCTTTTTCGCCGAGAGTTTAACTGGTCACCGCGTGATATTCGCCAGGGACGGGATGGCTTT
CTGCAATAATACTAGCTCGGTACCAAAGACGAACAATAAGACGCTGAAAAGCGTCTTTTTTCGTTTTGGT
CCCTCTAGTATTAATTTTGTCTTCATGCATGAAGACAACTAGG

Systematic Reporters

Minimal ~35 bp promoters were **bolded**.
-24 and -12 elements for the $\sigma^{54}$-promoter were underlined.
The full ORF of mRFP after ATG start codon was omitted to reduce redundancy.

>J1-**BBa_J23117**-RBS-mRFP-dblT
GCCTACGGTATCCACCGGAGACCTATGGCAGCCTCCGGCCGCCATAGGACACCTTTGGTTGCCAAGGGTG
ACCTATGGTGACCATGGGCCACCACGGGCGACCTCAGGTATCCTGCGGTGTCCTGCGGTTACCAAAGGCG
TCCTTTGGGTTCCACCGGATACCTCCGGAC**TTGACAGCTAGCTCAGTCCTAGGGATTGTGCTAGC**GAATT
CATTAAAGAGGAGAAAGGTACC**ATG**GCGAGTAGCGAAGACGTTATCAAAGAGTTCATGCGTTTCAAAGTT
CGTATGGAAGGTTCCGTTAACGGTCACGAGTTCGAAATCGAAGGTGAAGGTGAAGGTCGTCCGTACGAAG
GTACCCAGACCGCTAAACTGAAAGTTACCAAAGGTGGTCCGCTGCCGTTCGCTTGGGACATCCTGTCCCC
GCAGTTCCAGTACGGTTCCAAAGCTTACGTTAAACACCCGGCTGACATCCCGGACTACCTGAAACTGTCC
TTCCCGGAAGGTTTCAAATGGGAACGTGTTATGAACTTCGAAGACGGTGGTGTTGTTACCGTTACCCAGG
ACTCCTCCCTGCAAGACGGTGAGTTCATCTACAAAGTTAAACTGCGTGGTACCAACTTCCCGTCCGACGG
TCCGGTTATGCAGAAAAAAACCATGGGTTGGGAAGCTTCCACCGAACGTATGTACCCGGAAGACGGTGCT
CTGAAAGGTGAAATCAAAATGCGTCTGAAACTGAAAGACGGTGGTCACTACGACGCTGAAGTTAAAACCA
CCTACATGGCTAAAAAACCGGTTCAGCTGCCGGGTGCTTACAAAACCGACATCAAACTGGACATCACCTC
CCACAACGAAGACTACACCATCGTTGAACAGTACGAACGTGCTGAAGGTCGTCACTCCACCGGTGCT**TAA**
GGATCCAAACTCGAGTAAGGATCTCCAGGCATCAAATAAAACGAAAGGCTCAGTCGAAAGACTGGGCCTT
TCGTTTTATCTGTTGTTTGTCGGTGAACGCTCTCTACTAGAGTCACACTGGCTCACCTTCGGGTGGGCCT
TTCTGCGTTTATA

>J1-**sodCp**-RBS-mRFP-dblT
GCCTACGGTATCCACCGGAGACCTATGGCAGCCTCCGGCCGCCATAGGACACCTTTGGTTGCCAAGGGTG
ACCTATGGTGACCATGGGCCACCACGGGCGACCTCAGGTATCCTGCGGTGTCCTGCGGTTACCAAAGGCG
TCCTTTGGGTTCCACCGGATACCTCCGGACGCCGTTCAAAAATGTGTCACTGGTTTACACTTATTCAGGG
AATTCATTAAAGAGGAGAAAGGTACC**ATG**___

>J1(-3)-**sodCp**-RBS-mRFP-dblT
GCCTACGGTATCCACCGGAGACCTATGGCAGCCTCCGGCCGCCATAGGACACCTTTGGTTGCCAAGGGTG
ACCTATGGTGACCATGGGCCACCACGGGCGACCTCAGGTATCCTGCGGTGTCCTGCGGTTACCAAAGGCG
TCCTTTGGGTTCCACCGGATACCTCCGGCCGTTCAAAAATGTGTCACTGGTTTACACTTATTCAGGGAAT
TCATTAAAGAGGAGAAAGGTACC**ATG**___

>J1-**rdbBp**-RBS-mRFP-dblT
GCCTACGGTATCCACCGGAGACCTATGGCAGCCTCCGGCCGCCATAGGACACCTTTGGTTGCCAAGGGTG
ACCTATGGTGACCATGGGCCACCACGGGCGACCTCAGGTATCCTGCGGTGTCCTGCGGTTACCAAAGGCG
TCCTTTGGGTTCCACCGGATACCTCCGGAC**TTGTAAAAGGCGAACTTGGCCGCCACAAACAAATT**GAATT
CATTAAAGAGGAGAAAGGTACC**ATG**___

>J1-yieEp-RBS-mRFP-dblT
GCCTACGGTATCCACCGGAGACCTATGGCAGCCTCCGGCCGCCATAGGACACCTTTGGTTGCCAAGGGTG
ACCTATGGTGACCATGGGCCACCACGGGCGACCTCAGGTATCCTGCGGTGTCCTGCGGTTACCAAAGGCG
TCCTTTGGGTTCCACCGGATACCTCCGGAC**GAACTTTTAGCCGCTTTAGTCTGTCCATCATTCCA**GAATT
CATTAAAGAGGAGAAAGGTACC**ATG**___

>J1-glnAp2-RBS-mRFP-dblT
GCCTACGGTATCCACCGGAGACCTATGGCAGCCTCCGGCCGCCATAGGACACCTTTGGTTGCCAAGGGTG
ACCTATGGTGACCATGGGCCACCACGGGCGACCTCAGGTATCCTGCGGTGTCCTGCGGTTACCAAAGGCG
TCCTTTGGGTTCCACCGGATACCTCCGGAC**TGGCACAGATTTCGCTTTATCTTTTTTACGGCGAC**GAATT
CATTAAAGAGGAGAAAGGTACC**ATG**___

>J1-glnAp2(shifted)-RBS-mRFP-dblT
GCCTACGGTATCCACCGGAGACCTATGGCAGCCTCCGGCCGCCATAGGACACCTTTGGTTGCCAAGGGTG
ACCTATGGTGACCATGGGCCACCACGGGCGACCTCAGGTATCCTGCGGTGTCCTGCGGTTACCAAAGGCG
TCCTTTGGGTTCCACCGGATACCTCCGGAC**TTTAAAAGTTGGCACAGATTTCGCTTTATCTTTTT**tGAAT
TCATTAAAGAGGAGAAAGGTACC**ATG**___

>J1-pspAp-RBS-mRFP-dblT
GCCTACGGTATCCACCGGAGACCTATGGCAGCCTCCGGCCGCCATAGGACACCTTTGGTTGCCAAGGGTG
ACCTATGGTGACCATGGGCCACCACGGGCGACCTCAGGTATCCTGCGGTGTCCTGCGGTTACCAAAGGCG
TCCTTTGGGTTCCACCGGATACCTCCGGAC**ATAAAAAATTGGCACGCAAATTGTATTAACAGTTCA**GAAT
TCATTAAAGAGGAGAAAGGTACC**ATG**___

>J1(-1)-pspAp-RBS-mRFP-dblT
GCCTACGGTATCCACCGGAGACCTATGGCAGCCTCCGGCCGCCATAGGACACCTTTGGTTGCCAAGGGTG
ACCTATGGTGACCATGGGCCACCACGGGCGACCTCAGGTATCCTGCGGTGTCCTGCGGTTACCAAAGGCG
TCCTTTGGGTTCCACCGGATACCTCGGAC**ATAAAAAATTGGCACGCAAATTGTATTAACAGTTCA**GAATT
CATTAAAGAGGAGAAAGGTACC**ATG**___

>W1-BBa_J23117-RBS-mRFP-dblT
GCATGCCCGATCAACGTCTCATTTTCGCCAGATATCAAGCAGAGGAGCAAAAGCTCATTTCTGAAGAGGA
CTTGTTGCGGAAACGACGAGAACAGTTGAAACACAAACTTGAACAGCTACGGAACTCTTGTGCGTAAGGA
AAAGTAAGGAAAACGATTCCTTCTAACAGAAATGTCCTGAGCAATCACCTATGAACTGTCGACTCGAGCC
TCTATGGATTATCACCTTGGCTGCAGGCCGGATCTTCCACAACACGCACGGTGTTACATTAGGCATACCG
GTC**TTGACAGCTAGCTCAGTCCTAGGGATTGTGCTAGC**GAATTCATTAAAGAGGAGAAAGGTACC**ATG**__
_

>J3-BBa_J23117-RBS-mRFP-dblT
AGCATTTGCGATCATTCACGCAGCGCTTATTCAGTTGCTCACTGCGATGTCATAATCATCGCTACGAGCT
GTGAAAGATGCATAAAGCTCGTACGACGCGTTCGCTCGTCTCCTCACTTCTCCTACGGAGCGTTCTGGAC

ACAACGTCGTCTTGAAGTTGCGATTATAGA**TTGACAGCTAGCTCAGTCCTAGGGATTGTGCTAGC**GAATT
CATTAAAGAGGAGAAAGGTACC**ATG**___

>J1-J3(100)-**BBa_J23117**-RBS-mRFP-dblT
GCCTACGGTATCCACCGGAGACCTATGGCAGCCTCCGGCCGCCATAGGACACCTTTGGTTGCCAAGGGTG
ACCTATGGTGACCATGGGCCACCACGGGCGACCTCAGGTATCCTGCGGTGTCCTGCGGTTACCAAAGGCG
TCCTTTGGGTTCCACCGGATACCTCCGGACGTGAAAGATGCATAAAGCTCGTACGACGCGTTCGCTCGTC
TCCTCACTTCTCCTACGGAGCGTTCTGGACACAACGTCGTCTTGAAGTTGCGATTATAGA**TTGACAGCTA
GCTCAGTCCTAGGGATTGTGCTAGC**GAATTCATTAAAGAGGAGAAAGGTACC**ATG**___

>J1-J2(100)-J3(100)-**BBa_J23117**-RBS-mRFP-dblT
GCCTACGGTATCCACCGGAGACCTATGGCAGCCTCCGGCCGCCATAGGACACCTTTGGTTGCCAAGGGTG
ACCTATGGTGACCATGGGCCACCACGGGCGACCTCAGGTATCCTGCGGTGTCCTGCGGTTACCAAAGGCG
TCCTTTGGGTTCCACCGGATACCTCCGGACCTACTTGAGTGAGCACTGAAAGCGCGCTGTCGCGCGACTG
ACCTGACGCATCCTGAGGACGTGTTCGGCTACTACACAAGTATTAAGAGACAATGCGCTCGTGAAAGATG
CATAAAGCTCGTACGACGCGTTCGCTCGTCTCCTCACTTCTCCTACGGAGCGTTCTGGACACAACGTCGT
CTTGAAGTTGCGATTATAGA**TTGACAGCTAGCTCAGTCCTAGGGATTGTGCTAGC**GAATTCATTAAAGAG
GAGAAAGGTACC**ATG**___

>J1-J3(122)-**BBa_J23117**-RBS-mRFP-dblT
GCCTACGGTATCCACCGGAGACCTATGGCAGCCTCCGGCCGCCATAGGACACCTTTGGTTGCCAAGGGTG
ACCTATGGTGACCATGGGCCACCACGGGCGACCTCAGGTATCCTGCGGTGTCCTGCGGTTACCAAAGGCG
TCCTTTGGGTTCCACCGGATACCTCCGGACGTCATAATCATCGCTACGAGCTGTGAAAGATGCATAAAGC
TCGTACGACGCGTTCGCTCGTCTCCTCACTTCTCCTACGGAGCGTTCTGGACACAACGTCGTCTTGAAGT
TGCGATTATAGA**TTGACAGCTAGCTCAGTCCTAGGGATTGTGCTAGC**GAATTCATTAAAGAGGAGAAAGG
TACC**ATG**___

>K1T-J3(112)-**BBa_J23117**-RBS-mRFP-dblT
TTCATGGAGAAAGACATCCAAGTCTTGAGAATGCCTCCCACCGAGCGAGAATAGACTTCGACTGCCCAGA
AACGACCGGCAGACGTGAGAAAGGTGATAGTGCGAAGAGAATGCGGACCACTTGACCAGAATAAAGTCTC
GCAGATGAGAATGGGTTCACAGCGACAAGAAACGACCTGCTCGCTACGAGCTGTGAAAGATGCATAAAGC
TCGTACGACGCGTTCGCTCGTCTCCTCACTTCTCCTACGGAGCGTTCTGGACACAACGTCGTCTTGAAGT
TGCGATTATAGA**TTGACAGCTAGCTCAGTCCTAGGGATTGTGCTAGC**GAATTCATTAAAGAGGAGAAAGG
TACC**ATG**___

>K1B_+10-J3(112)-**BBa_J23117**-RBS-mRFP-dblT
GCAGGTCGTTTCTTGTCGCTGTGAACCCATTCTCATCTGCGAGACTTTATTCTGGTCAAGTGGTCCGCAT
TCTCTTCGCACTATCACCTTTCTCACGTCTGCCGGTCGTTTCTGGGCAGTCGAAGTCTATTCTCGCTCGG
TGGGAGGCATTCTCAAGACTTGGATGTCTTTCTCCATGAAAGATGCGCTCTCGCTACGAGCTGTGAAAGA
TGCATAAAGCTCGTACGACGCGTTCGCTCGTCTCCTCACTTCTCCTACGGAGCGTTCTGGACACAACGTC
GTCTTGAAGTTGCGATTATAGA**TTGACAGCTAGCTCAGTCCTAGGGATTGTGCTAGC**GAATTCATTAAAG
AGGAGAAAGGTACC**ATG**___

>I1T_+15-J3(112)-**BBa_J23117**-RBS-mRFP-dblT

ATTTCGCTGACACCACAGTCCTTTCGCTTTGAAGTCGGTACTTTACTGCCAAGCCTGGGCCTTTATCGCT
ATGCGGACGGCTTTATCTGCGGCCAGCACCATTTCGTGGCACAACTATAGGTTTAATATGGCTCCCGGTC
ATTTCCGTCGAATCATCGGGCTTTCAGTTCGCGATAGATGCGCTCTCGCTACGAGCTGTGAAAGATGCAT
AAAGCTCGTACGACGCGTTCGCTCGTCTCCTCACTTCTCCTACGGAGCGTTCTGGACACAACGTCGTCTT
GAAGTTGCGATTATAGA**TTGACAGCTAGCTCAGTCCTAGGGATTGTGCTAGC**GAATTCATTAAAGAGGAG
AAAGGTACC**ATG**___

>I1B-J3(112)-**BBa_J23117**-RBS-mRFP-dblT
GAACTGAAAGCCCGATGATTCGACGGAAATGACCGGGAGCCATATTAAACCTATAGTTGTGCCACGAAAT
GGTGCTGGCCGCAGATAAAGCCGTCCGCATAGCGATAAAGGCCCAGGCTTGGCAGTAAAGTACCGACTTC
AAAGCGAAAGGACTGTGGTGTCAGCGAAATTCGCTACGAGCTGTGAAAGATGCATAAAGCTCGTACGACG
CGTTCGCTCGTCTCCTCACTTCTCCTACGGAGCGTTCTGGACACAACGTCGTCTTGAAGTTGCGATTATA
GA**TTGACAGCTAGCTCAGTCCTAGGGATTGTGCTAGC**GAATTCATTAAAGAGGAGAAAGGTACC**ATG**___

>J4-J1-**pspAp**-RBS-mRFP-dblT
TCGAGAGAAGTCACTCTTGACGACACAACTGCGTGTCACGTAAAGCGAGATTGCGTAAGCGTTAGCAGAG
TCAAAGTTACATCGCAGTTGACCACGGGCGACCTCAGGTATCCTGCGGTGTCCTGCGGTTACCAAAGGCG
TCCTTTGGGTTCCACCGGATACCTCCGGAC**ATAAAAAAT<u>TGGCAC</u>GCAAAT<u>TGTAT</u>TAACAGTTCA**GAAT
TCATTAAAGAGGAGAAAGGTACC**ATG**___

>J4(J110/J102)-**pspAp**-RBS-mRFP-dblT
TCGAGAGAAGTCACTCTTGACGACACAACTGCGTGTCACGTAAAGCGAGATTGCGTAAGCGTTAGCAGAG
TCAAAGTTACATCGCAGTTGCGCTGTTCGACCCTCAGGTATCCTGCGGTGTCCTCCGGAAGTGAAAGGAT
GCCTTTGGGTTCCACCGGATACCTCCGGAC**ATAAAAAAT<u>TGGCAC</u>GCAAAT<u>TGTAT</u>TAACAGTTCA**GAAT
TCATTAAAGAGGAGAAAGGTACC**ATG**___

sgRNA/scRNA/crRNA

20 bp spacers are underlined.
Full rrnB terminator was omitted to reduce redundancy.

>BBa_J23119(SpeI)-sgRNA-TrrnB
TTGACAGCTAGCTCAGTCCTAGGTATAATACTAGTNNNNNNNNNNNNNNNNNNNNGTTTTAGAGCTAGAA
ATAGCAAGTTAAAATAAGGCTAGTCCGTTATCAACTTGAAAAAGTGGCACCGAGTCGGTGCTTTTTTTGA
AGCTTGGGCCCGAACAAAAACTCATCTCAGAAGAGGATCTGAATAGCGCCGTCGACCATCATCATCATCA
TCATTGAGTTTAAACGGTCTCCAGCTTGGCTGTTTTGGCGGATGAGAGAAGATTTTCAGCCTGATACAGA
TTAAATCAGAACGCAGAAGCGGTCTGATAAAACAGAATTTGCCTGGCGGCAGTAGCGCGGTGGTCCCACC
TGACCCCATGCCGAACTCAGAAGTGAAACGCCGTAGCGCCGATGGTAGTGTGGGGTCTCCCCATGCGAGA
GTAGGGAACTGCCAGGCATCAAATAAAACGAAAGGCTCAGTCGAAAGACTGGGCCTTTCGTTTTATCTGT
TGTTTGTCGGTGAACT

>BBa_J23119(SpeI)-scRNA.1xMS2.b2-TrrnB
TTGACAGCTAGCTCAGTCCTAGGTATAATACTAGTNNNNNNNNNNNNNNNNNNNNGTTTTAGAGCTAGAA
ATAGCAAGTTAAAATAAGGCTAGTCCGTTATCAACTTGAAAAAGTGGCACATGAGGATCACCCATGTGCT
TTTTTTGAAGCTTGGGCCC___

>BBa_J23119(SpeI)-scRNA.1xPP7.b1-TrrnB
TTGACAGCTAGCTCAGTCCTAGGTATAATACTAGTNNNNNNNNNNNNNNNNNNNNGTTTTAGAGCTAGAA
ATAGCAAGTTAAAATAAGGCTAGTCCGTTATCAACTTGAAAAAGTAACATAAGGAGTTTATATGGAAACC
CTTATGTTTTTTTGAAGCTTGGGCCC___

>BBa_J23119(SpeI)-scRNA.2xBoxB-TrrnB
TTGACAGCTAGCTCAGTCCTAGGTATAATACTAGTNNNNNNNNNNNNNNNNNNNNGTTTGAGAGCTAGGG
CCCTGAAGAAGGGCCCTAGCAAGTTCAAATAAGGCTAGTCCGTTATCAACTTGGGCCCTGAAGAAGGGCC
CAAGTGGCACCGAGTCGGTGCTTTTTTTGAAGCTTGGGCCC___

>BBa_J23119(SpeI)-scRNA.2xMS2-TrrnB
TTGACAGCTAGCTCAGTCCTAGGTATAATACTAGTNNNNNNNNNNNNNNNNNNNNGTTTGAGAGCTAGGC
CAACATGAGGATCACCCATGTCTGCAGGGCCTAGCAAGTTCAAATAAGGCTAGTCCGTTATCAACTTGGC
CAACATGAGGATCACCCATGTCTGCAGGGCCAAGTGGCACCGAGTCGGTGCTTTTTTTGAAGCTTGGGCC
C___

>BBa_J23119(SpeI)-Sth1-sgRNA-TrrnB
TTGACAGCTAGCTCAGTCCTAGGTATAATACTAGTNNNNNNNNNNNNNNNNNNNNGTTTTTGTACTCGAA
AGAAGCTACAAAGATAAGGCTTCATGCCGAAATCAACACCCTGTCATTTTATGGCAGGGTGTTTTTTTTT
GAAGCTTGGGCCC___

>BBa_J23119(SpeI)-Sth1-scRNA.1xMS2.Append-TrrnB
TTGACAGCTAGCTCAGTCCTAGGTATAATACTAGTNNNNNNNNNNNNNNNNNNNNGTTTTTGTACTCGAA
AGAAGCTACAAAGATAAGGCTTCATGCCGAAATCAACACCCTGTCATTTTATGGCAGGGTGTTACGCACA
TGAGGATCACCCATGTGCTTTTTTTGAAGCTTGGGCCC___

>BBa_J23119(SpeI)-Sth1-scRNA.1xMS2.Extend-TrrnB
TTGACAGCTAGCTCAGTCCTAGGTATAATACTAGTNNNNNNNNNNNNNNNNNNNNGTTTTTGTACTCGAA
AGAAGCTACAAAGATAAGGCTTCATGCCGAAATCAACACCCATGAGGATCACCCATGGGTGTTTTTTTTT
GAAGCTTGGGCCC___

>BBa_J23119(SpeI)-Sth1-scRNA.1xMS2.Replace-TrrnB
TTGACAGCTAGCTCAGTCCTAGGTATAATACTAGTNNNNNNNNNNNNNNNNNNNNGTTTTTGTACTCGAA
AGAAGCTACAAAGATAAGGCTTCATGCCGAAATCGCGCACATGAGGATCACCCATGTGCTTTTTTTGAAG
CTTGGGCCC___

>BBa_J23119(SpeI)-dCas12a-crRNA-BBa_K2680405
TTGACAGCTAGCTCAGTCCTAGGTATAATACTAGTTAATTTCTACTAAGTGTAGATNNNNNNNNNNNNNN
NNNNNNAACGCATGAGAAAGCCCCCGGAAGATCACCTTCCGGGGGCTTTTTTATTGCGC
